# Glutamate release is inhibitory and unnecessary for the long-term potentiation of presynaptic function

**DOI:** 10.1101/099515

**Authors:** Zahid Padamsey, Rudi Tong, Nigel Emptage

## Abstract

Long-term potentiation (LTP) and long-term depression (LTD) of transmitter release probability (P_r_) are thought to be triggered by the activation of glutamate receptors. Here, we demonstrate that glutamate release at CA3-CA1 synapses is in fact inhibitory and unnecessary for increases in P_r_. Instead, at active presynaptic terminals, postsynaptic depolarization alone can increase P_r_ by promoting the release of nitric oxide from neuronal dendrites in a manner dependent on L-type voltage-gated Ca^2+^ channels. The release of glutamate, in contrast, decreases P_r_ by activating presynaptic NMDA receptors (NMDAR). Thus, net changes in P_r_ are determined by the combined effect of both LTP-promoting and LTD-promoting processes, that is, by the amount of glutamate release and postsynaptic depolarization that accompany presynaptic activity, respectively. Neither of these processes directly depends on the activation of postsynaptic NMDARs. We further show that presynaptic changes can be captured by a simple learning rule, in which the role of presynaptic plasticity is to ensure that the ability for a presynaptic terminal to release glutamate is matched with its ability to predict postsynaptic spiking.

## Introduction

Learning and memory are thought to require synaptic plasticity, which refers to the capacity for synaptic connections in the brain to change with experience. The most frequently studied forms of synaptic plasticity are long-term potentiation (LTP) and long-term depression (LTD), which involve long-lasting increases and decreases in synaptic transmission. LTP and LTD can be expressed either postsynaptically, as changes in AMPA receptor (AMPAR) number or presynaptically, as changes in glutamate release probability (P_r_)^1-6^. Traditionally, postsynaptic NMDA receptor (NMDAR) activation is believed to be important for both pre- and postsynaptic forms of plasticity^2,7^. Postsynaptic changes, in particular, have been causally and convincingly linked to NMDAR-dependent Ca^2+^ influx which, via the activation of postsynaptic Ca^2+^-sensitive kinases and phosphatases, triggers changes in the number of synaptic AMPARs^7^. The link between NMDAR activation and presynaptic plasticity, however, is less clear. In the case of presynaptic LTP induction, it is traditionally thought that Ca^2+^ influx through postsynaptic NMDARs triggers the synthesis and release of a retrograde signal, most likely nitric oxide (NO), which in turn triggers increases in P_r_^6,8^ (but other forms of presynaptic plasticity exist^9,10^). Although several studies demonstrate that NMDAR blockade impairs presynaptic LTP^11-15^, some groups have found that presynaptic enhancement can be induced in the presence of NMDAR antagonists^16-20^ (but see^15,21^). This form of plasticity relies on L-type voltage-gated Ca^2+^ channels (L-VGCCs)^16-19^, but may still depend on NO signalling^6,22^. These findings suggest that presynaptic plasticity may not require either the activation of postsynaptic NMDARs or NMDAR-dependent NO synthesis.

The role of glutamate signalling in presynaptic plasticity may also differ from its role in postsynaptic plasticity. Glutamate release is inevitably necessary to drive the postsynaptic depolarization required for both pre- and postsynaptic LTP. This function of glutamate, however, is not site-specific, since depolarization triggered by one synapse will spread to another. The necessity for site-specific glutamate release in LTP induction, at least postsynaptically, is instead imposed by postsynaptic NMDARs^2,7^. Consequently, manipulations that enhance postsynaptic NMDAR signalling at the level of the single synapse reliably augment the induction of postsynaptic LTP^7,23-25^. However, that NMDARs may not to be necessary for the induction of presynaptic LTP^15,21^, suggests that the role of synapse-specific glutamate release in presynaptic plasticity may be different. Indeed, a common finding across a number of studies is that high P_r_ synapses are more likely to show presynaptic depression whereas low P_r_ synapses are more likely to show presynaptic potentiation^11,26-29^. Moreover, glutamate release is known to activate presynaptic NMDARs^30^, which can induce presynaptic LTD^31-33^. Thus enhanced glutamate release at a presynaptic terminal, unlike at a dendritic spine, may not necessarily result in enhanced potentiation, but instead promote depression. Several studies have also demonstrated that presynaptic terminals initially releasing little or no glutamate are reliably potentiated following tetanic stimulation^11,13,26-30,34^. How low P_r_ synapses, including those that are putatively silent, can undergo such activity-dependent potentiation raises questions as to the necessity of synapse-specific glutamate release in presynaptic plasticity.

Here we re-examined the mechanisms underlying activity-dependent presynaptic changes at CA3-CA1 hippocampal synapses, with a particular focus on understanding the role of glutamate in presynaptic plasticity. We bidirectionally manipulated glutamatergic signalling during synaptic activity and examined the resulting consequences on P_r_ at single synapses using Ca^2+^ imaging. Remarkably we found that site-specific glutamatergic signalling was unnecessary for the induction of presynaptic LTP. Instead, postsynaptic depolarization alone could increase P_r_ at active presynaptic terminals by promoting the release of NO from neuronal dendrites in a manner dependent on L-VGCC activation. This increase was both Hebbian, in that it required presynaptic activity to precede postsynaptic depolarization, and site-specific, in that it did not spread to inactive terminals. Glutamate release, in contrast, promoted decreases in P_r_ by activating presynaptic NMDARs. Our findings support a simple learning rule [ΔP_r_=η(P_depol_-P_glu_)], in which net changes in P_r_ (ΔP_r_) at a presynaptic terminal depend on the probability of glutamate release (P_glu_) and the probability of strong postsynaptic depolarization (P_depol_) that accompanies presynaptic activity. The learning rule incorporates both LTP- and LTD- promoting processes into a unified framework, and is capable of predicting changes in P_r_ across a number of experimental conditions.

## Results

### High frequency presynaptic activity inhibits presynaptic LTP and promotes presynaptic LTD

We were interested in understanding the mechanisms underlying activity-dependent presynaptic changes at CA3-CA1 synapses, with a particular focus on understanding the role glutamate plays in presynaptic plasticity. We started by examining how manipulating glutamatergic signalling at synapses would affect activity-driven changes in presynaptic function. We recorded excitatory postsynaptic potentials (EPSPs) in CA1 neurons in hippocampal slice cultures. Cells were recorded using patch electrodes (4-8MΩ) and EPSPs were evoked by Schaffer-collateral stimulation. Baseline EPSP recordings were kept short (≤ 5 minutes) to minimize dilution of postsynaptic factors that are necessary for LTP induction. Following baseline recording, we induced LTP by pairing presynaptic stimulation with postsynaptic depolarization (Figure 1A). Pairings were causal, in that presynaptic stimuli preceded postsynaptic spiking by 7-10ms. A single pairing was repeated 60 times at 5Hz. For postsynaptic depolarization, we injected current of sufficient amplitude to generate 3-6 postsynaptic spikes over a 50-60ms time course, with the first spike starting 7-10ms following the start of current injection. Spikes often tended to broaden over the time course of injection. The resulting waveform resembled a complex spike (Figure 1A), which is known to efficiently drive LTP *in vitro*^35-38^ and has been recorded in the hippocampus *in vivo*^39,40^. We found that this pairing protocol produced robust and reliable LTP (fold ΔEPSP_slope_: 1.88±0.24; n=6 cells; vs 1.0: p<0.05; Figure 1B,C), which had a presynaptic component of expression, as assessed by a decrease in the paired pulse ratio (PPR) (ΔPPR: -0.39±0.15; n=6 cells; vs 0: p<0.05; Figure 1D).

**Figure 1.**
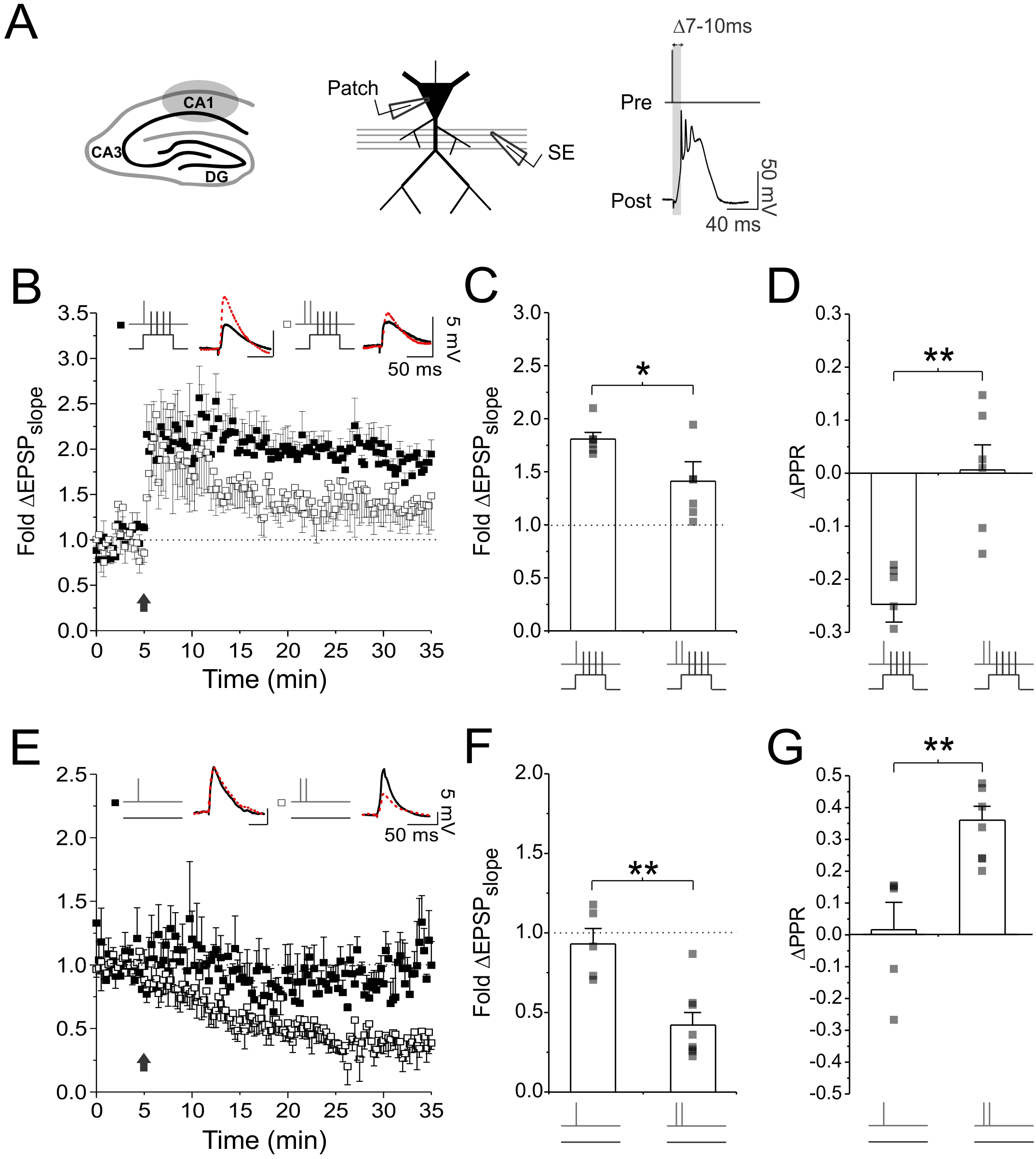
High frequency presynaptic activity inhibits presynaptic LTP and promotes presynaptic LTD. **(A)** Experimental setup. CA1 pyramidal neurons were recorded using whole-cell patch electrodes. LTP was induced by causal pairing of presynaptic activity with postsynaptic depolarization in the form of a complex spike. **(B,E)** Average fold changes of EPSP slope was plotted against time. Following baseline recording, 60 presynaptic stimuli were delivered at 5Hz, either in the presence (paired stimulation) or absence (unpaired stimulation) of postsynaptic depolarization. Presynaptic stimuli were delivered either as single pulses, or high frequency presynaptic bursts consisting of two pulses delivered 5ms apart. Sample EPSP traces at baseline (black trace) and 30 minutes after plasticity induction (broken red trace) are shown as an inset above. Average changes in **(C,F)** EPSP slope and **(D,G)** PPR measured 25-30 minutes after plasticity induction. High frequency bursts generated significantly less presynaptic LTP with paired stimulation, and more presynaptic LTD with unpaired stimulation, than did single pulse stimulation. Error bars represent S.E.M. (n=5-8 cells per condition). Asterisks denotes significance differences between groups (*p<0.05; **p<0.01; Mann-Whitney test).

We next examined the effects of elevating glutamatergic signalling during LTP induction on presynaptic plasticity. Under physiological conditions, elevated glutamate signalling arises from increased presynaptic activity. We therefore repeated our LTP experiments, but during induction, in the place of single presynaptic pulses, we delivered short, high frequency bursts of presynaptic stimuli to drive more glutamate release. The burst consisted of two pulses, delivered 5ms apart, and resembled high-frequency bursting activity recorded in CA3 neurons *in vivo*^41^. We found that this protocol produced significantly less LTP compared to single pulse pairings (fold ΔEPSP_slope_: 1.36±0.13; n=6 cells; vs. single pulse pairings: p<0.05), and was accompanied by no significant changes in PPR (ΔPPR: 0.00±0.04; n=6 cells; vs 0: p=0.84; vs. single pulse pairings: p<0.01; Figure 1D), suggesting that LTP was likely to be exclusively expressed postsynaptically. These findings suggest that high frequency presynaptic activity can inhibit the induction of presynaptic LTP.

We repeated our experiments, but during LTP induction, we omitted postsynaptic depolarization (unpaired stimulation). When single presynaptic pulses were delivered (60 pulses at 5 Hz) during induction, we observed no significant change in EPSP (fold ΔEPSP_slope_: 0.93±0.10; n=5 cells; vs 1.0: p=0.62; Figure 1E,F) or PPR (fold ΔPPR: 0.01±0.09; n=5 cells; vs 0: p=0.81; Figure 1F). However, when high frequency bursts were delivered during induction, we observed a robust decrease in the EPSP (fold ΔEPSP_slope_: 0.42±0.08; n=8 cells; vs. single pairing: p<0.01; Figure 1E,F) and an increase in PPR (fold ΔPPR: 0.36±0.04; n=8 cells; vs. single pulse pairing: p<0.01; Figure 1G), suggesting that we had induced LTD with a presynaptic component of expression. Collectively, these findings show that high frequency presynaptic stimulation not only inhibited the induction of presynaptic LTP, but also promoted the induction of presynaptic LTD.

### Glutamate photolysis inhibits presynaptic LTP and promotes presynaptic LTD

We next tested whether the effects of high frequency presynaptic stimulation on presynaptic plasticity were in fact due to the presynaptic terminal releasing more glutamate, as opposed to other effects, such as an increased Ca^2+^ influx in the presynaptic terminal. To do so, we used glutamate uncaging instead of high-frequency presynaptic stimulation to elevate glutamate release at synapses during LTP induction. Glutamate uncaging was restricted to single synapses. Activity-dependent changes in presynaptic function (i.e. P_r_) at these synapses was assessed by imaging postsynaptic Ca^2+^ transients^13,42^. This technique relies on the fact that at most CA3-CA1 synapses, single quanta of glutamate, through AMPAR-mediated depolarization, generates sufficient Ca^2+^ influx from NMDAR and voltage-gated Ca^2+^ channels (VGCCs) to be detected by Ca^2+^- sensitive dyes^1,43,44^. Consequently, the proportion of trials in which single presynaptic stimuli generate postsynaptic Ca^2+^ transients can be used to calculate P_r_ at single synapses^44^. Notably, estimates of P_r_ measured at resting membrane potential are resilient to reductions or enhancements in postsynaptic Ca^2+^ influx (Supplemental Figure 1, Supplemental Figure 8A-C).

CA1 pyramidal neurons were loaded with the Ca^2+^ sensitive dye Oregon Green BAPTA-1, and a fluorescently-labelled glass electrode was used to stimulate Schaffer-collaterals in the vicinity of the imaged dendrite (Figure 2A). Dendritic spines were sequentially scanned in order to identify those that were responsive to stimulation. To increase the likelihood of visually identifying responsive synapses, especially those with low basal release probabilities, we delivered two presynaptic stimuli, 70ms apart, to transiently increase P_r_ (Figure 2B). When a synapse was found that responded to stimulation, it always responded in an all-or-none manner, with Ca^2+^ transients largely restricted to the spine head. As expected, Ca^2+^ transients were also more likely to be elicited by the second of the two presynaptic stimuli because of the effects of short-term plasticity. P_r_ was calculated as the proportion of trials in which the first of the two presynaptic stimuli generated a fluorescent increase in the spine head; the second of the two presynaptic stimuli was ignored.

**Figure 2.**
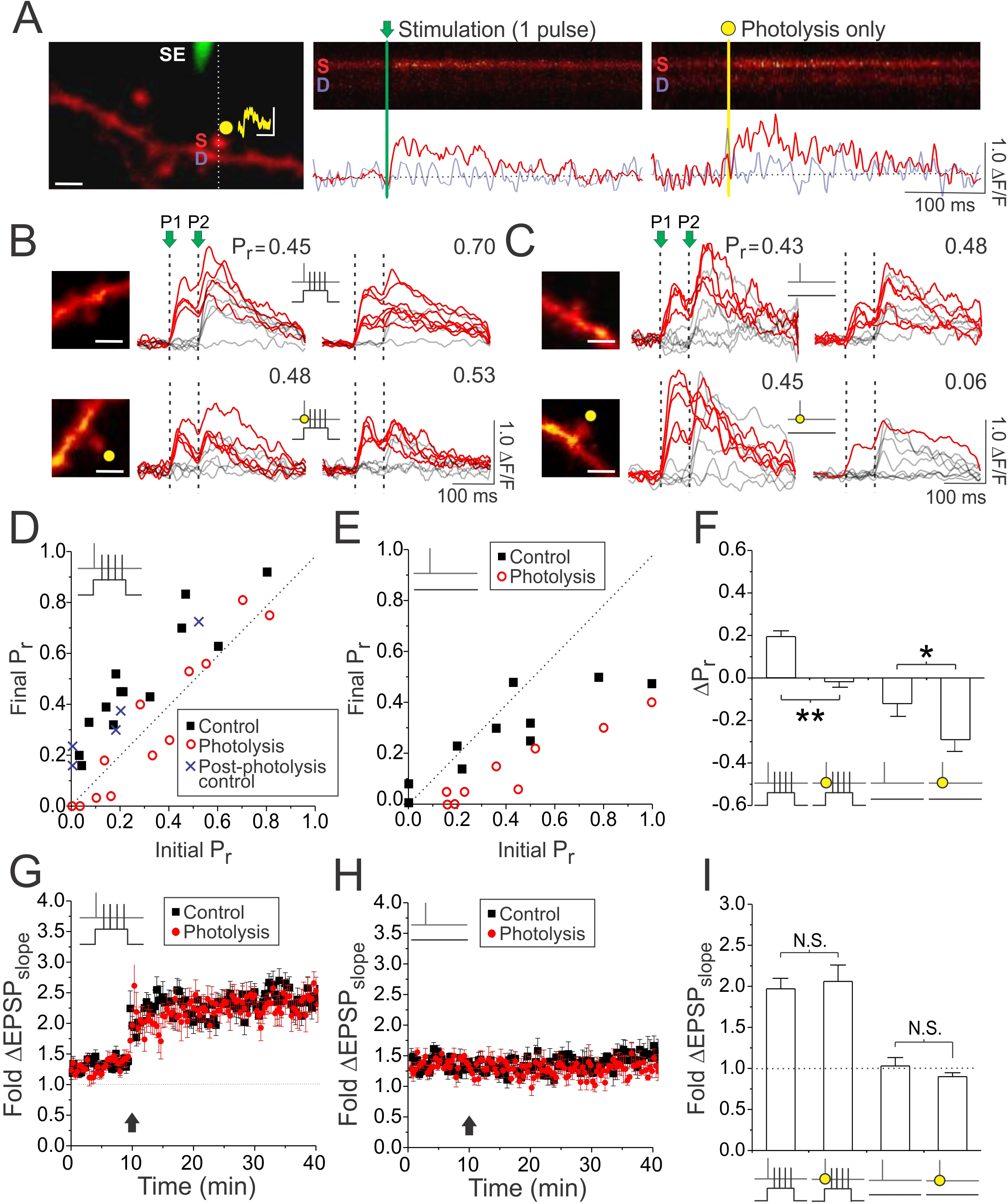
Glutamate photolysis inhibits presynaptic LTP and promotes presynaptic LTD. **(A)** Left, an image of a CA1 neuronal dendrite loaded with Oregon Green BAPTA-1 Ca^2+^- sensitive dye. A stimulating electrode (SE; green) was placed close to the dendrite in order to activate spines within the vicinity (scale bar: 2μm). A spine responsive to stimulation was located and then targeted for glutamate photolysis (yellow spot). An example of an uncaging-evoked synaptic potential is shown above the imaged spine (scale bar: 1mV by 100ms). During stimulation and photolysis, evoked Ca^2+^ transients were rapidly imaged by restricting laser scanning to a line across the spine head and underlying dendrite (broken line). Right, samples of these Ca^2+^ transients in both the spine (labelled S) and dendritic (labelled D) are shown. Below each line scan image are traces quantifying the fluorescence change (ΔF/F) for the spine (red trace; raw) and dendrite (purple trace; raw). Photolysis laser power was adjusted to elicit spine Ca^2+^ transients comparable to that induced by single presynaptic stimuli. **(B,C)** Samples of 10 superimposed Ca^2+^ traces evoked in imaged spines (white scale bar: 2μm) by paired pulse stimulation (P1 and P2 were delivered 70ms apart and are represented by vertical broken lines); red traces depict successful release events to the first (P1) of the two pulses. Ca^2+^ traces are shown during baseline and 25-30 minutes following paired or unpaired stimulation, delivered in either the absence or presence of glutamate photolysis (yellow circle). P_r_ was calculated as the proportion of total stimulation trials in which the first pulse (P1) resulted in a successful release event. Glutamate photolysis inhibited increases in P_r_ during paired stimulation, and promoted decreases in P_r_ during unpaired stimulation. **(D,E)** The final P_r_ measured 25-30 minutes following stimulation was plotted against the initial P_r_ measured at baseline for each imaged synapse. The broken diagonal line represents the expected trend if P_r_ was unchanged. The post-photolysis control group consisted of 5 synapses from the photolysis group that underwent a second round of paired stimulation but in the absence of glutamate photolysis. **(F)** Average changes in P_r_. **(G,H)** Average fold changes in EPSP slope was plotted against time for imaging experiments. The point at which paired or unpaired stimulation was delivered is denoted by the black arrow. **(I)** Average changes in EPSP slope. Error bars represent S.E.M. (n=9-14 spines per condition). Asterisks denotes significance differences between groups (*p<0.05; **p<0.01; Mann-Whitney test). N.S. denotes no significant difference.

Because of the additional time required to calculate P_r_, cells were recorded from using sharp microelectrodes (80-120MΩ) to minimize the dilution of postsynaptic contents. Following baseline measurements of P_r_, we induced LTP as before, by delivering 60 individual presynaptic stimuli at 5Hz, each paired with postsynaptic depolarization Consistent with our electrophysiological results, this protocol evoked an increase in P_r_ (ΔΡ_r_: 0.19±0.03; n=13 spines; vs. 0: p<0.01; Figure 2B,D,F) and a long-lasting potentiation of the recorded EPSP (fold ΔEPSP_slope_: 1.97±0.13; n=12 cells; vs 1.0: p<0.01 Figure 2G,I). We then repeated the experiment but this time, during LTP induction, each presynaptic stimulus was paired with photolysis of caged glutamate at the synapse. We adjusted the laser power to ensure that photolysis mimicked the fluorescent changes elicited by uniquantal glutamate release evoked by single presynaptic stimuli (ΔF/F; photolysis vs. stimulation: 0.38±0.08 vs. 0.43±0.09; n=12 spines; p=0.72; Figure 2A). Remarkably, under these conditions, increases in P_r_ at the target synapse were effectively abolished (ΔΡ_r_: - 0.02±0.03; n=12 spines; photolysis vs. control: p<0.01; Figure2B,D,F). This demonstrates that, consistent with our hypothesis, elevated glutamatergic signalling at synapses inhibited the induction of presynaptic LTP. Notably, in uncaging experiments, LTP induction resulted in a similar enhancement of the EPSP as in control experiments (fold ΔEPSP_slope_: 2.06±0.20; n=11 cells; photolysis vs. control: p=0.48; Figure 2G,I), suggesting that the failure for the imaged synapse to support LTP could not be attributed to the failure of the recorded cell or slice to support LTP. In five experiments, LTP induction was repeated for a second time at the same synapse, in the absence of caged glutamate, but with photolytic laser exposure; under these conditions, the expected increase in P_r_ was observed (ΔP_r_: 0.18±0.02; n=5 spines; vs. control: p=0.75; post-photolysis control in Figure 2D). Increases in P_r_ were also observed in a subset of control experiments, in which LTP induction was conducted in the presence of caged glutamate, but in the absence of photolytic laser exposure (ΔP_r_: 0.21±0.08; n=3 spines; vs. control: p=0.57). These results suggest that the inhibitory effect of photolysis on P_r_ was due to glutamate release, as opposed to non-specific effects of uncaging.

We also examined the effects of glutamate photolysis delivered in the absence of postsynaptic depolarization (unpaired stimulation). Delivery of 60 presynaptic stimuli at 5 Hz, as before, had no effect on the recorded EPSP (fold ΔEPSP_slope_: 1.03±0.10; n=9 spines; vs 1.0: p=0.94; Figure 2H,I), consistent with our previous result, and produced no changes in P_r_ at the majority of synapses imaged (Figure 2C,E,F). We did, however, notice that synapses with initially high release probabilities (P_r_>0.5), showed a modest decrease in P_r_ following unpaired stimulation (Figure 2E); this decrease was not likely to be detected by electrophysiological recordings because high P_r_ synapses comprise an estimated <10% of synapses in our preparation^42^. Remarkably, when we coupled each presynaptic stimulus with glutamate photolysis, we now observed decreases in P_r_ at all imaged synapses, regardless of their initial P_r_ (ΔP_r_ photolysis vs. control: -0.33±0.08 vs. -0.12±0.06; n=9,10 spines p<0.05; Figure 2C,E,F). These findings suggest that elevated glutamate release at a synapse had an inhibitory effect on P_r_. Notably, this inhibitory effect occurred regardless of the level of postsynaptic depolarization that accompanied presynaptic activity. P_r_ changes induced by paired or unpaired stimulation were always more negative compared to controls when glutamate signalling was augmented.

### Presynaptic LTP can be induced in complete glutamate receptor blockade

Given that augmenting glutamatergic signalling inhibited the induction of presynaptic LTP, we asked if glutamate release was required at all for driving increases in P_r_ during paired stimulation. Glutamate release is inevitably necessary to drive the postsynaptic spiking required for LTP, and all major classes of glutamate receptors including, AMPARs, Kainate receptors (KARs), NMDARs, and metabotropic receptors (mGluRs) can contribute to membrane depolarization^40,45-47^. However, if driving depolarization is the only function of glutamate the induction of presynaptic LTP, then we should be able to trigger presynaptic LTP in complete glutamate receptor blockade provided that, during induction, we replace the depolarizing-effects of glutamate with somatic current injection. If, however, glutamate release is additionally necessary for some form of synapse-specific signalling, as in the case of postsynaptic LTP, then the induction of presynaptic LTP should not be possible in complete glutamate receptor blockade.

To test this possibility we attempted to induce LTP at CA3-CA1 synapses in hippocampal slices with all known glutamate receptors (AMPARs, KARs, NMDARs, and mGluRs) pharmacologically inhibited (10μM NBQX, 100μM D-AP5, 0.5mM R,S-MCPG, 100μM LY341495). Given the additional time requirements for these experiments, we recorded from CA1 neurons using high-resistance patch electrodes (18-25MΩ) to limit dilution of postsynaptic factors. Following abolishment of the EPSP, synaptic activity was causally paired as before with strong postsynaptic depolarization triggered by somatic current injection (Figure 3A). The antagonist cocktail was washed out, and the EPSP was allowed to recover. As expected, drug washout was never complete (Figure 3C,D) and so it was necessary to compare the EPSP recorded from the pathway receiving paired stimulation to a second, independent control pathway recorded simultaneously (Figure 3A,B). We found that pairing induced a robust enhancement of the EPSP in the stimulated pathway (fold ΔEPSP_slope_; paired vs. control: 1.12±0.13vs. 0.71±0.12; n=7 cells; p<0.05; Figure 3B,D), which lasted for the duration of the recording (up to 40-90 minutes post-pairing); this enhancement was not seen when pairings were anti-causal, with presynaptic stimuli following postsynaptic spiking (Supplemental Figure 2). Causal pairings resulted in a 1.72±0.21 fold potentiation, which we estimated by normalizing the fold change in the EPSP of the paired pathway to that of the control pathway. Notably, EPSP recovery of the control pathway was not significantly different from experiments in which drugs were applied in the absence of paired stimulation (control vs. drugs-only: 0.71±0.12 vs. 0.54±0.11; n=7, 5 cells; p=0.59; Figure 3C,D), suggesting that LTP was restricted to only synapses that were active during the pairing.

**Figure 3.**
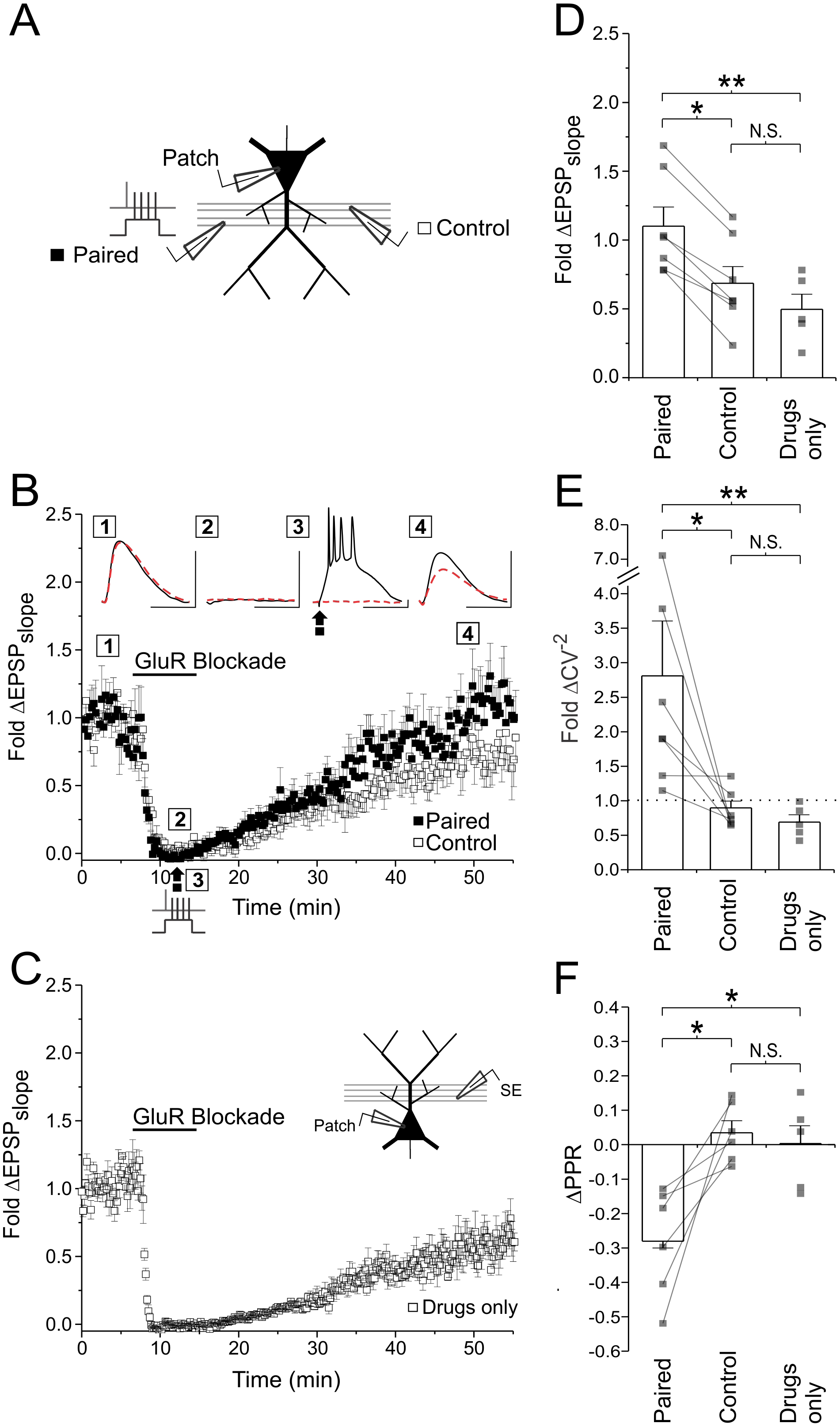
Induction of presynaptic LTP in glutamate receptor blockade. **(A)** Experimental setup. EPSPs were recorded from two independent Schaffer-collateral pathways. LTP was induced in a complete glutamate receptor blockade (100μM D-AP5, 10μM NBQX, 500μM R,S-MCPG, and 10μM LY341495) by paired stimulation. Only one pathway (black box) was active during paired stimulation, the other pathway served as a control (white boxes). **(B)** Average fold change in the EPSP slope is plotted against time for both the control and paired pathways. Sample EPSP traces shown are averages of 10 traces from the paired (solid line) and control (broken red line) pathway taken at four time points (1-4) from a single experiment (scale bar: 4.0mV by 40ms for paired pathway EPSP, 4.9mV by 40ms for control pathway EPSP). Stimulation artifacts have been removed for clarity. **(C)** Average fold change in EPSP slope is plotted for control experiments in which glutamate receptor antagonists were applied alone, in the absence of paired stimulation (drugs only group). Note that drug washout was incomplete. Group data and averages plotted for fold changes in **(D)** EPSP slope, **(E)** CV^-2^ and **(F)** PPR across experiments as measured 30 minutes following paired stimulation and drug washout. EPSP slope was higher in the paired pathway than in the control pathway, and was associated with an increase in CV^-2^ and a decrease in PPR, suggesting that LTP had been induced in the paired pathway, and had a presynaptic locus of expression. Error bars represent S.E.M (n=5-7 cells per condition). Asterisks denotes significance differences between groups (**p<0.01; *p<0.05; Wilcoxon signed rank test or Kruskal-Wallis with post-hoc Dunn’s test). N.S. denotes no significant differences between groups.

We next examined the locus of LTP expression. We found that LTP induction in glutamate receptor blockade was accompanied by a significant increase in the coefficient of variation parameter (CV^-2^) (paired vs. control: 2.80 ±0.79 vs. 0.89 ±0.10; n=7 cells; p<0.05; Figure 3E), and a significant decrease in PPR (paired vs. control ΔPPR: -0.28±0.06 vs. 0.03±0.03; n=6 cells; p<0.05; Figure 3F). Both of these changes are consistent with a presynaptic component of LTP expression, and both were found only in the paired pathway suggesting that LTP induction was site-specific. We also confirmed that similar site-specific presynaptic enhancements in recorded EPSPs could be induced under glutamate receptor blockade in acute hippocampal slices (Supplemental Figure 3 and 4).

Although several studies have demonstrated that NMDAR blockade alone impairs LTP induction, even presynaptically^11-15^, it is important to note that NMDARs, like all glutamate receptors, are known to contribute to postsynaptic depolarization. The NMDARs are particularly potent sources of depolarization especially given their role in dendritic spiking^45,48^ and burst firing^40^. Thus, it is possible that NMDAR blockade inhibits presynaptic LTP expression by inhibiting postsynaptic depolarization. This is less likely to be an issue when strong postsynaptic depolarization is driven via somatic current injection, as in our experiment (Figure 3), than when depolarization is driven by presynaptic stimulation. To test this, we induced LTP using standard 100Hz tetanic stimulation to drive postsynaptic spiking (Supplemental Figure 5A,D). This form of LTP had a presynaptic component of expression (Supplemental Figure 5E). As in previous studies, application of AP5 abolished LTP induction (Supplemental Figure 5B). However, we found that if tetanic stimulation was instead delivered while the postsynaptic cell was depolarized by 10mV (from -60mV to - 50mV) in order to augment postsynaptic spiking during tetanic stimulation, LTP induction in AP5 was rescued, at least presynaptically (Supplemental Figure 5C-E). These findings suggest that the importance of postsynaptic NMDAR signalling in presynaptic LTP induction is to provide a source of depolarization rather than any necessary source of synapse-specific signalling. These findings also underlie the importance of taking the level of postsynaptic depolarization into account when LTP is induced following the blockade of one or more glutamate receptor.

Collectively, our findings suggest that the role of glutamate signalling (including postsynaptic NMDAR signalling) in presynaptic LTP induction is to drive the postsynaptic depolarization, meaning that physiologically, presynaptic potentiation may not actually require any one presynaptic terminal to trigger glutamate release, provided that its activity coincides with postsynaptic depolarization, which could be triggered by glutamate release at other co-active synapses.

### LTP induction in glutamate receptor blockade is associated with an increase in P_r_

We then returned to Ca^2+^ imaging to determine whether we could directly observe increases in P_r_ at single synapses associated with the induction of LTP in full glutamate receptor blockade (Figure 4). To minimise dialysis for these experiments during drug wash-in, imaging was conducted in the absence of electrophysiological recordings on CA1 neurons that were pre-loaded with Ca^2+^ indicator dye (see Methods). Following an initial assessment of P_r_, glutamate receptor antagonists (10μM NBQX, 50μM D-AP5, 0.5mM R,S-MCPG, 100μM LY341495) were bath applied and the imaged cell was transiently patched in order to drive postsynaptic depolarization during paired stimulation. Consistent with electrophysiological findings, causal pairing of pre- and postsynaptic depolarization produced robust and reliable increases in P_r_ (ΔP_r_: 0.40±0.06; n=7 spines; vs 0: p<0.01; Figure 4A-C). No such changes were elicited by drug application in the absence of pairing (ΔP_r_: 0.04±0.04; n=7 spines; vs. causal pairing: p<0.05), or by either presynaptic stimulation alone (ΔP_r_: -0.03±0.03; n=7 spines; vs. causal pairing: p<0.01) or postsynaptic stimulation alone (ΔP_r_:: -0.01±0.04; n=6 spines; vs. causal pairing: p<0.01), or when postsynaptic depolarization preceded, rather than followed, presynaptic stimulation during pairing (ΔP_r_: -0.02±0.06; n=6 spines; vs. causal pairing: p<0.01) (Figure 4B,C). The induction of presynaptic LTP in the absence of glutamatergic signalling was therefore Hebbian, requiring presynaptic activity to be causally paired with strong postsynaptic depolarization.

**Figure 4.**
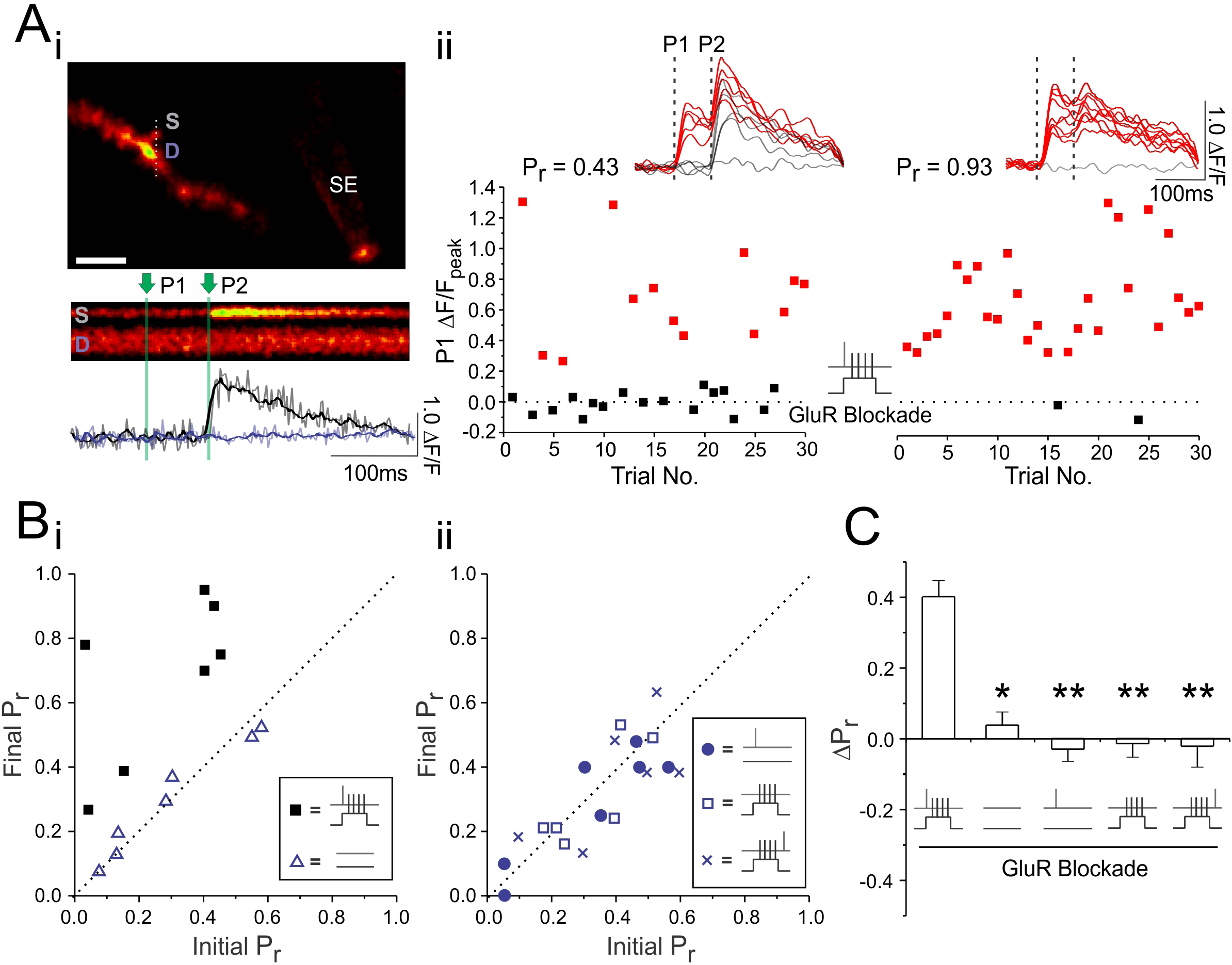
LTP induction in glutamate receptor blockade is associated with an increase in P_r_. **(Ai)** Example experiment. Top, an image of a CA1 neuronal dendrite loaded with Oregon Green BAPTA-1 Ca^2+^-sensitive dye with a stimulating electrode (SE) placed close to the dendrite in order to activate synapses within the vicinity (scale bar: 5μm). Example of Ca^2+^ transients were evoked by two stimulation pulses (P1 and P2; green vertical lines) are shown in the spine head (S) and underlying dendrite (D). Below the line scan are raw (grey) and smoothed (colored) traces quantifying fluorescence changes (ΔF/F) in the spine (black trace) and dendrite (purple trace). **(Aii)** The peak ΔF/F in the spine head following the first of the two stimulation pulses is plotted across 30 paired pulse stimulation trials given at baseline and 30 minutes following paired stimulation, which was delivered in complete glutamate receptor blockade (50μM D-AP5, 10μM NBQX, 500μM R,S-MCPG, and 10μM LY341495). Red squares denote fluorescent increases above noise. Smoothed Ca^2+^ traces from the last 10 trials are shown above each graph. P_r_ was calculated as the proportion of total stimulation trials in which the first pulse resulted in a fluorescent increase above noise. **(B)** Group data. For each experiment, the imaged synapse’s initial P_r_ is plotted against its final P_r_, calculated 30 minutes following one of five different stimulation paradigms. Only causal pairing of pre- and postsynaptic activity generated increases in P_r_ **(C)** Average change in P_r_. Error bars represent S.E.M. (n=6-10 spines per condition). Asterisks denotes significance differences between the first group in the graph (**p<0.01; *p<0.05; Kruskal-Wallis with post-hoc Dunn’s test).

### Postsynaptic depolarization increases P_r_ by promoting dendritic release of NO in a manner dependent on L-VGCCs

We next investigated the mechanism by which paired stimulation could trigger increases in P_r_ in the absence of glutamatergic signalling. The requirement for postsynaptic depolarization in the induction of presynaptic potentiation suggests a need for a diffusible retrograde messenger. Perhaps the most promising, albeit still controversial, retrograde signal implicated in LTP induction is nitric oxide (NO)^6^. Although NO synthesis has classically been associated with the activation of postsynaptic NMDARs^8^, there is some suggestion that Ca^2+^ influx from L-type voltage-gated Ca^2+^ channels (L-VGCCs), which have previously been implicated in presynaptic LTP ^18,19^, could trigger NO production^22,49,50^. If NO synthesis in neuronal dendrites can be triggered by L-VGCC activation, then NO production could occur in a manner dependent on postsynaptic depolarization, but independent of synapse-specific glutamatergic signalling. To test this, we first asked whether presynaptic LTP, induced in glutamate receptor blockade, was dependent on L-VGCC activation and NO signalling. Consistent with our hypothesis, we found that pairing-induced increases in P_r_ (ΔP_r_: 0.34±0.04; n=10 spines; p<0.01) were reliably abolished by bath application of the L-VGCC antagonist nitrendipine (20μM) (ΔP_r_: 0.04±0.04; n=6 spines; vs. blockade: p<0.01) and by the NO scavenger carboxy-PTIO (cPTIO), either bath applied (50-100μM) (ΔP_r_: -0.02±0.04; n=7 spines; vs. blockade: p<0.01) or injected into the postsynaptic neuron (5mM bolus-loaded) (ΔP_r_: -0.02±0.07; n=6 spines; vs. blockade: p<0.05) (Figure 5A). The NO synthase inhibitor, L-NAME (100μM) also blocked presynaptic enhancements in recorded EPSPs (Supplemental Figure 6). We confirmed our findings in acute slices, and found that nitrendipine and cPTIO blocked presynaptic enhancements induced under glutamate receptor blockade (Supplemental Figure 3 and 4), suggesting that, as in cultured slices, presynaptic efficacy in acute slices was similarly regulated by NO signalling.

We then examined whether NO production depended on L-VGCC activation. We transiently patched CA1 neurons in order to load them with the NO-sensitive dye, DAF-FM (250μM bolus-loaded), and then measured fluorescent changes in the apical dendrites prior to and following postsynaptic depolarization in glutamate receptor blockade. Given the poor signal-to-noise ratio associated with DAF-FM imaging, we drove strong postsynaptic depolarization by elevating extracellular K^+^ to 45mM, as previously described^49,50^. Under these conditions, we observed increases in fluorescence in neuronal dendrites (Figure 5B,C). These increases were dependent on NO synthesis as they could be prevented by postsynaptic injection of cPTIO (ΔF/F; control vs. cPTIO: 0.38±0.04 vs. - 0.03±0.05; n=5 cells/condition; p<0.05) or bath application of the NO synthase (NOS) inhibitor L-NAME (ΔF/F: 0.00±0.05; n=5 cells; vs. control: p<0.05). Importantly, fluorescent increases were reliably abolished with nitrendipine (ΔF/F: -0.02±0.06; n=5 cells; vs. control: p<0.05) (Figure 5B,C), suggesting that NO synthesis required L-VGCC activation.

We then attempted to image NO release in response to more physiologically-relevant means of postsynaptic stimulation, such as the complex spikes we were using to induce LTP. To do so we pre-loaded slices with 1,2-Diaminoanthraquinone (DAQ; 100μg/mL), a NO-sensitive dye, as previously described^51^. We then patched onto a single cell and imaged the DAQ-associated changes in the cell after stimulating the cell with 600 complex spikes delivered at 5Hz (Figure 9D). Stimulation was performed in complete glutamate receptor blockade. This protocol took advantage of the fact that DAQ forms an insoluble fluorescent precipitate upon reacting with NO, meaning that fluorescence would accumulate with stimulation and not readily wash away^52^. We found with 600 complex spikes, the accumulated NO signal in the dendritic arbour was sufficiently large to detect by our setup (Figure 9D,E). Notably, no signal was detected in the absence of any stimulation (ΔF/F; stimulated vs. unstimulated: 2.97±0.48; vs 0.22±0.73; n=9,7 cells; p<0.05), or when stimulation was delivered in the presence of nitrendipine (ΔF/F: - 0.14±0.65; n=7 cells; vs. control: p<0.05) or L-NAME (ΔF/F: 0.24±0.43; n=8 cells; vs. control: p<0.05). These findings suggest that postsynaptic depolarization alone can drive NO release from neuronal dendrites in a L-VGCC dependent manner.

### NO release induces increases in P_r_ at active presynaptic terminals

Once NO is released, is it alone sufficient to induce LTP at active presynaptic terminals? To address this, we examined whether increases in P_r_ could be elicited when presynaptic stimulation was paired with rapid photolytic release of NO (0.5-1mM RuNOCl_3_), in the absence of postsynaptic depolarization. We used Ca^2+^ imaging to determine basal P_r_ at a single synapse. We then paired 30-60 presynaptic stimuli, delivered at 5Hz in complete glutamate receptor blockade, with brief photolysis of NO, which was directed targeted to the spine head to emulate postsynaptic NO release. As with our standard LTP induction protocol, pairing was causal, with each NO photolysis event timed to occur 7-10ms after each presynaptic stimulus. Under these conditions, we found significant increases in P_r_ when assessed 30 minutes post-pairing (ΔP_r_: 0.29±0.07; n=10 spines; p<0.01; Figure 5F-H). No such changes were produced when pairing occurred in the presence of bath-applied cPTIO (ΔP_r_: 0.02±0.07; n=6 spines; vs. causal pairing: p<0.05; Figure 5G,H), suggesting that LTP did not result from non-specific effects associated with photolysis. Moreover, when presynaptic stimuli followed NO photolysis no significant change in P_r_ was observed (ΔP_r_: -0.01±0.04; n=8 spines; vs. causal pairing: p<0.01; Figure 5F-H), suggesting that NO-mediated potentiation was Hebbian, requiring presynaptic activity to precede, rather than follow, NO release.

**Figure 5.**
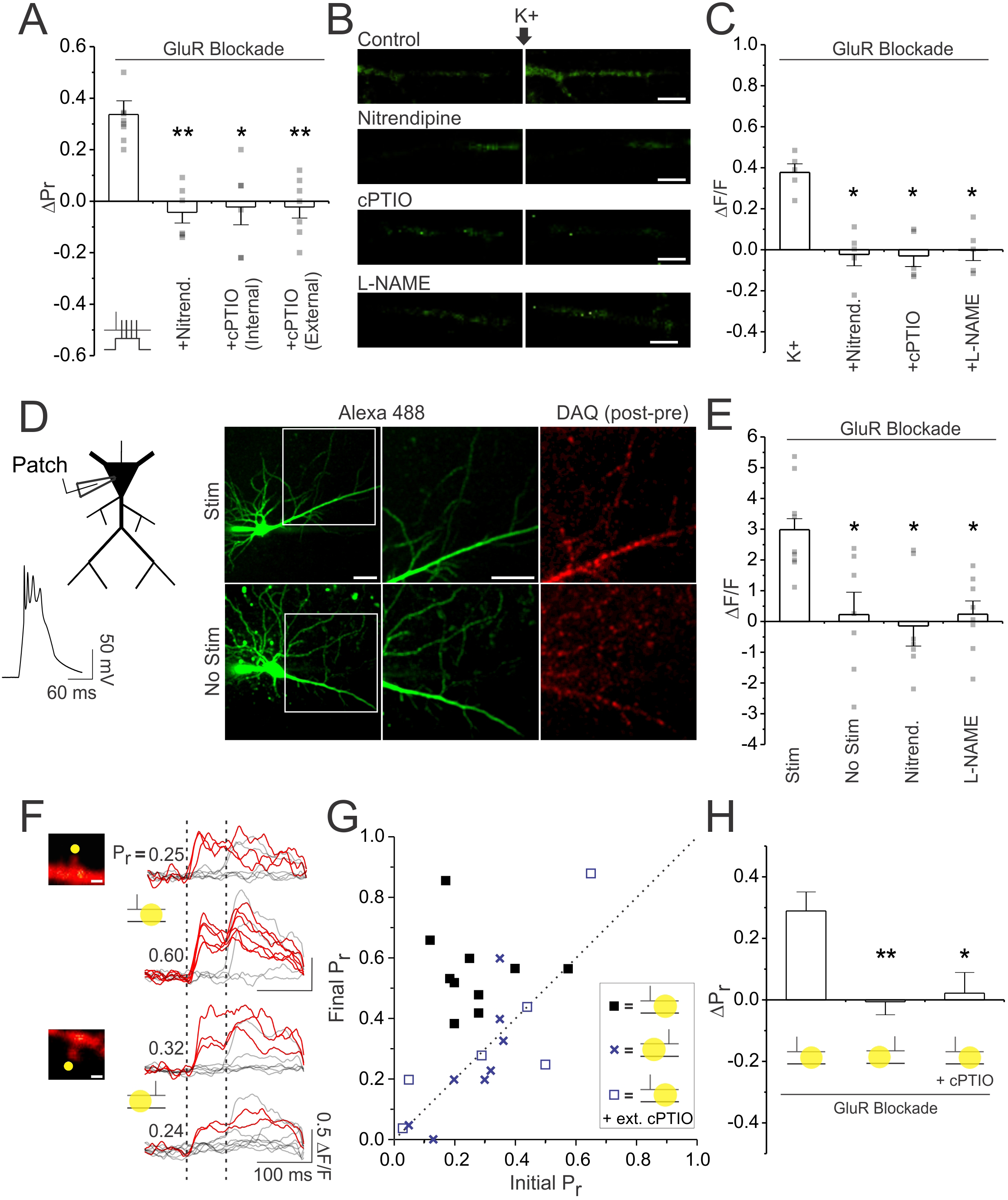
Postsynaptic depolarization increases Pr by promoting dendritic release of NO in a manner dependent on L-VGCCs. **(A)** Average changes in P_r_. Paired stimulation was delivered in full glutamate receptor blockade (50μM D-AP5, 10μM NBQX, 500μM R,S-MCPG, and 10μM LY341495), under control conditions, following treatment with the L-VGCC antagonist nitrendipine (20μM), or following either the bath (100μM) or intracellular (5mM bolus-loaded) application of the NO scavenger cPTIO. **(B)** Images of CA1 apical dendrites loaded with NO-sensitive dye DAF-FM (250μM bolus-loaded), prior to and following K^+^ (45mM) mediated depolarization (scale bar: 5μm) in control conditions, or in the presence of nitrendipine, cPTIO, or the NO synthesis inhibitor L-NAME (100μM; preincubation). K^+^ stimulation evoked NO-sensitive and L-VGCC dependent increases in DAF-FM fluorescence. **(C)** Average K^+^ induced fluorescence change (ΔF/F) of DAF-FM in apical dendrites. **(D)** Slices were preloaded with the NO-sensitive dye DAQ. Right. Example images of CA1 pyramidal neurons patched and loaded with Alexa Fluor 488 are shown, along with the associated change in DAQ fluorescence (post-pre stimulation images) recorded either following stimulation with 600 complex spikes (Stim) or following no stimulation (No stim). Images in white box are magnified in the adjacent image (scale bar: 5μm). Left. Example of the complex spikes used during stimulation. **(E)** Average K^+^ induced fluorescence change (ΔF/F) of DAF-FM in apical dendrites. Stimulation evoked NO-sensitive and L-VGCC dependent increases in DAQ fluorescence. **(F)** Samples of 10 superimposed Ca^2+^ traces evoked in imaged spines (scale bar: 1μm) by paired pulse stimulation (broken vertical bars); red traces depict successful release events to the first of the two pulses. Samples are taken from baseline and 25-30 minutes following a stimulation paradigm. The paradigm consisted of delivering presynaptic stimuli either (top) 7-10ms before or (bottom) 7-10ms after NO photolysis at the synapse (yellow circle); photolysis occurred in glutamate receptor blockade and in the absence of postsynaptic depolarization. Pr was calculated as the proportion of total stimulation trials in which the first pulse resulted in a successful release event. In some experiments NO photolysis was conducted in cPTIO. Only causal pairings of presynaptic activity and NO release led to increases in Pr. **(G)** Final Pr measured 25-30 minutes following the stimulation paradigm is plotted against the initial Pr for each synapse. The broken diagonal line represents the expected trend if Pr was unchanged. **(H)** Average change in Pr. Error bars represent S.E.M. (n=5-13 spines/cells per condition). Asterisks denote significant differences from the control group (**p<0.01; *p<0.05; Kruskal-Wallis with post-hoc Dunn’s test).

Previously, the effects of NO have primarily been examined by recording EPSPs in acute slices^6^. We therefore sought to confirm our findings using NO photolysis in a similar preparation (Supplemental Figure 7). We loaded CA1 pyramidal neurons in acute slices with caged NO (100μM RuNOCl_3_; bolus-loaded) while recording EPSPs in the presence of AP5. Wide-field photolysis was triggered using a 1ms flash from a UV lamp. Causal pairings of presynaptic activity with photolysis resulted in an enhancement of the EPSP and a decrease in PPR. These increases were absent when pairings were anti-causal, or when pairings occurred in the presence of cPTIO, which instead resulted in a modest depression of the EPSP. These findings confirm that NO can trigger presynaptic LTP provided that its release it precedes rather than follows presynaptic activity.

### Glutamate release decreases P_r_ via activation of presynaptic NMDARs

Our findings suggest that a presynaptic terminal need not release glutamate in order to become potentiated, provided that its activity precedes strong postsynaptic depolarization. In fact, in our initial experiments we found that glutamate release, if anything, inhibited presynaptic LTP and promoted presynaptic LTD (Figure 1,2). Theses findings, however, were based on elevating glutamatergic signalling at the synapse either by using high-frequency presynaptic bursts or glutamate photolysis. We therefore sought to examine whether endogenous glutamate release also had a similar inhibitory effect on P_r_. To do so we conducted single-spine Ca^2+^ imaging experiments in control conditions and under glutamate receptor blockade to examine how changes in P_r_ were affected by glutamate signalling. Remarkably, we found that increases in P_r_ produced in glutamate receptor blockade were significantly larger than that of controls (ΔP_r_; blockade vs. control: 0.34±0.04 vs. 0.18±0.02; n=10 spines; p<0.05; Figure 6A-C), suggesting that even endogenously released glutamate impeded increases in P_r_ induced by paired stimulation. We also examined the effects of glutamate receptor blockade on unpaired stimulation, during which presynaptic stimuli were delivered alone. This protocol reliably induced decreases in P_r_ at synapses with high release probabilities (P_r_>0.5) under control conditions. However, no such decreases were observed in glutamate receptor blockade (ΔP_r_ blockade vs. control: 0.00±0.03 vs. -0.21±0.05; n=10, 9 spines; p<0.05; Figure 6D,E). These findings suggest that endogenous glutamate release has an inhibitory effect on P_r_, regardless of the level of postsynaptic depolarization. Across conditions, P_r_ changes were always more positive compared to controls when glutamate signalling was blocked, suggesting that endogenous glutamate release drives decreases in P_r_.

**Figure 6.**
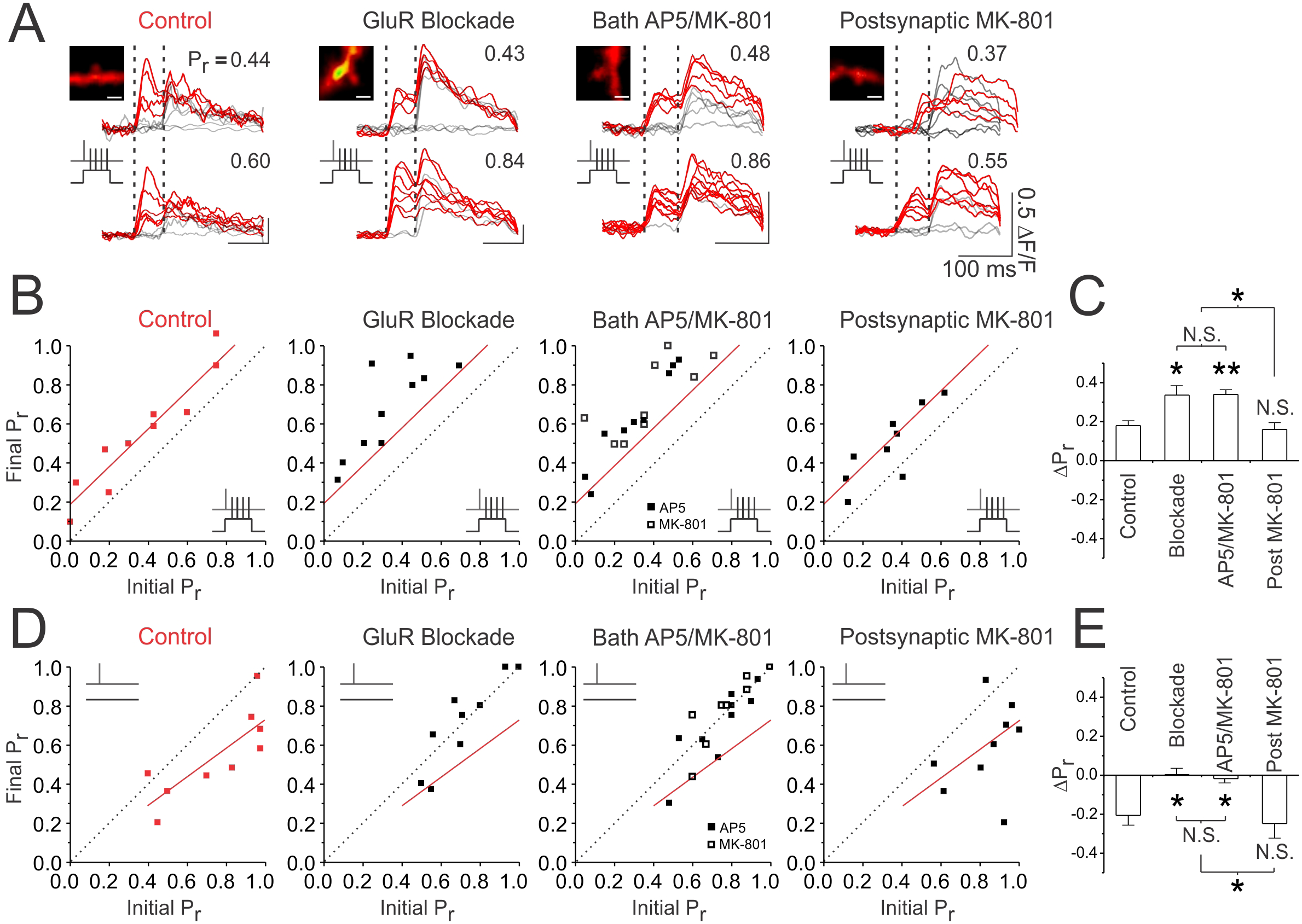
Glutamate release decreases Pr by activating presynaptic NMDARs. **(A)** Samples of 10 superimposed Ca^2+^ traces evoked in imaged spines (white scale bar: 1μm) by paired pulse stimulation (2 stimuli delivered 70ms apart; represented by the vertical broken lines); red traces depict successful release events to the first of the two pulses. P_r_ was calculated as the proportion of total stimulation trials in which the first pulse resulted in a successful release event. For each spine, sample Ca^2+^ traces are shown during baseline and 25-30 minutes following paired stimulation. Experiments were also conducted with unpaired stimulation, in which presynaptic activity was delivered in the absence of postsynaptic depolarization. **(B, D)** Paired or unpaired stimulation was delivered in control conditions or following: complete glutamate receptor blockade (50μM D-AP5, 10μM NBQX, 500μM R,S-MCPG, and 10μM LY341495), bath application of AP5 (D-AP5; 50μM)/MK-801 (20μM), or intracellular application of MK-801 postsynaptically (5mM bolus-loaded). For each experiment, the final P_r_ measured 25-30 minutes following (B) paired or (E) unpaired stimulation is plotted against the initial P_r_ measured at baseline. Red trendlines have been fitted to control data to guide visual comparison. The broken diagonal line represents the expected trend if P_r_ was unchanged. Inhibition of presynaptic NMDARs by full glutamate receptor blockade or by bath application of AP5 or MK-801 augments increases in P_r_ produced by paired stimulation and prevents decreases in P_r_ produced by unpaired stimulation. **(C,E)** Average changes in Pr. Changes in P_r_ were always more positive under conditions in which presynaptic NMDARs are blocked, suggesting that presynaptic NMDAR signalling was driving decreases in P_r_. Error bars represent S.E.M. (n=9-18 spines per condition). Asterisks denote significant differences (**p<0.01; *p<0.05; Kruskal-Wallis with post-hoc Dunn’s test). N.S. denotes no significant difference. All comparisons are made against control data unless otherwise specified in the figure.

How might glutamate release exert an inhibitory effect on P_r_? We have previously reported that presynaptic NMDARs are found at CA3-CA1 synapses; these receptors act as reliable detectors for uniquantal glutamate release^30^, and have been implicated in presynaptic LTD^31-33,53-55^. We therefore examined whether these receptors mediated the inhibitory effects of glutamate observed on presynaptic function. Given the difficulties associated with selectively blocking pre-, as opposed to post-, synaptic NMDARs, several groups have investigated the role of presynaptic NMDARs in plasticity by comparing the effects of bath application of 50μM D-AP5 or 20μM MK-801, which blocks both pre- and postsynaptic NMDARs, with that of intracellular MK-801 (5mM bolus-loaded) application, which selectively blocks postsynaptic NMDARs^55-58^. We sought to use a similar approach. However, because MK-801 does not readily washout, and since postsynaptic NMDARs greatly contribute to spine Ca^2+^ influx^43,44,59^, we first examined whether the permanent loss of postsynaptic NMDAR signalling affected our ability to measure P_r_ using postsynaptic Ca^2+^ imaging. We found that at about 50% of synapses, NMDAR blockade reduced, but did not entirely abolish synaptically-evoked Ca^2+^ transients (Supplemental Figure 8). The residual Ca^2+^ transients were mediated by activation of voltage-gated Ca^2+^ channels (VGCCs) in response to AMPAR-mediated depolarization, and could be used to accurately measure P_r_ (Supplemental Figure 8). Importantly, the average P_r_ of these synapses did not significantly differ from that of synapses lacking a residual Ca^2^+ transient in NMDAR blockade (ΔPr; AP5-sensitive vs. AP5-insensitive: 0.42±0.07 vs. 0.47±0.11; n=8 spines/condition; p=0.67; Supplemental Figure 8BC). These findings suggest that, in NMDAR receptor blockade, VGCC-dependent spine-Ca^2+^ influx can be used as a means of calculating P_r_ at a sizeable and representative proportion of presynaptic terminals; nonetheless the use of VGCC-dependent Ca^2+^ transients presents an unavoidable selection bias in our study.

Using VGCC-dependent spine Ca^2+^ transients, we found that when both pre- and postsynaptic NMDARs were blocked by bath application of either AP5 (n=9 spines) or MK- 801 (n=9 spines), paired stimulation triggered increases in P_r_ (ΔPr: 0.34±0.03; n=18 spines) that were not significantly different from those produced in complete glutamate receptor blockade (p>0.99), but that were greater than increases produced under control conditions (p<0.01) (Figure 6A-C). Bath application of AP5 or MK-801, like glutamate receptor blockade, also blocked decreases in P_r_ produced by unpaired stimulation (ΔP_r_: - 0.02±0.02; n=17 spines; vs. control; p<0.05; vs. blockade: p>0.99; Figure 6D,E). To then specifically block postsynaptic NMDARs, we bolus-loaded cells intracellularly with MK-801 (see Methods). This loading protocol minimized dilution of postsynaptic factors, and in control experiments, abolished NMDAR-mediated EPSPs and spine Ca^2+^ transients (Supplemental Figure 9A-C). In contrast to bath application of AP5 or MK-801, we found that with postsynatpic application of MK-801, increases in P_r_ produced by paired stimulation (ΔP_r_: 0.16±0.04; n=9 spines; vs. control: p>0.99; Figure 6A-C) and decreases in P_r_ produced by unpaired stimulation (ΔP_r_: -0.25±0.07; n=9 spines; vs. control: p>0.99; Figure 6D,E) did not significantly differ from control conditions (p>0.99), and were significantly different from changes in P_r_ induced in glutamate receptor blockade (p<0.05) and extracellular NMDAR blockade (p<0.05). Collectively, these results suggest that pre-, but not post-, synaptic NMDARs mediate the inhibitory effect of glutamate release on changes in P_r_ observed with both paired and unpaired stimulation.

The inhibitory effect of presynaptic NMDARs on P_r_ would also explain why glutamate photolysis in our earlier experiments (Figure 2) inhibited presynaptic LTP and promoted presynaptic LTD. To confirm this, we repeated photolysis experiments in the presence of MK-801, either intracellularly or extracellularly applied, again to differentially block presynaptic and postsynaptic NMDAR signalling (Figure 7). Consistent with our hypothesis, we found that bath, but not intracellular, application of MK-801 rescued LTP induction, despite the presence of uncaged glutamate at the synapse during postsynaptic depolarization (ΔP_r_: bath MK-801 vs. photolysis: 0.37±0.05; vs. -0.02±0.02 n=7, 12 spines; p<0.01; intracellular MK-801: -0.02±0.04; n=7 spines; vs. photolysis: p>0.99; Figure 7A,B). Incidentally, increases in P_r_ were not only rescued by bath application of MK-801 but were greater than in control conditions (ΔP_r_: control: 0.20±0.03; n=12 spines; vs. bath MK-801: p<0.01; Figure 7A,B). This is likely because bath application of MK-801, in addition to blocking the inhibitory effects of caged glutamate, additionally blocked the inhibitory effect of endogenous glutamate release that was present in control conditions (Figure 6B,C). As expected, bath, but not intracellular, application of MK-801 also prevented photolysis-induced augmentation of LTD, in which stimuli were delivered in the absence of postsynaptic depolarization (ΔP_r_: bath MK-801 vs. photolysis: -0.02±0.03; vs. -0.29±0.06; n=7,9 spines; p<0.01; intracellular MK-801: -0.28±0.08; n=7 spines; vs. photolysis: p>0.99; Figure 7C,D). Bath MK-801 application additionally prevented LTD induction present at high P_r_ synapses under control condition (Figure 7C,D) again, likely by blocking the inhibitory effects of endogenous glutamate release (Figure 6D,E). Collectively, these findings suggest that activation of presynaptic NMDARs are necessary to drive decreases in P_r_, regardless of the levels of postsynaptic depolarization.

**Figure 7.**
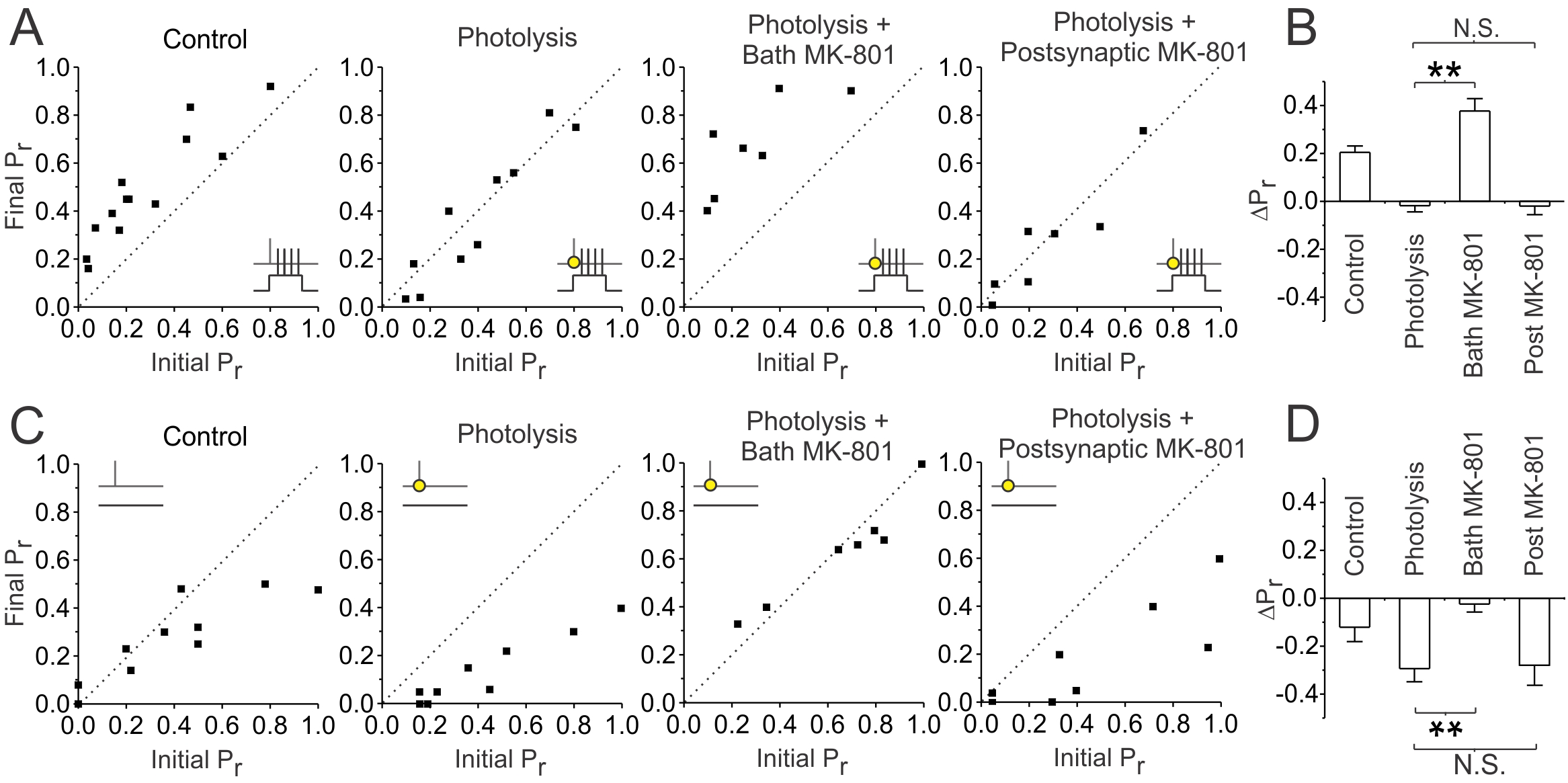
Inhibition of presynaptic NMDARs rescues the effects of glutamate photolysis on presynaptic plasticity. **(A, C)** Photolysis experiments conducted in Figure 1 are shown, in which paired or unpaired stimulation was delivered in the presence or absence of glutamate photolysis. Experiments were repeated following either the bath (20μM) or intracellular application (5mM; bolus-loaded) of MK-801. For each experiment, the final P_r_ measured 25-30 minutes following (A) paired or (C) unpaired stimulation is plotted against the initial P_r_ measured at baseline. The broken diagonal line represents the expected trend if P_r_ was unchanged. Bath, but not intracellular, application of MK-801 prevented photolysis-induced inhibition of LTP and photolysis-induced augmentation of LTD, suggesting that the effects of photolysis are mediated by presynaptic NMDAR signalling. **(B,D)** Average change in P_r_. Error bars represent S.E.M. (n=7-13 spines per condition). Asterisks denote significant differences from (**p<0.01; Kruskal-Wallis with post-hoc Dunn’s test). N.S. denotes no significant differences.

Our results provide strong evidence that presynaptic NMDAR activation by glutamate drives decreases in P_r_. We next sought to confirm these findings electrophysiologically (Figure 8). EPSPs generated by Schaffer-collateral stimulation were recorded at baseline and following paired stimulation to induce LTP. As with Ca^2+^ imaging, experiments were conducted under control conditions, following bath application of AP5 or MK-801, or following postsynaptic application of 1mM MK-801, which was included in the recording electrode. We found that paired stimulation induced robust LTP in the presence of bath applied AP5 or MK-801, which was not significantly different in magnitude than that produced under control conditions (fold ΔEPSP_slope_; bath AP5/MK-801 vs. control: 1.91±0.13 vs. 2.07±0.10; n=8,12 cells; p>0.99; Figure 8A,B). However, given that NMDAR blockade is known to inhibit postsynaptic LTP, the entirety of the LTP induced under NMDAR blockade was likely to be presynaptic in expression as compared to the mixed pre- and postsynaptic expression of LTP that was likely to arise under control conditions^16,18^. This would suggest that the presynaptic component of LTP was actually larger following the extracellular blockade of NMDARs. Indeed, LTP induced following bath application of AP5 or MK-801 was associated with a much larger decrease in PPR than that induced under control conditions (ΔPPR; bath AP5/MK-801 vs. control: -0.48±0.08 vs. - 0.24±0.03; n=8,12 cells; p<0.05; Figure 8C), consistent with the inhibitory role of presynaptic NMDARs in presynaptic LTP.

**Figure 8.**
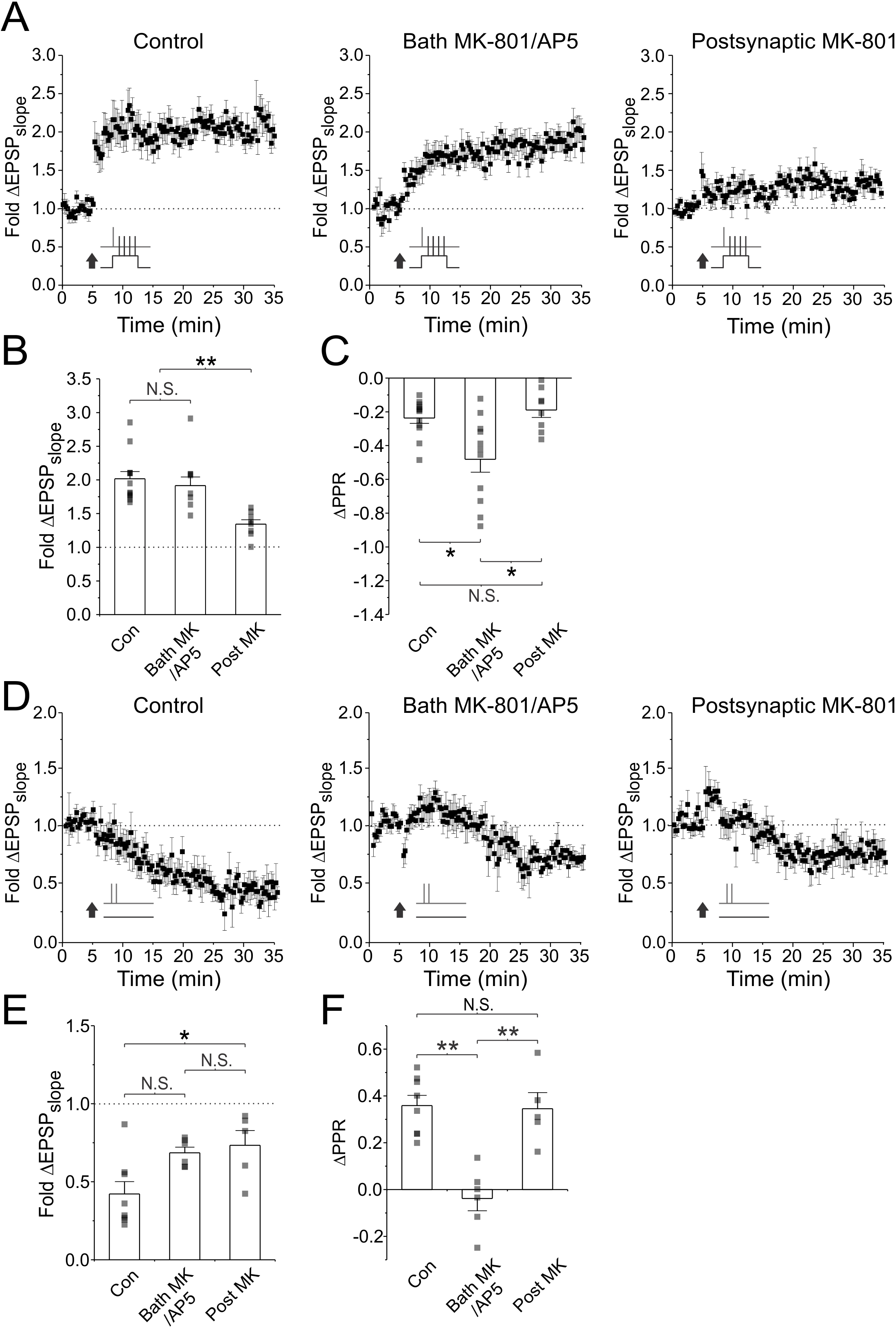
Inhibition of presynaptic NMDARs augments presynaptic LTP and abolishes presynaptic LTD. **(A,D)** Average fold change in EPSP slopes are plotted against time for plasticity induction under control conditions, after bath application of MK-801 (20μM) or AP5 (D-AP5; 50μM), and after bolus-loading of the postsynaptic neuron with MK-801 (5mM). Plasticity was induced either using paired stimulation (60 pairings at 5Hz) to induce LTP, or by delivering high frequency bursts of presynaptic stimuli (2 pulses at 200Hz repeated 60 times at 5Hz) in the absence of postsynaptic depolarization to induce LTD. (B,E) Average EPSP slope changes across experiments. **(C,F)** Average PPR changes across experiments. Bath application of AP5/MK-801, but not intracellular application of MK-801, augmented increases in PPR produced by paired stimulation, and abolished decreases in PPR produced by high frequency presynaptic stimulation alone. This suggests that presynaptic NMDAR signalling augmented presynaptic LTP and abolished presynaptic LTD. Error bars represent S.E.M. (n=5-12 cells per condition). Asterisks denote significant differences from (*p<0.05;**p<0.01; Kruskal-Wallis with post-hoc Dunn’s test). N.S. denotes no significant differences.

Surprisingly, paired stimulation under conditions of postsynaptic NMDAR blockade with intracellular application of MK-801 failed to induce LTP (Supplemental Figure 9D-F). This was unexpected given 1) LTP induction was successful following complete blockade of NMDARs by external application of MK-801 and 2) that we were able to obtain increases in P_r_ with paired stimulation following application of postsynaptic MK-801 in our Ca^2+^ imaging experiments (Figure 6B,C). However, in our imaging experiments we were bolus-loading MK-801 to allow for imaging to take place in the absence of chronic whole-cell patch recordings. Bolus-loading entailed patching on to the cell with 5mM MK-801 in the patch electrode for 60s before patching off again. The drug was then given approximately 20 minutes to diffuse and take effect prior to starting the experiment. In the current set of electrophysiological experiments, MK-801 was instead constantly present at 1mM in the recording electrode, and therefore would have been diffusing into the cell for 5 minutes at the point of LTP induction. We reasoned that this protocol may have resulted in a higher intracellular concentration of MK-801 as compared to bolus-loading. MK-801 is a highly potent NMDAR antagonist with a K_d_ value in the low nanmolar range (approximately 37nM)^60^. At high concentrations (> 100 μM), however, it can inhibit voltage-gated K^+^ and Ca^2+^ channels^61,62^. We were therefore concerned that chronic patch loading of 1mM MK- 801 may have disrupted L-VGCC function, which we had found to be necessary for presynaptic LTP induction (Figure 5A-E). To test this possibility, we imaged L-VGCC-mediated Ca^2+^ influx in dendrites in response to back-propagating action potentials (Supplemental Figure 9G-I). We found that 1mM patch loading of MK-801 for 5 minutes reduced L-VGCC-mediated Ca^2+^ influx by approximately 50%. Our bolus-loading procedure, in contrast, produced no change in L-VGCC function despite effectively inhibiting NMDAR function (Supplemental Figure 9A-C). Our findings suggest that there is a serious confound for loading high concentrations of MK-801 postsynaptically, particularly in LTP experiments.

Based on our findings, we switched our MK-801 loading protocol to bolus-loading. Under these conditions paired stimulation was able to successfully induce LTP (fold ΔEPSP_slope_: 1.34±0.07; n=8 cells; p<0.01; Figure 8A,B). LTP induction was smaller in magnitude compared with control conditions (p<0.01), though this is not surprising given that LTP induction under conditions of NMDAR-blockade was likely to only have a presynaptic component of expression, as compared to the mixed pre- and postsynaptic components of expression induced under control conditions^16,18^. Notably, LTP induction in postsynaptic NMDAR blockade was associated with a comparable decrease in PPR as seen in controls (fold ΔPPR: -0.19±0.04; n=8 cells; vs. control: p>0.99; Figure 8C), suggesting that the presynaptic component of LTP did not differ in magnitude between the two conditions. However, when compared to results obtained under extracellular blockade of NMDARs, LTP induced in postsynaptic NMDAR blockade was of smaller magnitude (p<0.01), and associated with smaller decreases in PPR (p<0.05). Consistent with our imaging experiments, these findings suggest that pre-, but not post- synaptic NMDAR activation reduces the magnitude of presynaptic LTP induced by paired stimulation.

We also examined the effects of presynaptic NMDARs on LTD induction induced by unpaired stimulation. For unpaired stimulation, we used our high-frequency burst protocol (Figure 1), since it produced robust depression in recorded EPSPs. We delivered 60 bursts of presynaptic stimuli (2 stimuli delivered 5ms apart) at 5Hz in the absence of depolarization. This induced comparable levels of LTD across conditions, though LTD induced following extracellular or postsynaptic NMDAR blockade (fold ΔEPSP_slope_; bath AP5/MK-801 vs. postsynaptic MK-801; 0.68±0.04 vs. 0.73±0.10; n=6,5 cells; p>0.99; Figure 8D,E) tended to be smaller than LTD produced under control conditions (fold ΔEPSP_slope_: 0.42±0.08; n=8 cells; vs. bath AP5/MK-801; p=0.11; vs. postsynaptic MK-801: p<0.05). Nonetheless, control LTD was associated with a comparable increase in PPR to that induced under conditions of postsynaptic NMDAR blockade (ΔPPR; control vs. postsynaptic MK-801: 0.36±0.04 vs. 0.34±0.07; n=8, 5 cells; p>0.99; Figure 8F). In contrast, no PPR changes were observed following unpaired stimulation when NMDARs were blocked by extracellularly (ΔPPR: -0.04±0.05; n=6 cells; vs. control: p<0.01; vs. postsynaptic MK-801: p<0.01 Figure 8F). This suggests that our protocol produced LTD with both pre- and postsynaptic components of expression, and that inhibition of presynaptic NMDAR signalling abolished the presynaptic component of LTD. This is consistent with our imaging experiments, and suggests that presynaptic NMDAR signalling is necessary for the induction of presynaptic LTD. Collectively, our findings provide electrophysiological support to our prior conclusion that presynaptic NMDARs drive decreases in P_r_, independent of the levels of postsynaptic depolarization; across conditions, changes in PPR were always more negative compared to controls when presynaptic NMDAR signalling was inhibited.

Our results demonstrate that presynaptic LTD requires presynaptic NMDAR signalling. Spike-timing dependent (STDP) LTD depends on presynaptic NMDAR activation, as well as endocannabinoid signalling^33,55^. We therefore tested the role of endocannabinoids in the induction of LTD generated by our unpaired stimulation protocol. We found that the endocannabinoid antagonist AM-251 (2μM) failed to prevent decreases P_r_ (Supplemental Figure 10). We also found that presynaptic LTD could be induced even after 1 hour of recording with normal patch electrodes (4-8MΩ), suggesting that it was generally quite resistant to prolonged dilution of postsynaptic factors. LTP, in contrast, failed to be induced by paired stimulation after 10 minutes of recording with a standard patch electrode (Supplemental Figure 11). Our findings suggest that the action of glutamate on the presynaptic NMDAR is sufficient to drive decreases in P_r_, and likely does not require additional postsynaptic factors, such as endocannabinoid signalling.

### A simple learning rule predicts activity-dependent changes in P_r_

Our findings thus far suggest that changes in P_r_ at active presynaptic termianls are driven by two processes: 1) postsynaptic depolarization, which promotes increases in P_r_ through L-VGCC-dependent release of NO from neuronal dendrites, and 2) glutamate release, which promotes decreases in P_r_ through presynaptic NMDAR activation (Figure 9A). We have demonstrated that net changes in P_r_ will depend on both processes. This suggests that changes in P_r_ at a presynaptic terminal are driven by a *mismatch* between 1) the amount of postsynaptic depolarization and 2) the amount of glutamate release that accompanies presynaptic activity. We therefore asked whether these two variables could be incorporated into a mathematical framework that could predict changes in P_r_ (ΔP_r_) in our data set. For ease of calculation, we chose to quantify both these variables in terms of probabilities, that is, 1) the probability that presynaptic activity was accompanied by strong postsynaptic depolarization (P_depol_) and 2) the probability that glutamate was released at the synapse (P_glu_). The simplest mathematical framework to model changes in P_r_ would be: ΔPr=η(P_depol_-P_glu_), where net changes in P_r_ are proportional to the relative difference or *mismatch* between P_depol_ and P_glu_ as defined by (P_depol_-P_glu_), multiplied by some constant η, defined as the learning rate (Figure 9B).

**Figure 9.**
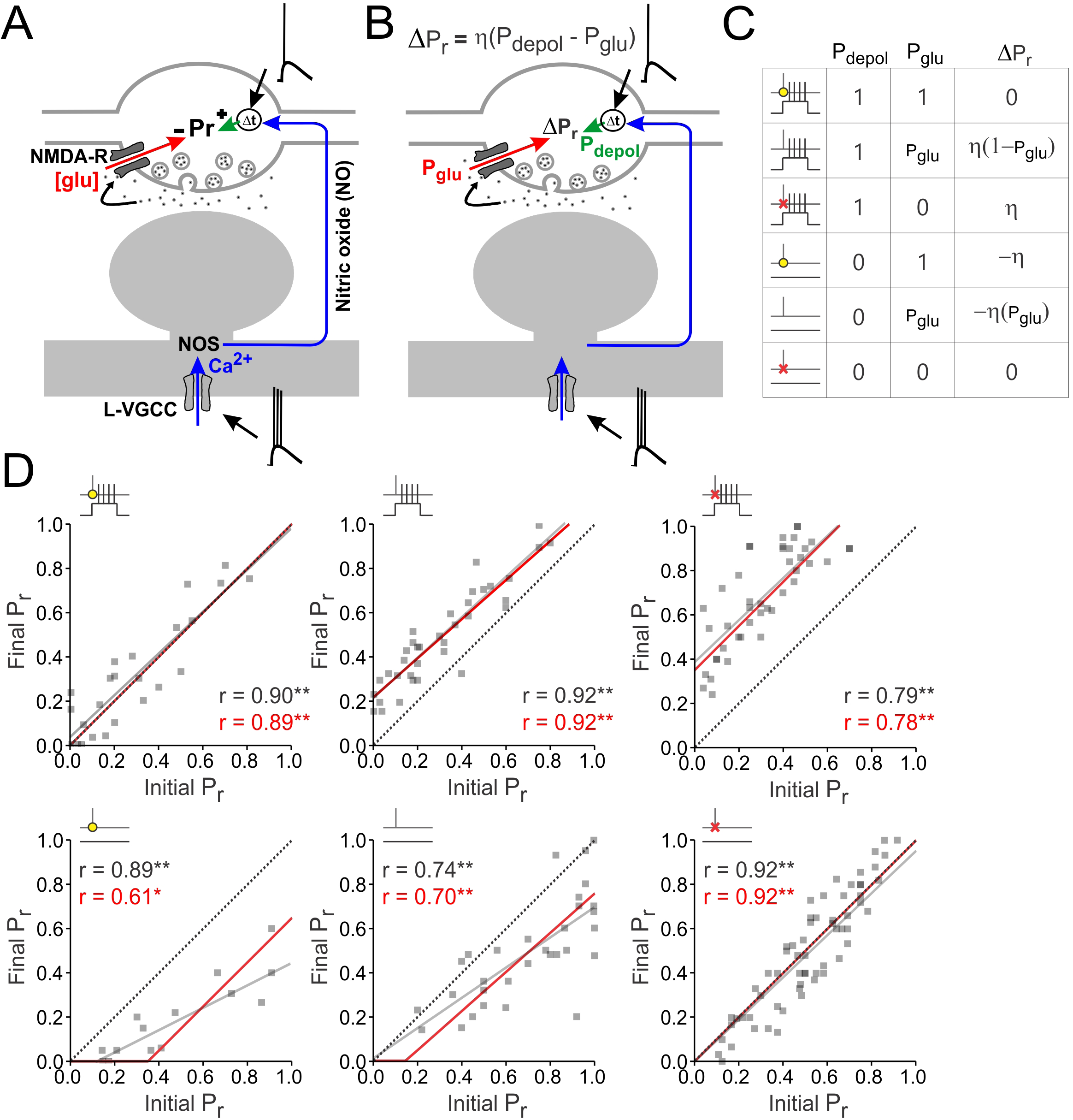
A simple learning rule predicts activity-dependent changes in P_r_. **(A)** Experimental model of presynaptic plasticity. Changes in P_r_ at active presynaptic terminals are determined by two opponent processes. 1) Increases in P_r_ are driven by strong postsynaptic depolarization, which triggers the release of NO from neuronal dendrites. NO synthesis is dependent on NO synthase (NOS), activation of which is triggered by Ca^2+^ influx through L-VGCCs. Importantly, NO drives an increase in P_r_, but only at presynaptic terminals whose activity precedes its release. The detection of such an event requires a detector (Δt) that is sensitive to the relative timings of NO release and presynaptic activity. 2) Decreases in P_r_ are driven by glutamate release ([glu]), via presynaptic NMDAR signalling. Net changes in P_r_ depend on the strengths of both processes, and therefore on the levels of postsynaptic depolarization and glutamate release that accompany presynaptic activity. **(B)** Proposed learning rule of presynaptic plasticity, in which ΔP_r_=η(P_depol_-P_glu_). P_depol_ is the probability that presynaptic activity is causally accompanied by strong postsynaptic depolarization during plasticity induction. P_glu_ is the probability that glutamate was released at the synapse during plasticity induction. η is a constant defined as the learning rate, which was determined to be 0.35 to achieve the best fit. **(C)** Data (n=216 synapses) were divided into 6 categories based on experimental manipulations during plasticity induction in which 60 presynaptic stimuli were delivered at 5Hz. These manipulations include all experimental conditions in which the effects of postsynaptic depolarization were present or absent, whether glutamate photolysis (yellow dot) was present or absent, and whether presynaptic NMDAR blockade (red cross) was present or absent. These categories, along with the associated value of P_depol_, P_glu_, and ΔPr, are summarized in table form. Where P_glu_ did not take the value of 0 or 1, it was calculated using the following formula: P_glu_=0.475*basal P_r_ + 0.2175; see Supplemental Figure 12 for further details. **(D)** For each category, the initial P_r_ is plotted against final P_r_ for all synapses within the category. The broken diagonal line represents the expected trend if P_r_ is unchanged. A line of best fit for the data is shown in grey. The predictions from the learning rule are shown in red. Pearson correlation coefficients (r) calculated for each model are shown, along with their level of significance (*p<0.05, *p<0.01). The learning rule’s predictions are in close agreement with the lines of best fit, except for the fourth category, in which presynaptic stimulation is delivered in the presence of glutamate photolysis but in the absence of postsynaptic depolarization. Although the lines of best fit had greater r values, the learning rule achieved a substantially better (i.e. lower) BIC than the lines of best fit (BIC_line of best fit_ − BIC_learning rule_=34.97). This was also the case when categories having similar trends were combined (category 1 and 6) to minimize the free parameters used by the lines of best fit from 12 to 10 (BIC_line of best fit_ − BIC_learning_ rule=24.25). Thus, the proposed learning rule represents a parsimonious framework that captures the general trend of the data despite only having a single free parameter (η, the learning rate), as compared to a total of 12 free parameters collectively used by the lines of best fits (2 free parameters per line, the slope and the intercept).

We applied the learning rule ΔPr=η(P_depol_-P_glu_) to our data set of imaged synapses across all experimental conditions. As a reminder, to induce plasticity we had delivered 60 presynaptic stimuli at 5Hz, where each stimulus was either consistently paired or unpaired with postsynaptic depolarization. For conditions with paired stimulation, we set Pdepol to 1, since depolarization was present for every presynaptic stimulus. However, for conditions in which NO and L-VGCC signalling was inhibited, P_depol_ was set to 0, as the effects of depolarization would be absent for every presynaptic stimulus. For all other conditions, in which presynaptic stimuli were not paired with presynaptic stimulation, we set P_depol_ to 0. We then determined values for P_glu_. For experiments involving glutamate photolysis, P_glu_ was set to 1, as glutamate was available for every presynaptic stimulus. For experiments in which presynaptic NMDARs were inhibited, P_glu_ was set to 0, as the inhibitory effects of glutamate were absent for every presynaptic stimulus. For all other conditions, P_glu_ had to be calculated for each synapse. Although P_glu_ will likely be proportional to the initial P_r_ of a synapse, it will not be equivalent to initial P_r_ owing to short-term plasticity effects induced during the 5Hz stimulation train^30^. To obtain a more accurate estimate of P_glu_, we conducted an additional set of experiments, in which we used Ca^2+^ imaging to examine glutamate release at single synapses stimulated with 60 presynaptic pulses delivered at 5Hz (Supplemental Figure 12). We calculated P_glu_ as the total number of glutamate release events in the train divided by the number of pulses in the train (i.e. 60). When plotted against initial P_r_, we obtained a linear relationship with which to relate P_glu_ to initial P_r_ (P_glu_=0.475*initial P_r_ + 0.2175; p<0.01; Supplemental Figure 12). We used this relationship to estimate P_glu_ for synapses in the remainder of our data set. Finally, we determined the learning rate η, which obtained the best fit of the model (η=0.35).

To assess the goodness of fit of our learning rule, we divided our data (n=216 synapses) into 6 categories, depending on whether the effects of postsynaptic depolarization were present or absent, whether glutamate photolysis was present or absent, and whether presynaptic NMDAR blockade was present or absent. These 6 categories, along with the learning rule’s predictions for ΔP_r_ are shown in Figure 9C and are summarized in the table below. For a detailed description of the experimental conditions included in each category see the Methods section.

For each category, we plotted initial P_r_ against final P_r_ for all synapses within the category, and superimposed the learning rule’s predictions as a red trend line (Figure 9D). These predictions were compared to lines of best fit (grey lines; Figure 9D) for the data in each category. The lines of best fit took the form of simple linear equations [Final P_r_ = m(Initial P_r_) + b; where m is the slope and b is the y-intercept], which were fit to the data using standard linear regression. The learning rule consistently achieved a predictive power that was comparable to that of the lines of best fit (Figure 9D,E), with the exception of category 4. Synapses in this condition received glutamate photolysis during unpaired stimulation and therefore underwent augmented depression. The model tended to overestimate the amount of depression for synapses with a low initial P_r_ and underestimate the amount of depression for synapses with a high initial P_r_. Although the goodness of fit (as measured by the Pearson correlation coefficient) was greater for the lines of best fit (Figure 9D), our model only had 1 free parameter compared to a total of 12 free parameters used by the lines of best fit across all 6 conditions (2 free parameters per line of best fit, the slope and the intercept). As a means of comparing which model was superior for explaining the data given the number of parameters used in each model, we calculated the Bayesian Information Criterion (BIC) associated with the learning rule and the lines of best fit. The learning rule obtained a substantially better (i.e. lower) BIC value than the lines of best fit (BIC_lines of best fit_ – BIC_learning rule_=34.97; a difference in BIC value of greater than 10 is in favour of the model with the lower BIC value^63^{{). This was also the case when we combined categories having similar trends (category 1 and 6) to minimize the free parameters used by the lines of best fit from 12 to 10 (BIC_lines of best fit_ – BIClearning rule=24.25). Thus, the proposed learning rule represents a simple framework that is capable of effectively predicting changes in P_r_ across a range of experimental conditions.

**Table 1.**
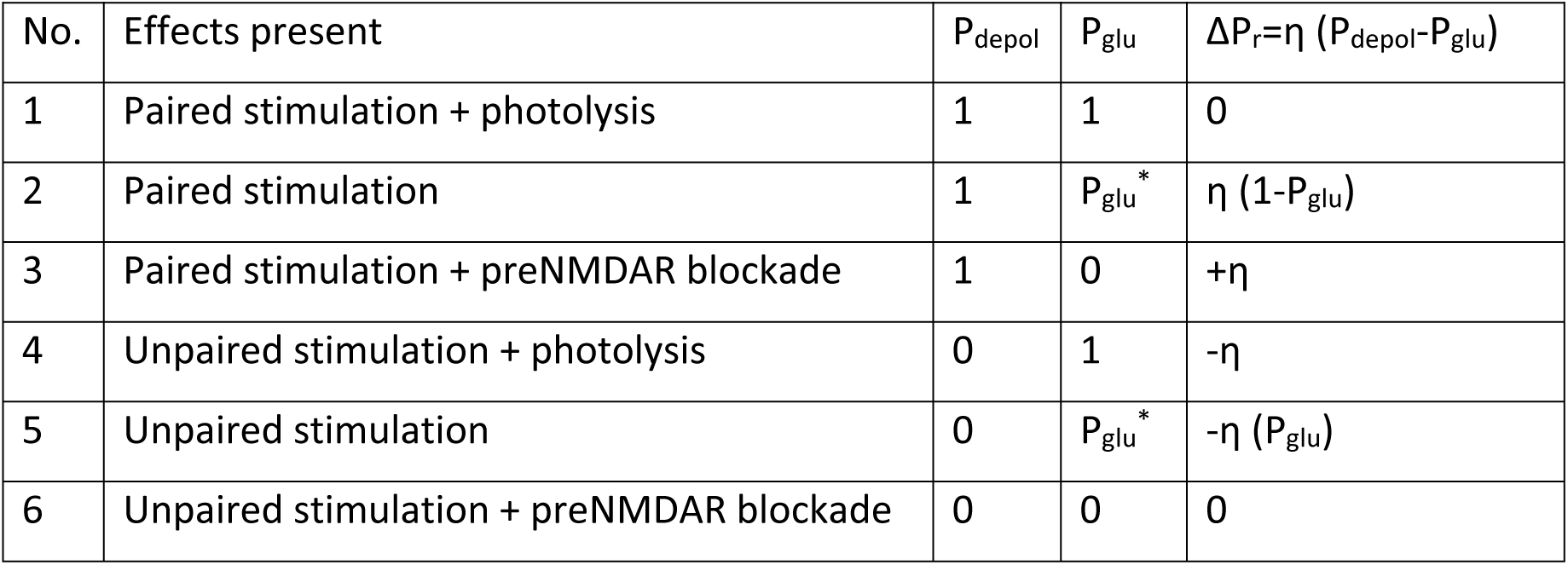
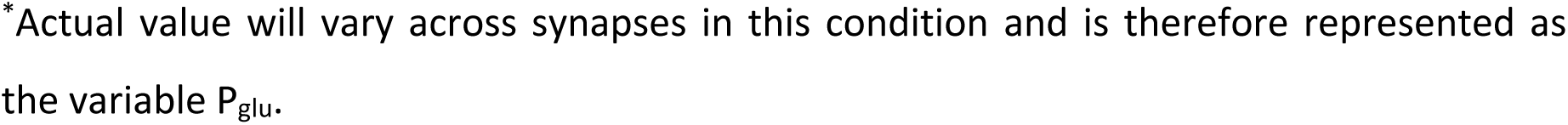
Summary of experimental conditions and model parameters.

## Discussion

We have explored the mechanisms of presynaptic plasticity at CA3-CA1 synapses in hippocampal slices. Based on our findings we present a unified framework of presynaptic plasticity, which suggests that, at active presynaptic terminals, changes in P_r_ are driven by two processes: 1) postsynaptic depolarization, which promotes increases in P_r_ through the L-VGCC-dependent release of NO from neuronal dendrites (LTP-promoting process), and 2) glutamate release, which promotes decreases in P_r_ through presynaptic NMDAR activation (LTD-promoting process). Both processes operate together to tune presynaptic function, with net changes in P_r_ depending on the strength of each process during presynaptic activity. (Figure 9A,B). We also found that this model could be formalized as a simple learning rule [ΔP_r_=η(P_depol_-P_glu_)] that was able to accurately predict changes in P_r_ in our data given the probability that presynaptic activity was associated with strong postsynaptic depolarization (P_depol_) and the probability presynaptic activity was associated with glutamate release (P_glu_).

### Presynaptic LTP can occur in the absence of synapse-specific glutamate signalling

Importantly, the mechanism of plasticity proposed in this study enables presynaptic terminals releasing little or no glutamate to become potentiated provided that their activity is accompanied by strong postsynaptic depolarization. This feature of presynaptic LTP may have important implications for plasticity. Most central synapses have low glutamate release probabilities, with some synapses appearing to release no glutamate in response to presynaptic stimulation^64,65^. This is true for synapses recorded in both *in vitro* preparations from young rodents and *ex vivo* preparations from adult rodents. In fact, under electron microscopy, a significant portion of synapses (up to 35-50%) in the adult rodent hippocampus have presynaptic zones lacking synaptic vesicles in their near proximity (<170nm); these so-called “nascent zones” have been hypothesized to be functionally silent^66^. Although the existence of *bona fide* presynaptically silent synapses remains controversial^64^, the low release probabilities (average P_r_ of approximately 0.2^67^) of central synapses suggests that it is possible that activity at a presynaptic terminal may not elicit glutamate release at the synapse, but may still coincide with strong postsynaptic depolarization, driven by glutamate release at other co-active synapses. Under such conditions the presynaptic mechanisms of plasticity elucidated in this study, could enable the efficient induction of Hebbian potentiation presynaptically.

Our finding that presynaptic enhancements can occur without glutamatergic signalling at the synapse raises the question as to why many studies show that LTP induction can be abolished or impaired by blockade of one or more glutamate receptor subtypes^59,68,69^. To address this question, it is first important to recognize that not all LTP induction protocols are associated with presynaptic enhancements^6^. This is because presynaptic LTP requires higher levels of postsynaptic depolarization than postsynaptic LTP^6,17,18^. This likely reflects the necessity of L-VGCC signalling in presynaptic potentiation^16-19^, which have high voltage activation thresholds^6^. Thus, successful induction of presynaptic LTP will depend on the levels of postsynaptic depolarization achieved by a given stimulation protocol^16,18^, which in turn will be influenced by a variety of experimental factors, including the frequency and intensity of stimulation. L-VGCCs also exhibit voltage-dependent inactivation during extended periods of depolarization (>1s)^70,71^. As such, the use of pairing protocols, in which the postsynaptic cell is voltage clamped at 0mV for prolonged periods of time (>10s) during presynaptic stimulation, are likely to be unsuitable for driving presynaptic enhancements, despite being commonly used to drive postsynaptic enhancements ^6^.

Whether presynaptic enhancements are obtained in the first place will therefore depend on the levels of postsynaptic depolarization achieved during LTP induction. Nonetheless, even studies reporting presynaptic LTP also find that inhibition of glutamate receptors, in particular NMDARs, abolish or reduce presynaptic enhancements^11-15^. It is however important to recognize that AMPARs, KARs, NMDARs, and mGluRs all have been shown to contribute to postsynaptic depolarization^40,45-47^. Given that presynaptic changes rely on the voltage-dependent release of NO, it is possible that blockade of any of these glutamate receptor classes would abolish or reduce presynaptic LTP in an indirect way, by reducing postsynaptic depolarization and the activation of L-VGCCs. This may explain, in part, why experimental manipulations that augment the levels of postsynaptic depolarization reliably rescue LTP in AMPAR^59^,^72^, NMDAR^6,14,17-19,73-76^, and mGluR blockade^77^. Importantly, our LTP induction protocol used strong postsynaptic depolarization, which was elicited by somatic current injection, and therefore independent of synaptic activity. This circumvented the need for any glutamate receptor-dependent depolarization during paired stimulation and enabled us to directly assess the function of glutamate signalling in presynaptic LTP, independent of its effects on postsynaptic depolarization. Based on these results, we would argue that the physiological role of glutamate release in presynaptic potentiation is for driving postsynaptic spiking as opposed to conveying a synapse-specific signal; this contrasts with the role of glutamate release in postsynaptic plasticity, in which synapse-specific activation of postsynaptic NMDARs is necessary for LTP induction.

While our approach for inducing LTP resembles that of traditional STDP protocols, which rely on NMDAR activation^78^, there is are two key differences. Firstly, in our study, postsynaptic depolarization took the form of complex spikes, which included a brief period (7-10ms) depolarization before the first spike. This period of subthreshold depolarization is known to facilitate the induction of LTP, possibly by inactivating voltage-gated potassium channels within the dendrite, which otherwise impede action potential backpropagation^79-84^. Moreover, like complex spikes recorded in vivo^39^, the spike trains we triggered contained broadened action potentials, which likely reflect strong depolarization in the dendrites^81,83^. Consequently, the postsynaptic waveforms used in our study were likely to generate stronger levels of postsynaptic depolarization, and in a manner independent of glutamate release and NMDAR activation, than those used in STDP studies.

Secondly, our study differed from others in the method used to selectively block postsynaptic NMDARs. Traditionally, postsynaptic NMDAR blockade is achieved by recording from neurons with a high concentration of MK-801 (0.5-1mM) in the patch electrode^31,54,55^. The use of such high concentrations of drug, which is several orders of magnitude greater than the K_d_ value with which MK-801 binds the NMDARs^60^, is likely important for achieving a rapid and effective block of NMDARs in diffusionally isolated compartments, such as narrow dendrites and spines distal from the soma. Lower concentrations of intracellular MK-801 (1μM)) have been successfully used to block postsynaptic NMDAR, but presumably require a longer time to take effect^85^, and would therefore not be conducive for plasticity experiments, in which the time of patch recordings needs to be minimized to avoid dilution of postsynaptic factors. The use of high concentrations of MK-801 during intracellular loading, however, comes at the potential cost of off-target effects^61,62^. Indeed, we found in our study that within 5 minutes of patching on to a neuron with 1mM MK-801, L-VGCC Ca^2+^ transients were reduced by about 50%. Given the necessity of L-VGCCs in the induction of presynaptic LTP, the use of high intracellular concentrations of MK-801 presents a serious confound in plasticity experiments. In our study, we found that the ideal method of MK-801 was bolus-loading. This entailed transiently patching the cell with a high concentration of MK-801 (5mM) for a short period of time (60 seconds). After about 20 minutes the drug effectively abolished NMDAR signalling at recorded synapses, which were within 100μm of the soma, without effecting L-VGCC Ca^2+^ transients. The cell could then be re-patched with normal internal solution if required. Using this protocol, paired stimulation reliably produced LTP, which was otherwise abolished when 1mM of MK-801 was instead included in the patch pipette from the start of recording.

### Presynaptic LTP requires nitric oxide signalling

It has long been recognized that the induction of LTP at the presynaptic locus requires a retrograde signal^86^. The most promising candidate is NO^8^. The role of NO in LTP has been a source of much controversy, and some studies have concluded that NO signalling is not necessary in LTP induction^8^. However, given that NO is likely to be important for presynaptic strengthening, the effect of NO signalling on synaptic plasticity will depend on whether presynaptic enhancements are obtained following LTP induction^6^. Indeed, studies that actually confirm presynaptic changes following LTP induction, including our own, consistently demonstrate that presynaptic enhancements depend on the synthesis and release of NO in both acute and cultured hippocampal preparations^12,87-89^.

It has generally been assumed that NO synthesis is dependent on Ca^2+^ influx from postsynaptic NMDARs^8^; however, several studies have demonstrated that induction of presynaptic LTP is possible in NMDAR blockade suggests that this signalling pathway is not necessary for presynaptic potentiation^17-19^. Consistent with this notion, in our experiments, we found that specific blockade of postsynaptic NMDARs using intracellular application of MK-801 had no effect on changes in P_r_ induced by paired stimulation. Instead, we found an alternative pathway for NO synthesis that was crucial for presynaptic strengthening, and that depended on strong postsynaptic depolarization and activation of L-VGCCs; this is in keeping with findings from a recent study{Pigott, 2016 #8896}. The differential importance of NO signalling mediated by NMDARs and L-VGCCs in presynaptic LTP may result from differences in the magnitude, kinetics, and/or spatial extent of NO signalling associated with the activation of each channel. Unfortunately, the poor sensitivity of NO-indicator dyes makes this possibility difficult to investigate with currently available tools.

It has previously been demonstrated that exogenous NO can potentiate synaptic transmission, and that this potentiation is restricted to synapses that are active during NO release^90,91^. Here, we extend these findings by showing that photolysis of NO at single synapses can actually drive increases in Pr, and that this increase can occur in the absence of glutamatergic signalling. Moreover, we demonstrate that the potentiating effects of NO are not only restricted to active synapses, but specifically at synapses whose activity precede, rather than follow, NO release; thus, the requirements of NO signalling are consistent with those of Hebbian and spike-timing dependent plasticity^78^. It is important to mention that some groups have found no effect of exogenous NO application on synaptic plasticity^6,8^. However, like glutamate, the effects of NO will depend on the spatiotemporal dynamics of NO signalling and the pattern of concurrent synaptic activity, which will likely vary across studies; therefore it is not surprising that NO application, like that of glutamate, can potentiate, depress, or have no effect on synaptic input depending on experimental conditions^6^.

Although our study has focussed on phasic NO signalling, it is important to mention that LTP is additionally likely to require a tonic, low-level of NO signalling^92^. In our study, we used cPTIO to scavenge NO. It should be noted that cPTIO reacts with NO with a low rate constant, and therefore would preferentially block tonic as opposed to phasic NO signalling ^93^.

### A presynaptic co-incident detector for Hebbian activity

Presynaptic LTP does not require synapse-specific glutamate signalling but still requires presynaptic activity to precede strong postsynaptic depolarization. This means that the induction of presynaptic LTP is Hebbian, but the detector of causal pre- and postsynaptic activity must not directly rely on glutamatergic signalling, and therefore cannot be the postsynaptic NMDAR. This suggests that another co-incident detector must be present at the synapse. Given that we could induce presynaptic LTP by causally pairing presynaptic activity with NO release, it would suggest that the likely Hebbian integrator for presynaptic LTP should be sensitive to 1) presynaptic activity, which could be potentially signalled by action potential-triggered Ca^2+^ influx via presynaptic VGCCs and 2) NO release. One candidate integrator for this purpose might be guanylate cyclase, which is a protein that has previously been implicated in the induction of presynaptic LTP^94,95^ and is also known to be modulated by both NO and intracellular Ca^2+^ ^96^. In fact, in platelets and in lung endothelial cells, Ca^2+^ influx appears to enhance the NO-sensitivity of guanylate cyclase by re-locating the protein to the lipid membrane, where NO is likely to be most concentrated^96^. Such a mechanism, if present at, presynaptic terminals, would explain the timing-sensitivity of presynaptic LTP. NO release from postsynaptic depolarization would be maximally detected if guanylate cyclase was present at the membrane shortly before, but not after, postsynaptic depolarization. This would mean that if relocation of guanylate cyclase to the membrane could be triggered by presynaptic activity, via VGCC-mediated Ca^2+^ influx, then maximal NO detection would require presynaptic activity to precede postsynaptic depolarization.

### Glutamate drives presynaptic LTD by activating presynaptic NMDARs

At active presynaptic terminals, whereas postsynaptic depolarization drives increases in P_r_, we show, unexpectedly, that glutamate release drives decreases in P_r_ by acting on presynaptic NMDARs. We found that presynaptic NMDAR signalling operated both during LTP and LTD induction paradigms to reduce P_r_. This was true regardless of whether NMDARs were blocked with bath application of AP5 or MK-801, suggesting that the inhibitory effects of presynaptic NMDAR signalling on P_r_ were mediated by ionotropic rather than metabotropic functions of the channel^97^. Importantly, our finding suggests that the potentiating effects of postsynaptic depolarization and the depressing effects of endogenous glutamate release occur simultaneously during synaptic activity, regardless of the nature of postsynaptic depolarization. Thus, the processes underlying LTP and LTD induction are not temporally distinct mechanisms as traditionally believed, but operate jointly to tune synaptic function during presynaptic activity. Our results may explain why sometimes the same pairing protocol that produces LTP at low Pr synapses, produces LTD at high P_r_ synapses; presumably the postsynaptic depolarization achieved by such protocols is not of sufficient magnitude to prevent the depressing effects of glutamate release at high P_r_ synapses^28,29^. Our results also may explain why the locus of LTP expression, whether pre- or postsynaptic, appears to depend on initial P_r_^27^. With higher basal release probabilities, more glutamate is released for a given LTP induction protocol, meaning that postsynaptic LTP is favoured owing to greater postsynaptic NMDAR-signalling, whereas presynaptic LTP is inhibited owing to greater presynaptic NMDAR-signalling. Thus low P_r_ synapses will have a tendency to express LTP presynaptically while high P_r_ synapses have a tendency to express LTP postsynaptically^27^.

In contrast to our findings, pharmacological inhibition of presynaptic NMDARs at neocortical synapses does not appear to effect LTP^32,53,54^. It is possible that the low frequency (0.2Hz) of presynaptic stimulation used during LTP induction in these studies did not result in sufficient levels of glutamate release to elicit presynaptic depression via presynaptic NMDAR activation. By contrast, in our study LTP induction involved presynaptic stimulation at a theta frequency, which is effective at promoting glutamate release at Schaffer-collateral synapses^30^. As such, the inhibitory effects of presynaptic NMDARs on LTP may only be evident at higher stimulation frequencies.

Presynaptic NMDARs may, however, operate at lower stimulation frequencies in the context of spike-timing dependent plasticity. Recently, it was shown that low-frequency (0.2Hz), anti-causal pairing of pre- and postsynaptic spiking induced presynaptic LTD at hippocampal synapses, in a manner dependent on presynaptic NMDAR signalling^31^. This form of LTD, in addition to glutamate release, required endocannabinoid and astrocytic signalling, similar to spike-timing dependent LTD in the neocortex^33^. Incidentally, inhibition of endocannabinoid signalling does enhance the levels of presynaptic LTP at neocortical synapses induced by pairing pre- and postsynaptic activity, suggesting that the mechanisms underlying this form of LTD may also be engaged during LTP induction^98^. The LTD in our study, however, did not depend on endocannabinoid signalling and resembled the presynaptic NMDAR-mediated self-depression of presynaptic function that is present at developing neocortical synapses^53^. At these synapses, this form of LTD is dependent on the downstream activation of the Ca^2+^-sensitive phosphatase, calcineurin^53^.

Previously we have demonstrated that presynaptic NMDARs at hippocampal synapses facilitate transmitter release during theta stimulation^30^. Given our current findings, presynaptic NMDARs appear to be important for presynaptic facilitation in the short-term, but presynaptic depression in the long-term. This is consistent with the finding that presynaptic NMDARs in the neocortex also mediate short-term plasticity of glutamate release, and yet are similarly implicated in presynaptic LTD^32,33,56^. It may appear peculiar for a single protein to mediate seemingly disparate functions; however, another way to view the presynaptic NMDAR is as a dynamic regulator of presynaptic activity, appropriately tuning glutamate release depending on the patterns of pre- and postsynaptic activity. As such, the receptor may aid glutamate release during theta-related activity, but, triggers presynaptic LTD when this release fails to elicit sufficiently strong levels of postsynaptic depolarization.

### Presynaptic plasticity optimally tunes glutamate release

We found a simple mathematical framework [ΔPr=η(P_depol_-P_glu_)] could predict changes in P_r_ based on a mismatch between 1) the probability that presynaptic activity is accompanied by strong depolarization (P_depol_) and 2) the probability that presynaptic activity is accompanied by glutamate release (P_glu_) during synaptic stimulation. According to this model, presynaptic plasticity is driven only when there is a mismatch between P_glu_ and P_depol_. Plasticity, by changing P_r_, works to drive the value of P_glu_ closer to that of P_depol_. This has a very intuitive interpretation. P_glu_ is a measure of the synapse’s ability to drive postsynaptic activity. P_depol_ is a measure of the synapse’s ability to predict postsynaptic activity. By driving the value of P_glu_ closer to P_depol_, presynaptic plasticity changes a synapse’s ability to drive postsynaptic activity, so that it matches its ability to predict postsynaptic activity.

It is important to recognize that P_glu_ will not only depend on basal P_r_, but also on the pattern of presynaptic activity, owing to the effects of short-term plasticity^99^. Moreover, for any given pattern of pre- and postsynaptic activity, there exists a value of P_r_ for which P_glu_ will equal P_depol_. That is, there exists a value of P_r_ for which the probability of glutamate release for a given pattern of presynaptic stimulation (P_glu_) will be equal to the probability that presynaptic stimulation is accompanied by strong postsynaptic depolarization (P_depol_). The model predicts that this value of P_r_ represents a target value that all synapses will tend to for a given pattern of pre- and postsynaptic activity. Synapses below this target value will be potentiated because P_depol_>P_glu_, whereas those above this target value will be depressed because P_depol_<P_glu_. This can explain why in the literature, P_r_ has previously been reported to tend to a certain value when a particular pattern of pre- and postsynaptic stimulation is applied^28^.

According to our model, the target P_r_ value given by pairing high frequency presynaptic stimulation with strong postsynaptic depolarization is lower than pairing low frequency presynaptic stimulation with strong postsynaptic depolarization. In both cases, P_depol_ is high, meaning that presynaptic plasticity will continue to drive P_r_ until Pglu is also high. However, with high frequency stimulation, a low target P_r_ achieves a high P_glu_ owing to the effects of short-term plasticity. In contrast, with lower frequency stimulation, a higher target P_r_ is required to achieve a similarly high P_glu_. Consequently, as we showed in our experiments with Figure 1, when paired with strong postsynaptic depolarization, low frequency presynaptic stimulation will produce greater presynaptic potentiation than higher frequency stimulation. That P_r_ is kept low at synapses where presynaptic bursts are good predictors of postsynaptic spiking ensures that only presynaptic bursts mobilize glutamate, rather than individual presynaptic stimuli, which may not be good predictors of postsynaptic spiking. Therefore, presynaptic plasticity appears to tune P_r_, such that the patterns of presynaptic activity that predict postsynaptic spiking are the ones that most efficiently drive glutamate release.

### Presynaptic plasticity corrects prediction-errors that arise during synaptic activity

According to our learning rule, the role of presynaptic plasticity is to ensure that the ability for a presynaptic terminal to release glutamate is matched with its ability to predict postsynaptic spiking. The learning rule could also be interpreted from a prediction-error framework. Prediction error is a highly influential learning theory that has been applied widely in systems-level neuroscience^100^. When viewed through the prediction-error framework, our model [ΔPr=η(P_depol_-P_glu_)] is essentially a form of the Rescorla-Wagner learning rule [Learning = η(Prediciton-Outcome); where η is the learning rate], in which learning is driven by mismatches between prediction and outcome^101^. Accordingly, the probability that glutamate will be released by an active presynaptic terminal (P_glu_) can be viewed as a prediction of the probability that strong postsynaptic depolarization (P_depol_) that is to follow. For example, activity at synapses releasing little glutamate is not expected to be accompanied by strong postsynaptic depolarization. A mismatch between the predicted and actual probabilities of postsynaptic depolarization (P_depol_≠P_glu_), such as that which occurs when activity at a synapse releasing low levels of glutamate is unexpectedly accompanied by strong postsynaptic depolarization, results in a change in P_r_ which by affecting glutamate release, improves the accuracy of future prediction. Synaptic plasticity, therefore, would be preferentially driven by novel patterns of neuronal activity, as these will generate the largest prediction errors. Such a mechanism may provide a synaptic-level explanation as to why learning, at the behavioral level, preferentially occurs in novel and unexpected situations.

## Methods

### Cultured hippocampal slices

All animal work was carried out in accordance with the Animals (Scientific Procedures) Act, 1986 (UK). Hippocampal slice cultures were used for most experiments owing to the excellent optical and electrophysiological access to cells and synapses afforded by the preparation. Cultured hippocampal slices (350μm) were prepared from male Wistar rats (P7-P8), as previously described^13^. Slices were maintained in media at 37^°^C and 5% CO2 for 7-14 days prior to use. Media comprised of 50% Minimum Essential Media, 25% heat-inactivated horse serum, 23% Earl’s Balanced Salt Solution, and 2% B-27 with added glucose (6.5g/L), and was replaced every 2-3 days. During experimentation, slices were perfused with artificial cerebrospinal fluid (ACSF; 1mL/min), which was constantly bubbled with carbogen (95% O2 and 5% CO2) and heated to achieve near-physiological temperatures in the bath (31-33^°^C). ACSF contained (in mM) 145 NaCl, 16 NaHCO_3_, 11 glucose, 2.5 KCl, 2-3 CaCl_2_, 1-2 MgCl_2_, 1.2 NaH_2_PO_4_, and, to minimize photodynamic damage, 0.2 ascorbic acid and 1 Trolox.

### Acute hippocampal slices

Acute hippocampal slices were used to confirm key findings in cultured hippocampal slices. When this preparation was used, it is clearly stated in the text and figure captions. Coronal acute hippocampal slices (400μm) were prepared from 2-4 week old male Wistar rats. Tissue was dissected in a sucrose-based ACSF solution (inmM: 85 NaCl, 65 sucrose, 26 NaHCO_3_, 10 glucose, 7 MgCl_2_, 2.5 KCl, 1.2 NaH_2_PO_4_, and 0.5 CaCl_2_). The whole brain was sliced into coronal sections using a Microm HM 650V vibratome (Thermo Scientific) and hippocampi were carefully dissected from each slice. Hippocampal tissue were allowed to recover at room temperature in normal ACSF (120 NaCl, 2.5 KCl, 2 CaCl_2_, 1 MgCl_2_, 1.2 NaH_2_PO_4_, 26 NaHCO_3_, and 11 glucose), which was bubbled with 95% O2 and 5% CO2. Slices were given at least 1 hour to recover before use. During experimentation, slices were perfused with ACSF (3mL/min) containing picrotoxin (100μM). The ACSF was constantly bubbled with carbogen (95% O2 and 5% CO2) and heated to achieve near-physiological temperatures in the bath (31-33^°^C).

### Electrophysiological Recordings

CA1 pyramidal neurons were recorded from either using low (4-8MΩ) or high resistance patch electrodes (18-25MΩ) filled with standard internal solution (in mM: 135 KGluconate, 10 KCl, 10 HEPES, 2 MgCl_2_, 2 Na_2_ATP and 0.4 Na_3_GTP), or sharp microelectrodes (80-120MΩ) filled with 400mM KGluconate. For some experiments, low resistance (4-8MΩ) patch electrodes were filled with ATP regenerating solution (in mM: 130 KGluconate, 10 KCl, 10 HEPES, 10 NaPhosphocreatine, 4 MgATP, 0.4 Na_3_GTP and 50U/mL creatine phosphokinase) ^73^. The recording method used in a given experiment is clearly indicated in the text.

### Stimulation protocols

A glass electrode (4-8MΩ), filled with ACSF, was placed in stratum radiatum. Continuous basal stimulation (0.05-0.10Hz) was present for all experiments, and was only interrupted to deliver paired-pulse or tetanic stimulation. Stimulation intensity was adjusted to evoke a 5-10mV EPSP; pulse duration was set at 100μs. Paired-pulse stimulation consisted of 2 presynaptic stimuli delivered 70ms apart. LTP induction consisted of 60 single pulses delivered at 5Hz each paired with postsynaptic depolarization. Postsynaptic depolarization took the form of a complex spike. To emulate a complex spike, we injected a postsynaptic current waveform (2-3nA) that was approximately 60ms in duration and resulted in 3-6 spikes at ∼100Hz, with the first spike occurring 7-10ms after the presynaptic stimulus. This was done by injecting a (2-3nA) waveform with a 7-10ms rising phase, 20ms plateau phase, 30-33ms falling phase. However, in experiments shown in main Figure 2, the current waveform took the form of a 50ms flat current step; though this led to poorer control over the start of postsynaptic bursting. Stimulating electrodes were placed within 50-70μm of the soma to ensure that postsynaptic depolarization reached stimulated synapses without significant attenuation. LTD induction consisted of either 60 single or paired (inter-stimulus interval of 5ms) presynaptic pulses delivered at 5Hz in the absence of postsynaptic depolarization. For induction of LTD with paired pulses, the membrane was hyperpolarized (-90 mV) to prevent somatic and dendritic spiking. For induction of LTD with single pulses, the postsynaptic neuron was hyperpolarized only in some cases since stimulation rarely generated postsynaptic spikes.

### Electrophysiology and analysis

All electrophysiological data was recorded using WinWCP (Strathclyde Electrophysiology Software) and analyzed using Clampfit (Axon Insturments) and Excel (Microsoft). The initial EPSP slope, calculated during the first 3ms of the response, was used to analyze changes in the EPSP throughout the recording. This was done to ensure only the monosynaptic component of the response was analyzed. All data was normalized to the average EPSP slope recorded during baseline to yield ΔEPSP slope. Paired pulse ratio (PPR) was calculated as the average EPSP slope evoked by the second stimulation pulse divided by the average EPSP slope evoked by the first stimulation pulse, as previously described ^102^; averages were calculated from 10-20 paired pulse trials. Decreases in PPR are thought to reflect increases in release probability^103^. The coefficient of variation parameter CV^-2^, which reflects the mean^2^/variance, was calculated using the EPSP slopes collected over 25-30 trials. The CV^-2^ calculated from 25-30 stimulation trials taken 30 minutes following LTP induction was normalized to the CV^-2^ calculated from 25-30 stimulation trials at baseline to yield ΔCV^-2^. Increases in CV^-2^ reflect possible increases in P_r_^104^. To control for the effects of incomplete drug washout, for experiments with glutamate receptor blockade, ΔEPSP slope and ΔCV^-2^ for the tetanized pathway were compared to values obtained for the control pathway.

### Bolus-loading

Bolus-loading was used to fill CA1 neurons with dye or drugs whilst minimizing the amount of time the cell was patched on to. Loading was achieved by transiently patching onto cells (60 seconds) using low-resistance patch electrodes (4-8MΩ) containing a high-concentration of drug or dye dissolved in standard internal solution. Slow withdrawal of the patch using a piezoelectric drive ensured re-sealing with no observable adverse effects to cell health. Cells were then subsequently re-patched for the purposes of delivering postsynaptic depolarization if and when required.

### Ca^2+^ imaging and analysis

Cells were bolus-loaded with OGB-1 (0.5-1mM), except in instances of sharp microelectrode recordings, in which dye (0.5mM OGB-1) was present in the microelectrode (80-120MΩ). A stimulating glass electrode (4-8MΩ) was then brought near (5-20μm) to a branch of imaged dendrite within stratum radiatum. For visualization purposes, electrode tips were coated with bovine serum albumin Alexa Fluor 488 conjugate (Invitrogen), as previously described^105^. Briefly, a 0.05% BSA-Alexa 488 solution was made with 0.1M phosphate-buffered saline containing 3mM NaN3. Pipette tips were placed in the solution for 2-5 minutes.

Ca^2+^ transients were evoked with pairs of stimuli (70ms apart) and monitored by restricting confocal laser scanning to a single line through the spine and underlying dendrite; images were acquired at a rate of 500Hz and analyzed using ImageJ and Microsoft excel. Increases in fluorescence intensity (ΔF/F=F_transient_−F_baseline_/F_baseline_) following the delivery of the first stimulus reflect successful glutamate release from the presynaptic terminal ^13,44^. The proportion of successful fluorescent responses to the first stimulus across stimulation trials was used to calculate P_r_. P_r_ was assessed on the basis of 15-40 trials at baseline and at 25-30 minutes post-tetanus. For high P_r_ synapses (>0.8) the number of stimulation trials was limited to 15-20 to avoid photodynamic damage that results from imaging the frequent Ca^2+^ responses generated at these synapses. For all other synapses, P_r_ was generally assessed using 20-35 trials of stimulation. Synapses with initial P_r_ values of 0-0.7 were used for LTP experiments. Since our LTD protocol did not elicit depression in low P_r_ synapses (Figure 2), for the majority of LTD experiments, synapses with P_r_ values of 0.5-1.0 were used. In experiments involving glutamate receptor blockade, P_r_ was measured prior to drug application at baseline, and measured post-tetanus, following drug washout. In experiments involving NMDAR blockade, using either AP5 or MK-801, drugs were present for the duration of the experiment and, therefore, present for both the baseline and post-tetanus measurements of P_r_.

### Nitric oxide imaging

Experiments involving DAF-FM (Invitrogen) imaging were carried out in Tyrodes buffer (inmM: 120 NaCl, 2.5 KCl, 30 glucose, 4 CaCl_2_, 0 MgCl_2_, and 25 HEPES) containing 50μM D-AP5, 10μM NBQX, 500μM MCPG, and 100μM LY341495 (Abcam) to block glutamate receptors, as well as 1μM Bay K-8644 (Abcam) to prevent L-VGCC desensitization during K^+^ application as previously described^49,50^. CA1 pyramidal neurons were bolus-loaded with 250μM of DAF-FM. Apical dendrites, often secondary or tertiary branches, within 100μm of the soma were imaged at one focal plane, once prior to, and once 5-10s following, the addition of a high K^+^ Tyrodes solution (inmM: 32.5 NaCl, 90 KCl, 30 glucose, 4 CaCl_2_, 0 MgCl_2_, 25 HEPES, 50μM D-AP5, 10μM NBQX, 500μM MCPG, and 100μM LY341495). Laser power and exposure was kept to a minimum to avoid photobleaching. In our hands, DAF-FM basal fluorescence was not quenched by intracellular addition of cPTIO.

1,2-Diaminoanthraquinone (DAQ; Sigma) was used to image activity-dependent NO release under more physiological conditions. DAQ was loaded as previously described^51^. DAQ was prepared as a 5mg/mL stock solution dissolved in DMSO. Hippocampal slices cultures were treated with 100μg/mL of the solution for 2 hours at 37^°^C and 5% CO2. Slices were then placed on the rig, and washed heated and oxygenated ACSF for 30 minutes prior to imaging. DAQ was imaged in full glutamate receptor blockade using 488nm excitation light and a 570nm long-pass emission filter prior to and following stimulation of a single patched CA1 neuron with 600 complex spikes at 5Hz (see Stimulation protocols section). Control cells were left unstimulated. Following DAQ imaging, cells were re-patched, loaded with Alexa Fluor 488 (100μm; Invitrogen), and imaged. Alexa Fluor 488 fluorescence was used to determine the proportion of imaged DAQ fluorescence that co-localized to the recorded cell.

### Photolysis

A 405nm laser (Photonics) was used for spot photolysis. The laser was focussed to a small spot (∼1.2μm diameter) by overfilling the back aperture of a 60x water-immersion lens (Olympus). Electrode manipulators and recording chambers were mounted on a movable stage, which enabled a region above the spine head to be positioned beneath the photolysis spot. Laser exposure was controlled using a fast shutter (LS6; Uniblitz). For glutamate photolysis, MNI glutamate (Tocris) was focally delivered through a glass pipette (4-8MΩ; 10mM MNI glutamate) using a picospritzer (Science Products). Laser exposure was limited to ∼2ms and, in each experiment, the laser intensity (0.5-2mW) was adjusted to generate a Ca^2+^ response in the underlying spine that was comparable to the response generated by electrical stimulation. For NO photolysis we used ruthenium nitrosyl chlorides (RuNOCl3) which are known to have sub-millisecond release kinetics^106^. For spot photolysis, 0.5-1mM RuNOCl_3_ (Sigma) was bath applied and uncaged using 30-60 laser pulses (25ms; 2mW) delivered at 5Hz; presynaptic stimulation either preceded or followed NO photolysis by 10ms. Using the NO-indicator, DAF-FM (Invitrogen), we calibrated laser power to liberate approximately 10nM of NO per pulse. We did this by targeting the soma of DAF-FM loaded neurons for photolysis at different laser powers while recording the resulting increases in fluorescence using the confocal laser in line scan mode (500Hz). We aimed for an increase in fluorescence of about 6-7% (averaged across several trials), which based on the manufacturer’s data on the concentration-dependent fluorescence of DAF-FM, amounts to a release of approximately 10nM of NO. For wide-field UV photolysis, 100μM RuNOCl3 was added to the patch electrode and allowed to diffuse into the cell for 10-15 minutes prior to commencing the experiment. A UV Flash Lamp (HI-TECH Scientific; 100V) was used to deliver a 1ms wide-field uncaging pulse that was timed to occur 7-10ms before or after presynaptic stimulation. Because of the time required for the UV lamp to recharge between flashes, about 20 of the 60 presynaptic pulses delivered at 5Hz were not associated with a flash.

### Pharmacology

Glutamate receptor blockade was achieved using D-AP5 (50-100μM; Abcam), NBQX (10μM; Abcam), R,S-MCPG (500μM; Abcam) and LY341495 (100μM; Abcam). In experiments requiring both pre- and postsynaptic NMDARs to be blocked, either D-AP5 (50-100μM) or MK-801 (20μM) was added in bath for the duration of the experiment. In the case of MK-801, slices were pre-incubated with the drug for at least 1 hour prior to experimentation. Experiments in which NMDARs were blocked with bath application of AP5 and with bath application of MK-801 produced similar results and so conditions were combined for data analysis. Postsynaptic NMDARs were blocked by bolus-loading of 5mM MK-801 (Abcam). L-VGCCs were blocked with nitrendipine (20μM; Abcam). L-VGCC Ca^2+^ transients were isolated by using mibefradil (10μM; Tocris), SNX-482 (0.3μM; Abcam), and w-conotoxin-MVIIC (1μM; Abcam) (cite). NO synthase was inhibited by incubation with L-NAME (100μM; Sigma), which started 20 minutes prior to experimentation. Extracellular NO was scavenged by bath application of cPTIO (50-100μM; Sigma). Intracellular NO was scavenged by bolus-loading cells with 5mM cPTIO. Endocannabinoid signalling (CB1 receptor) was inhibited by bath application of the AM-251 (2μM; Tocris).

### Presynaptic plasticity model

The model ΔP_r_=η(P_depol_-P_glu_) was used to predict changes in P_r_ in our data set of imaged synapses, across all experimental conditions. Data was divided into 6 categories, depending on whether the effects of postsynaptic depolarization were present or absent, whether the effects of glutamate photolysis was present or absent, and whether the effects of presynaptic NMDAR signalling was present or absent. These 6 categories, along with the models predictions for ΔP_r_ are shown in Figure 9C and are summarized in the table below.

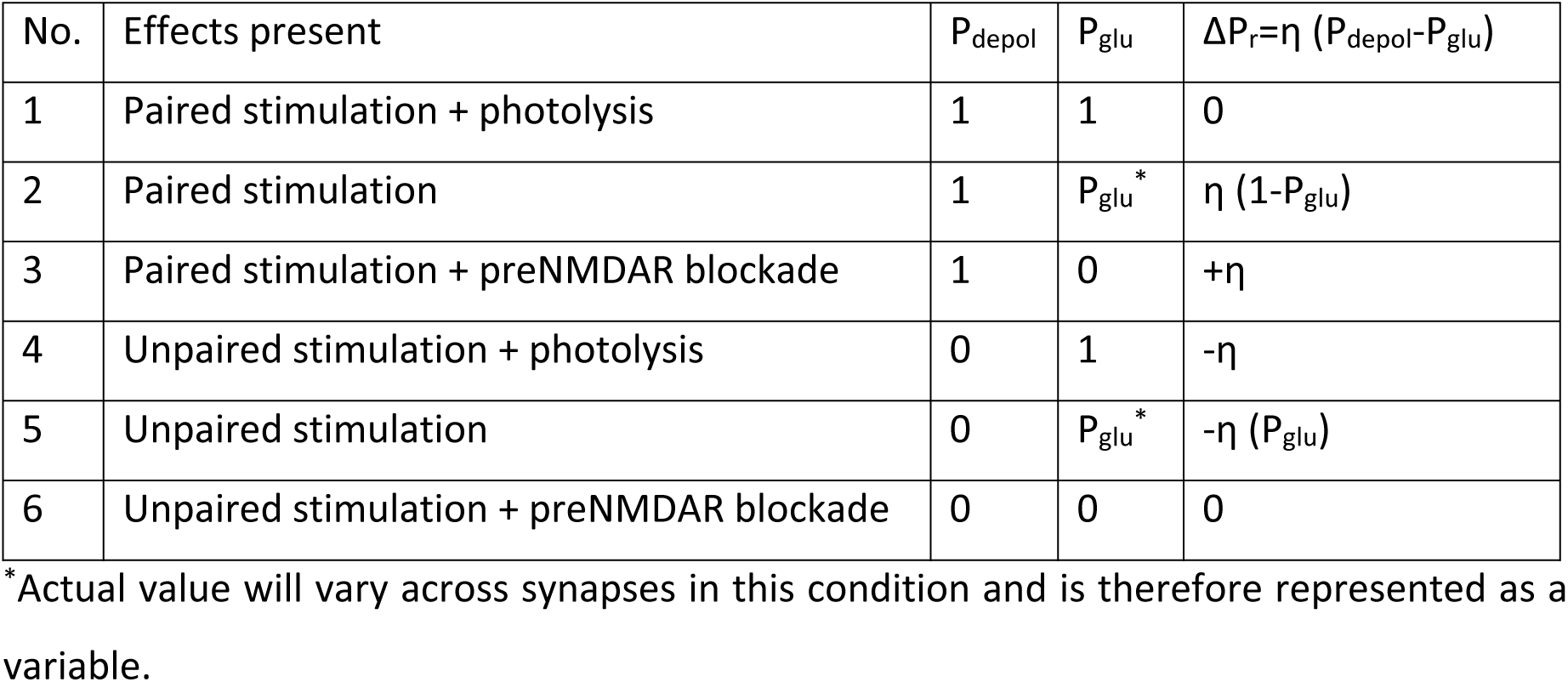

The experimental data in each category are as follows.

Category 1 – included data from experiments in which the effects of postsynaptic depolarization and glutamate photolysis were present during presynaptic stimulation (Figure 2D, 7A). This included experiments in which MK-801 was applied intracellularly during photolysis (Figure 7A). For this category, P_depol_ was set to 1 and P_glu_ was set to 1.

Category 2 – included data from experiments in which the effects of postsynaptic depolarization and endogenous glutamate release were present during presynaptic stimulation (Figure 2D, 4B, 6B). This included experiments in which MK-801 was applied intracellularly (Figure 6B). For this category, P_depol_ was set to 1 and P_glu_ was calculated for each synapse based on: P_glu_=0.475*initial P_r_ + 0.2175, which was experimentally derived from data in Supplemental Figure 12.

Category 3 – included data from experiments in which the effects of postsynaptic depolarization were present but the effects of glutamate release were absent during presynaptic stimulation (Figure 6B,7A). This included experiments in which paired stimulation was delivered in complete glutamate receptor or during bath application of AP5 or MK-801 (Figure 6B). These also included pairing experiments involving glutamate photolysis, but in the presence of MK-801 in the bath (Figure 7A). For this category, P_depol_ was set to 1 and P_glu_ was set to 0.

Category 4 – included data from experiments in which the effects of postsynaptic depolarization were absent but the effects of glutamate photolysis was present during presynaptic stimulation (Figure 2E, 7C). This included experiments in which MK-801 was applied intracellularly during photolysis (Figure 7C). For this category, P_depol_ was set to 0 and P_glu_ was set to 1.

Category 5 – included data from experiments in which the effects of postsynaptic depolarization were absent but the effects of endogenous glutamate release were present during presynaptic stimulation (Figure 2E, 6D). This included experiments in which MK-801 was applied intracellularly during stimulation (Figure 6D). For this category, P_depol_ was set to 0 and P_glu_ was calculated for each synapse based on: P_glu_=0.475*initial P_r_ + 0.2175, which was experimentally derived from data in Supplemental Figure 12.

Category 6 – included data from experiments in which the effects of postsynaptic depolarization and glutamate release were both absent during presynaptic stimulation (Figure 4B, 5A, 6D, 7C). This included: 1) Experiments in which presynaptic stimulation was delivered in complete glutamate receptor blockade, and unaccompanied by postsynaptic depolarization (Figure 4B). 2) Experiments conducted in complete glutamate receptor blockade in which presynaptic stimulation was not delivered during postsynaptic depolarization (Figure 4B). 3) Experiments that took place in complete glutamate receptor blockade and in which the effects of postsynaptic depolarization were abolished, either by nitrendipine, extracellular cPTIO, or intracellular cPTIO (Figure 5A). 4) Experiments conducted with AP5 or MK-801 present in the bath, and in which postsynaptic depolarization was absent (Figure 6D). 5) Experiments done with bath application of MK-801, and in which postsynaptic depolarization was absent but glutamate photolysis was present. (Figure 7C). For this category, P_depol_ was set to 0 and P_glu_ was set to 0.

The learning rate η was determined to be 0.35, this gave the best fit of the model on a small training set of 10 synapses, 5 of which that underwent paired stimulation and 5 of which that underwent unpaired stimulation.

For each category, the initial P_r_ was plotted against final P_r_ for all synapses within the category. The model’s predictions were compared to that of a line of best fit, derived from simple linear regression. The model’s predictions for final P_r_ were set to 1 if they were >1, and were set to 0 if they were <0. For model selection, Bayesian Information Criterion (BIC) was calculated for the model and the lines of best fit as BIC=n*ln(RSS/n) + k*ln(n). Here, n is the number of data points (n=216 synapses), ln is the natural logarithm, RSS is the residual sum of squares and calculated by Σ(model predictions − actual value)^2^, and k is the number of free parameters, which was 1 for the proposed model, and 12 for the lines of best fit (2 free parameters per line of best fit, the slope and the intercept). The BIC aids model selection by favouring models that have a high predictive power (explained variance) but a small number of free parameters. A difference of BIC > 10 between two models strongly favors the model with the smaller BIC value^63^.

### Statistical analysis

Tests used to assess statistical significance are stated at the end of all Figure captions. Only non-parametric tests were used owing to small sample sizes ^107^. For single comparisons, two-tailed Mann-Whitney or Wilcoxon signed rank tests were used, depending on whether the data was unpaired or paired, respectively. Wilcoxon signed rank tests were also used to determine if data significantly differed from an expected value. For multiple comparisons, Kruskal-Wallis tests were used with post-hoc Dunn’s tests. Pearson correlation coefficients were calculated to determine the significance of linear trends. Bayesian Information Criterion (BIC) was used for model selection. Here, a difference of BIC > 10 between two models strongly favors the model with the smaller BIC value^63^ Averages and standard error of the mean (S.E.M.) are represented in the text as average±S.E.M.

## Acknowledgements

Z.P was funded by a Clarendon Scholarship, a scholarship from the National Science and Engineering Research Council of Canada, and a Junior Research Fellowship from Magdalen College, University of Oxford. Experimental work was funded by the Medical Research Council. The funders had no role in study design, data collection and analysis, decision to publish, or preparation of the manuscript

## Author Contributions

Z.P. conceived the idea for study. Z.P. and N.J.E. designed the experiments. Z.P. and R.T. conducted the experiments and analyzed the data. Z.P. and N.J.E prepared the manuscript. All authors have seen and approved the manuscript.

---

**Supplemental Figure 1.**
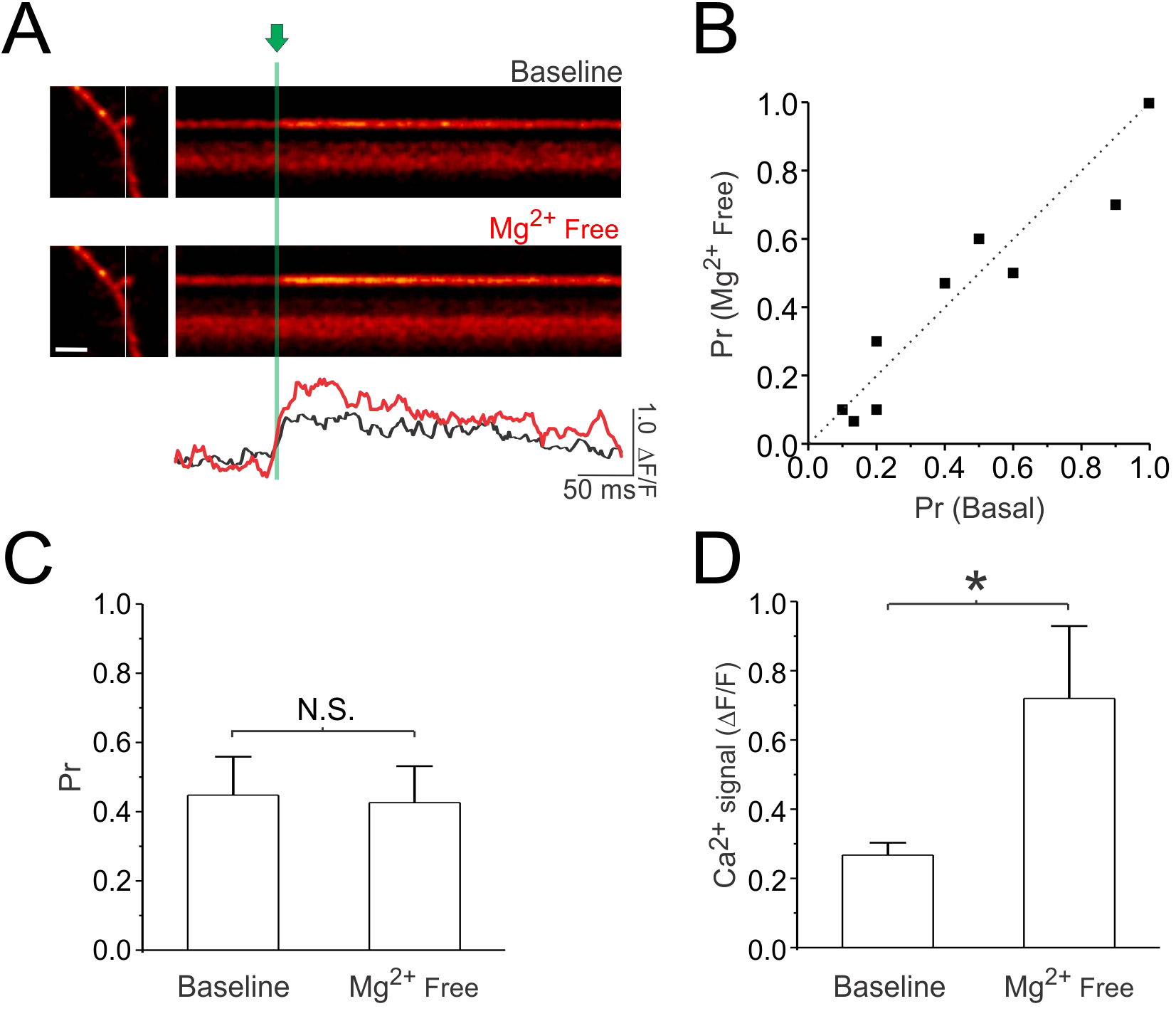
Large changes in Ca^2+^ signal do not alter P_r_ estimates. Ca^2+^ imaging was used to estimate P_r_ at single synapses under baseline conditions (2mM Mg^2+^) and following baseline removal of Mg^2+^ in order to maximize the Ca^2+^ signal in the spine. **(A)** Sample line scan of a spine and underlying dendrite stimulated with a presynaptic pulse (green vertical line) under baseline conditions and after removal of Mg^2+^ from the bath (scale bar: 2μm). The change in fluorescence intensity of the Ca^2+^ response (ΔF/F) is quantified below (black trace = baseline; red trace = Mg^2+^ free). Removal of Mg^2+^ increases the magnitude of the Ca^2+^ response. **(B)** For each synapse imaged, the P_r_ measured under basal conditions is plotted against the P_r_ measured following removal of Mg^2+^. The broken diagonal line represents the expected trend if P_r_ is unchanged. **(C)** Average P_r_ estimate. **(D)** Average change in Ca^2+^ signal (ΔF/F). Removal of Mg^2+^ more than doubles the Ca^2+^ signal on average, but does not change P_r_ estimates. Error bars represent S.E.M. (n=9 spines per condition). Asterisks denote significant differences (*p<0.05; Wilcoxon signed rank test). N.S. denotes no significant difference.

**Supplemental Figure 2.**
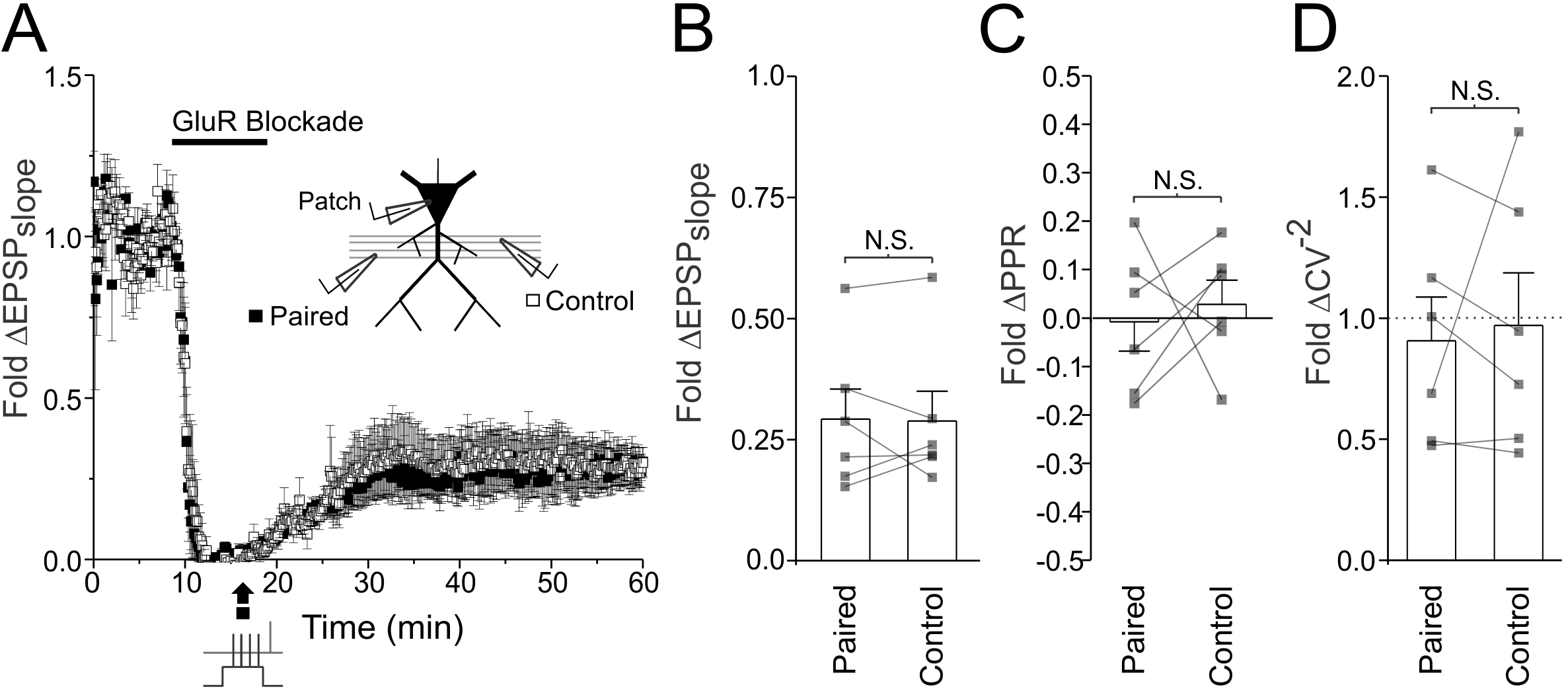
Anti-causal pairing of pre- and postsynaptic activity fails to induce presynaptic LTP in full glutamate receptor blockade. EPSPs were recorded from two independent Schaffer-collateral pathways in hippocampal slices. LTP was induced in full glutamate receptor blockade blockade (100μM D-AP5, 10μM NBQX, 500μM R, S-MCPG, and 10μM LY341495) by 60 anti-causal pairings at 5Hz of single presynaptic stimuli with postsynaptic depolarization in the form of a complex spike (see Figure 1A for example). Only the paired pathway was active during paired stimulation. **(A)** Average fold change in the EPSP slope is plotted against time for both control and paired pathways (n=6). **(B)** Average fold change in EPSP slope in the paired and control pathways 30 minutes following paired stimulation and drug washout. **(C)** Average fold change in CV^-2^ in the paired and control pathways. **(D)** Average change in PPR for paired and control pathways. When compared to the control pathway, anti-causal pairings failed to elicit changes in the EPSP slope. No changes in CV^-2^ or PPR were observed. Error bars represent S.E.M. (n=6 cells per condition). N.S. denotes no significant differences (Wilcoxon signed rank test).

**Supplemental Figure 3.**
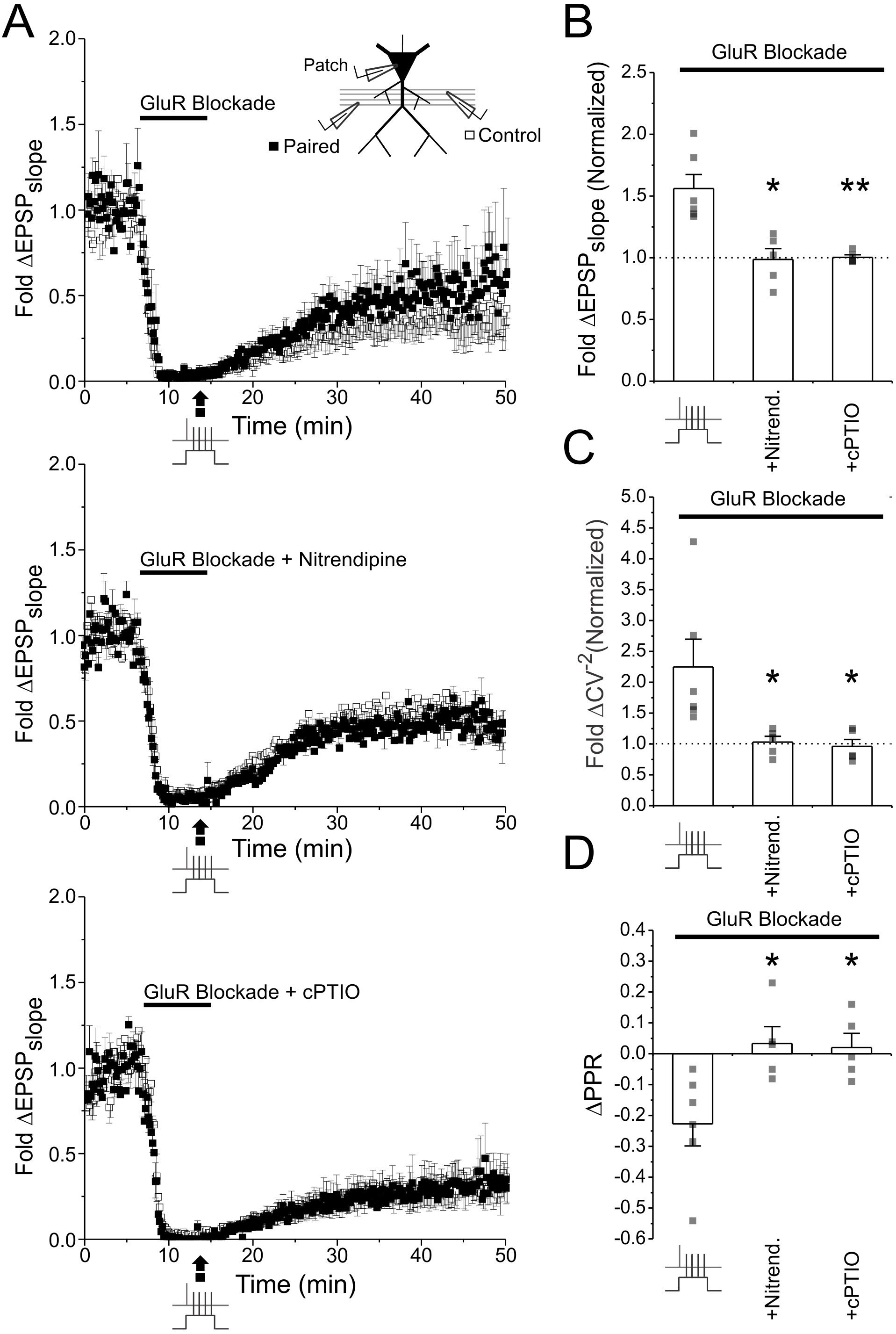
Induction of presynaptic LTP under full glutamate receptor blockade in acute hippocampal slices. EPSPs were recorded from two independent Schaffer-collateral pathways in acute hippocampal slices. LTP was induced by pairing each of 60 presynaptic stimuli, delivered at 5Hz in glutamate receptor blockade (100μM D-AP5, 10μM NBQX, 500μM R,S-MCPG, and 10μM LY341495), with postsynaptic depolarization in the form of a complex spike (see Figure 1A for example). Only the paired pathway was active during paired stimulation. **(A)** Average fold change in the EPSP slope is plotted against time for both control and paired pathways (n=6). Paired stimulation was delivered in glutamate receptor blockade alone, or in the additional presence of the L-VGCC blocker nitrendipine (20μM), or the NO scavenger cPTIO (100μM). **(B)** Average fold change in EPSP slope in the paired pathway 30 minutes following paired stimulation and drug washout; values are normalized to the EPSP slope recorded in the control pathway. **(C)** Average fold change in CV^-2^ in the paired pathway; values are normalized to the EPSP slope recorded in the control pathway. **(D)** Average change in PPR in the paired pathway. Paired stimulation elicited potentiation of the EPSP slope, and was accompanied by an increase in CV^-2^ and a decrease in PPR, suggesting that LTP induction had a presynaptic component of expression. No such changes were observed when pairing occurred in the presence of nitrendipine or cPTIO. Error bars represent S.E.M. (n=5-6 cells per condition). Asterisks denote significant differences from the first group in the graphs (*p<0.05; **p<0.01; Kruskal-Wallis with post-hoc Dunn’s test).

**Supplemental Figure 4.**
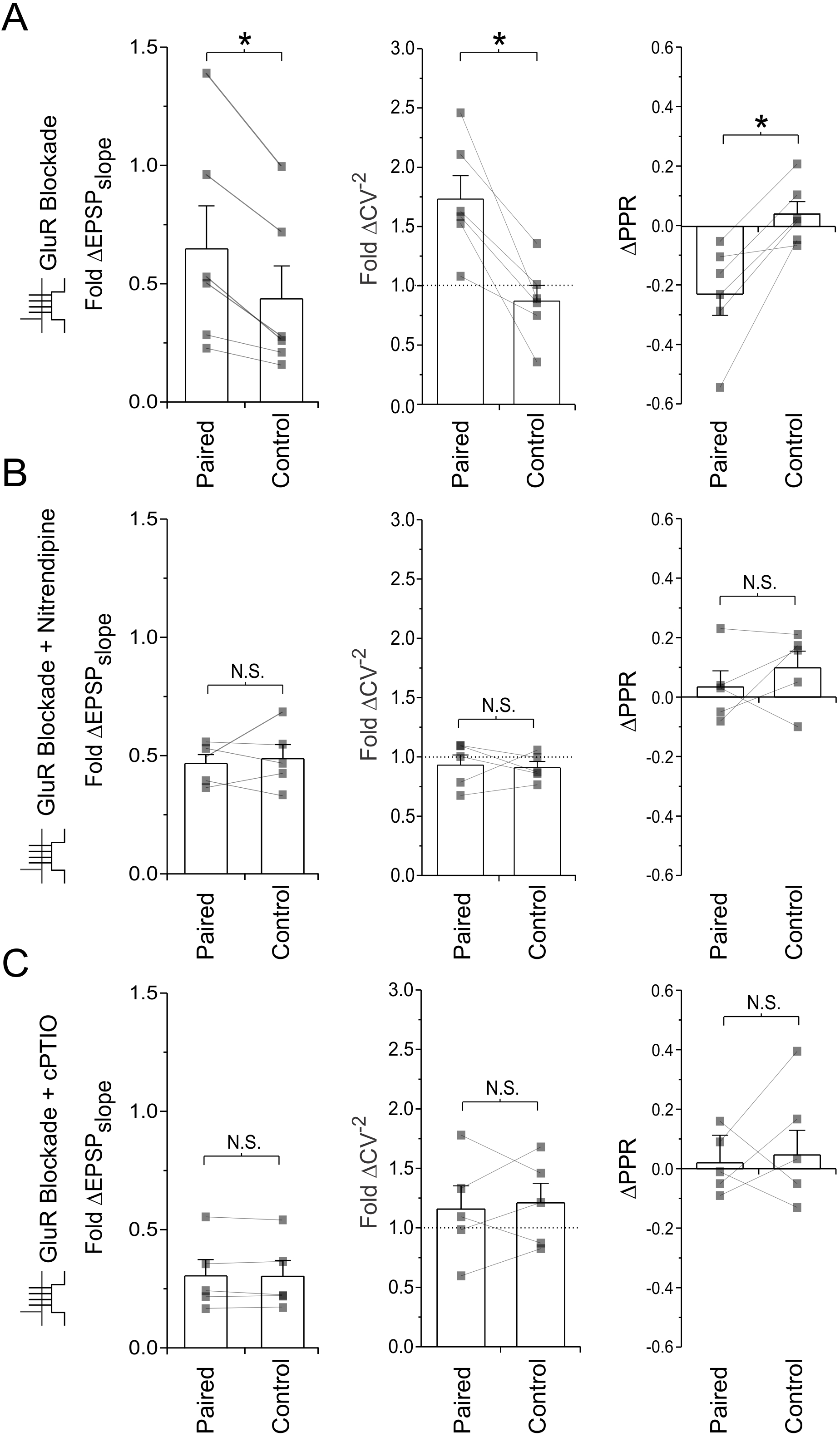
Data summary for acute slice experiments. Same experiment as Figure S1, but data are individually plotted for both the paired and control pathways, without normalization. Fold change in the EPSP slope, fold change in CV^-2^, and change in PPR are plotted for experiments involving **(A)** paired stimulation in the presence of glutamate receptor antagonists, **(B)** paired stimulation in the presence of glutamate receptor antagonists and the L-VGCC antagonist nitrendipine, or **(C)** paired stimulation in the presence of glutamate receptor antagonists and the NO scavenger cPTIO. The presence of nitrendipine and cPTIO prevented pairing-induced increases in EPSP slope, increases in CV^-2^, and decreases in PPR. Error bars represent S.E.M. (n=5-6 cells per condition). Asterisks denotes significance differences between groups (*p<0.05; Wilcoxon signed rank test). N.S. denotes no significant differences between groups.

**Supplemental Figure 5.**
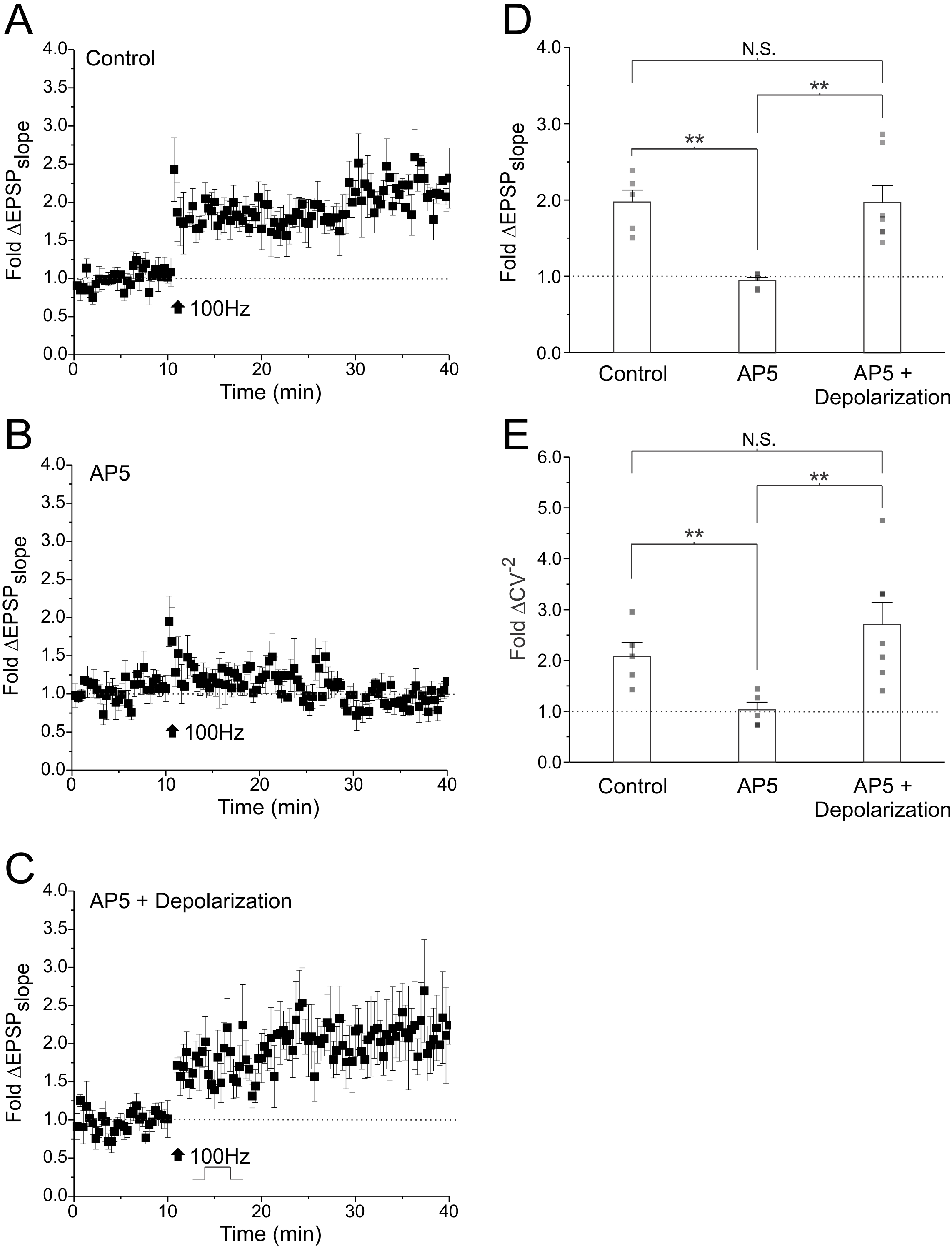
Membrane depolarization rescues LTP induction in NMDAR blockade. EPSPs were recorded from CA1 neurons in hippocampal slice cultures. After baseline recording, tetanic stimulation (black arrow; 3 trains of 20 pulses at 100 Hz delivered 10s apart) was delivered under **(A)** control conditions, **(B)** in the presence of AP5 (100μM D-AP5), and **(C)** in the presence of AP5 and membrane depolarization but in conjunction with of a 10mV membrane depolarization (from -60mV to -50mV) induced by somatic current injection. Membrane depolarization during tetanic stimulation was able to rescue the induction of LTP in AP5. **(D)** Average fold change in the EPSP slope. **(E)** Average fold change in CV^-2^. Values over 1 indicate that LTP induction was associated with a presynaptic component of expression. Error bars represent S.E.M. (n=5-7 cells per condition). Asterisks denotes significance differences between groups (**p<0.01; Kruskal-Wallis with post-hoc Dunn’s test). N.S. denotes no significant differences between groups.

**Supplemental Figure 6.**
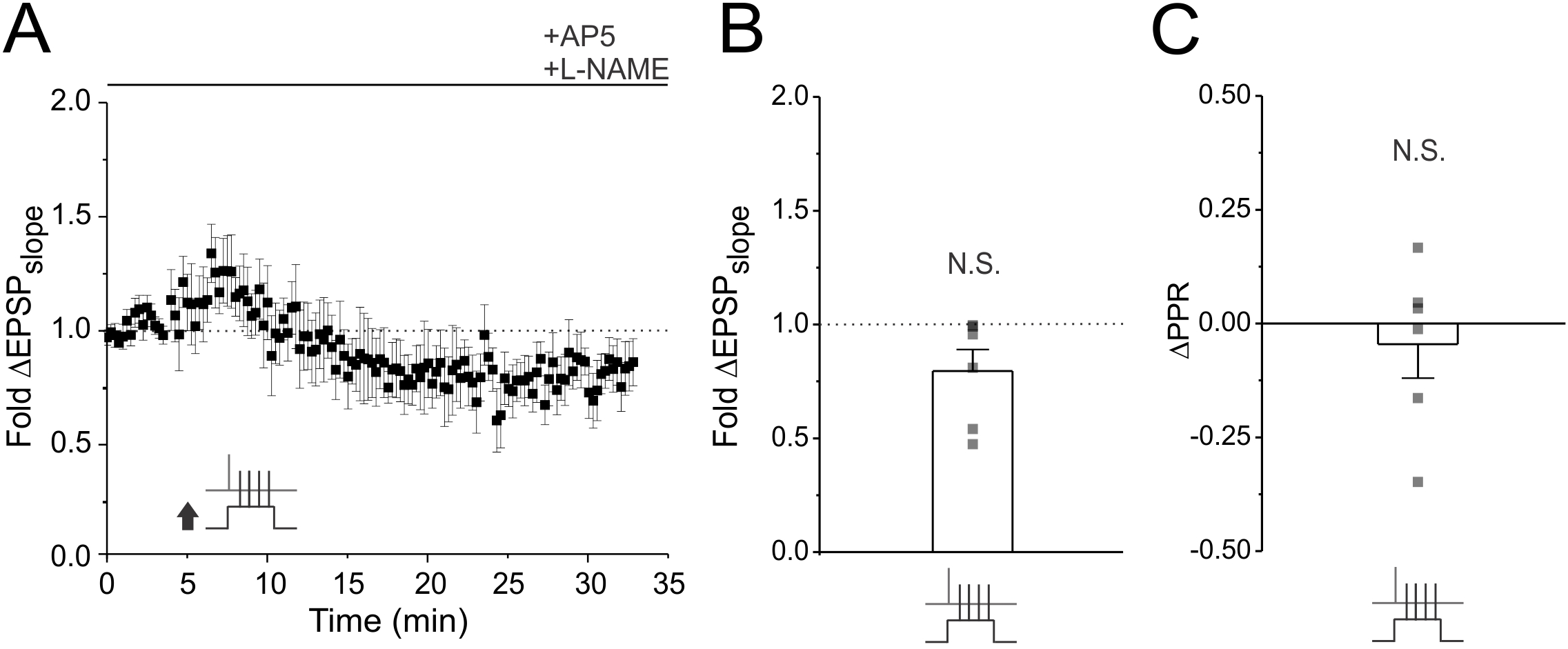
The NO synthase inhibitor L-NAME blocks presynaptic potentiation of EPSPs. **(A)** Average fold change in EPSP slope is plotted against time at baseline and following paired stimulation (60 pairings at 5Hz). Slices were treated with L-NAME to block NO signalling and AP5 to isolate the presynaptic component of LTP. **(B)** Average EPSP slope change across experiments. **(C)** Average PPR change across experiments. L-NAME prevent presynaptic enhancements induced by paired stimulation (Main Figure 8). Error bars represent S.E.M. (n=6 cells per condition). N.S. denotes no significant differences (Wilcoxon signed rank test).

**Supplemental Figure 7.**
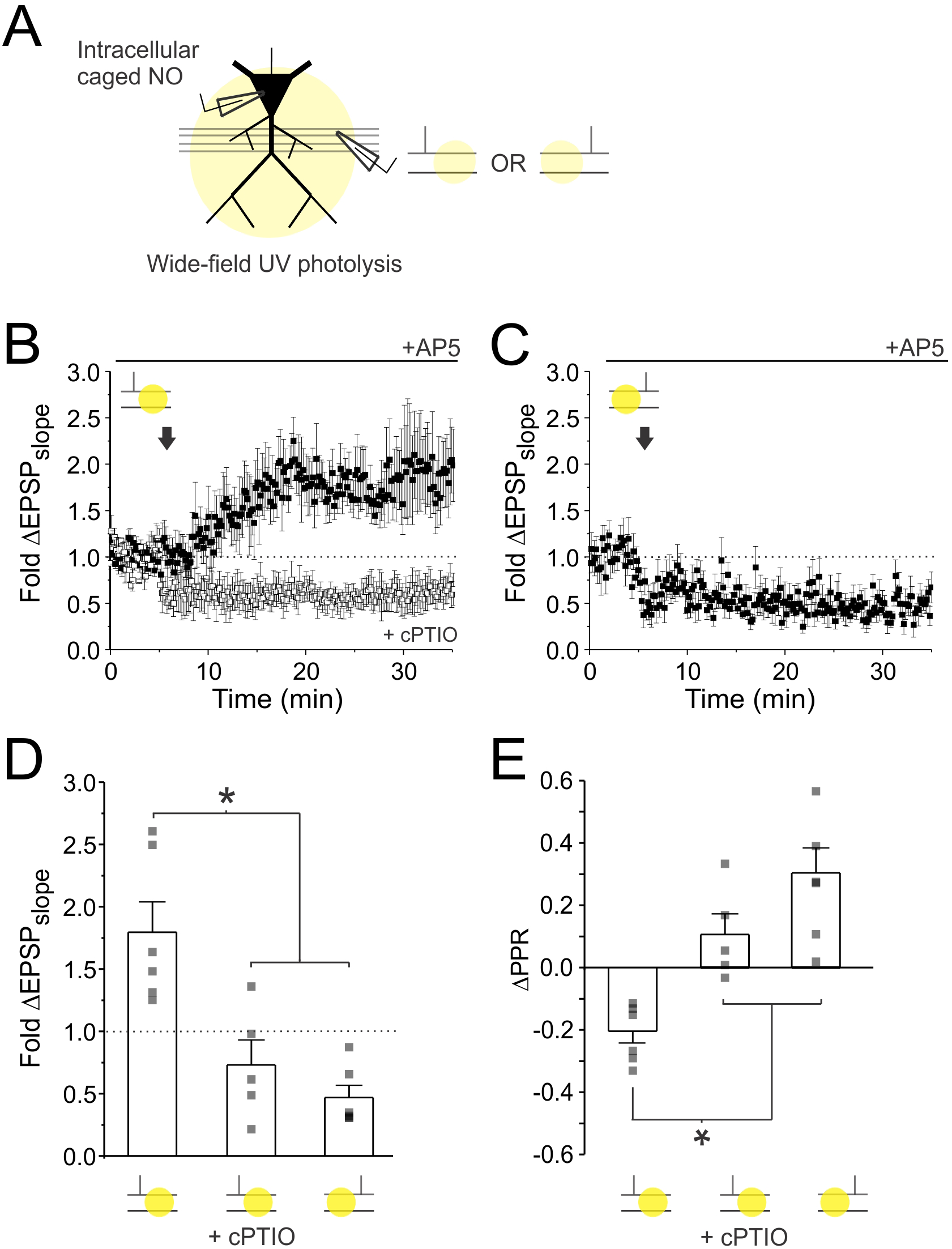
Causal pairing of NO photolysis with presynaptic stimulation induces presynaptic LTP of EPSPs recorded in acute slices. **(A)** Experimental setup. CA1 pyramidal neurons in acute slices were patched and intracellularly loaded with caged NO. EPSPs were recorded in AP5 (50μM D-AP5). A UV flash lamp was used to trigger rapid wide-field uncaging (large yellow circle) either 7-10ms before or after presynaptic stimulation (60 pulses at 5 Hz). **(B)** Average fold change in EPSP slope is plotted against time at baseline and following casual pairing of presynaptic activity with NO photolysis (arrow), either in the presence or absence of cPTIO. **(C)** Average fold changes in EPSP slope is plotted against time at baseline and following anti-causal pairing of presynaptic activity with NO photolysis. **(D)** Average fold change in EPSP slope. **(E)** Average change in PPR. Only casual pairings of presynaptic activity and NO photolysis induced presynaptic LTP. Error bars represent S.E.M. (n=5-6 cells per condition). Asterisks denotes significance differences between groups (*p<0.05; Kruskal-Wallis with post-hoc Dunn’s tests).

**Supplemental Figure 8.**
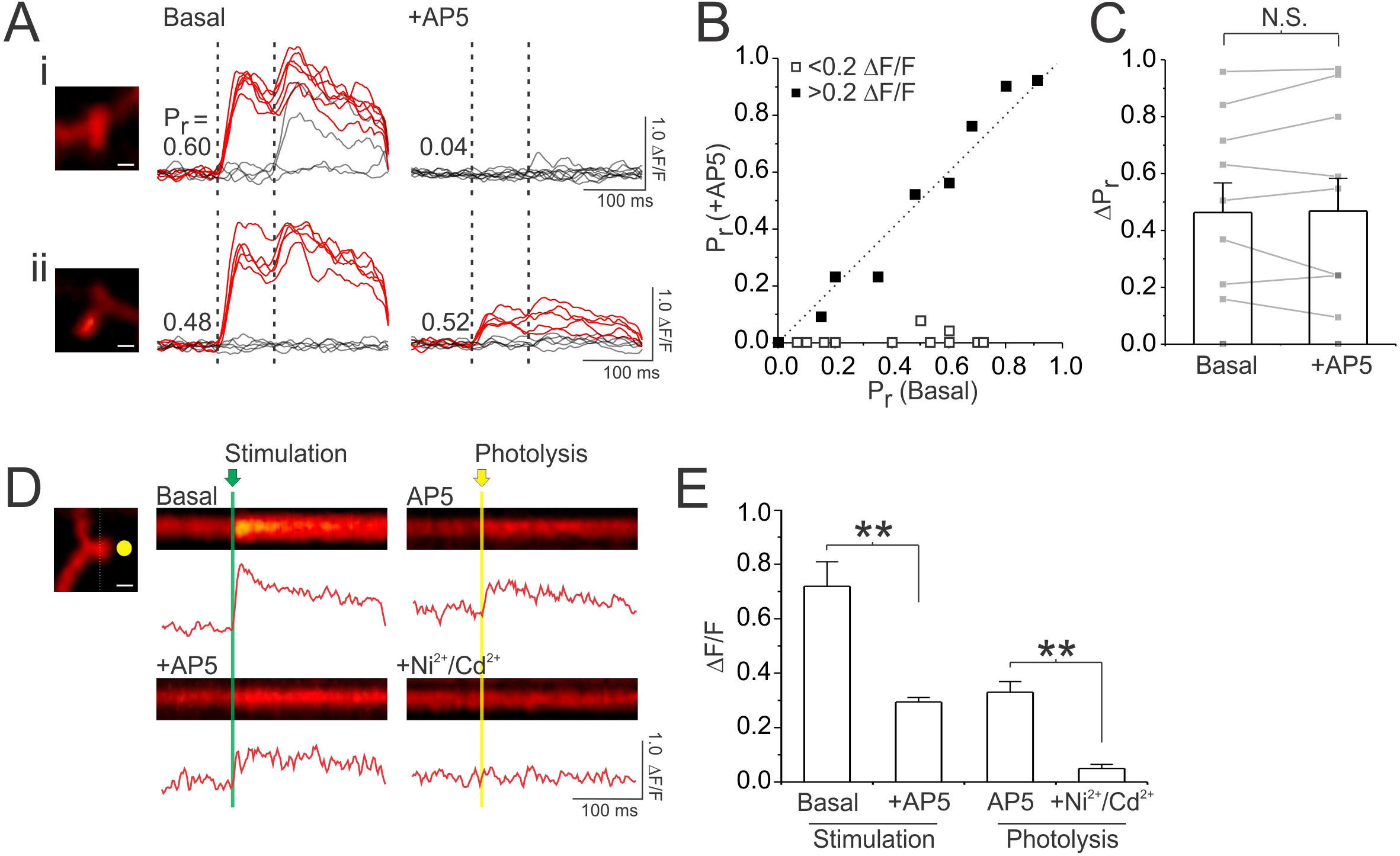
Spine Ca^2+^ transients in NMDA receptor blockade are mediated by voltage-gated Ca^2+^ channels, and can be used to accurately measure P_r_. **(A)** NMDAR blockade does not impede measurements of P_r_. Samples of 10 superimposed Ca^2+^ traces evoked in imaged spines (white scale bar: 1μm) by paired pulse stimulation (broken vertical lines show 2 stimuli delivered 70ms apart); red traces depict successful release events to the first of the two pulses. P_r_ was calculated as the proportion of total stimulation trials in which the first pulse resulted in a Ca^2+^ response in the dendritic spine. **(Ai,ii)** For two imaged spines, sample Ca^2+^ traces are shown prior to and following bath application of AP5 (50μM D-AP5). AP5 abolished Ca2^+^ transients (ΔF/F<0.2) in only one of the two spines. **(B)** For each synapse imaged, the P_r_ measured under basal conditions is plotted against the P_r_ measured during bath application of AP5. The broken diagonal line represents the expected trend if P_r_ is unchanged. For about 50% of synapses, a significant Ca^2+^ transient remained in AP5 (black boxes; ΔF/F > 0.2); at these sites, P_r_ measured under basal conditions was unchanged by AP5. **(C)** Group data depicted no significant difference between P_r_ measured under basal conditions and in AP5 for synapses for which an AP5-insensitive Ca^2+^ transient could be detected (ΔF/F > 0.2). **(D)** Residual Ca^2+^ influx in AP5 is mediated by VGCCs. Images of sample Ca^2+^ responses evoked by stimulation or glutamate photolysis at a single spine under various pharmacological manipulations. Below each image are traces quantifying the change in fluorescence intensity of the Ca^2+^ response (ΔF/F). At the spine shown, AP5 reduced, but did not abolish, Ca^2+^ responses evoked by stimulation. Inhibition of VGCCs by Ni^2+^ (100μM) and Cd^2+^(100μM) abolished the AP5-insensitive component of the Ca^2+^ transient; since Ni^2+^ and Cd^2+^ inhibit transmitter release, glutamate release was simulated via photolysis under these conditions. **(E)** Average peak amplitude of evoked Ca^2+^ responses (ΔF/F) across experimental conditions. Ca^2+^ responses evoked by glutamate photolysis did not significantly differ from those evoked by electrical stimulation, and was significantly reduced by Ni^2+^ and Cd^2+^ application. Error bars represent S.E.M. (n=5-12 spines per condition). Asterisks denote significant differences (**p<0.01; Mann-Whitney or Wilcoxon signed rank tests). N.S. denotes no significant difference.

**Supplemental Figure 9.**
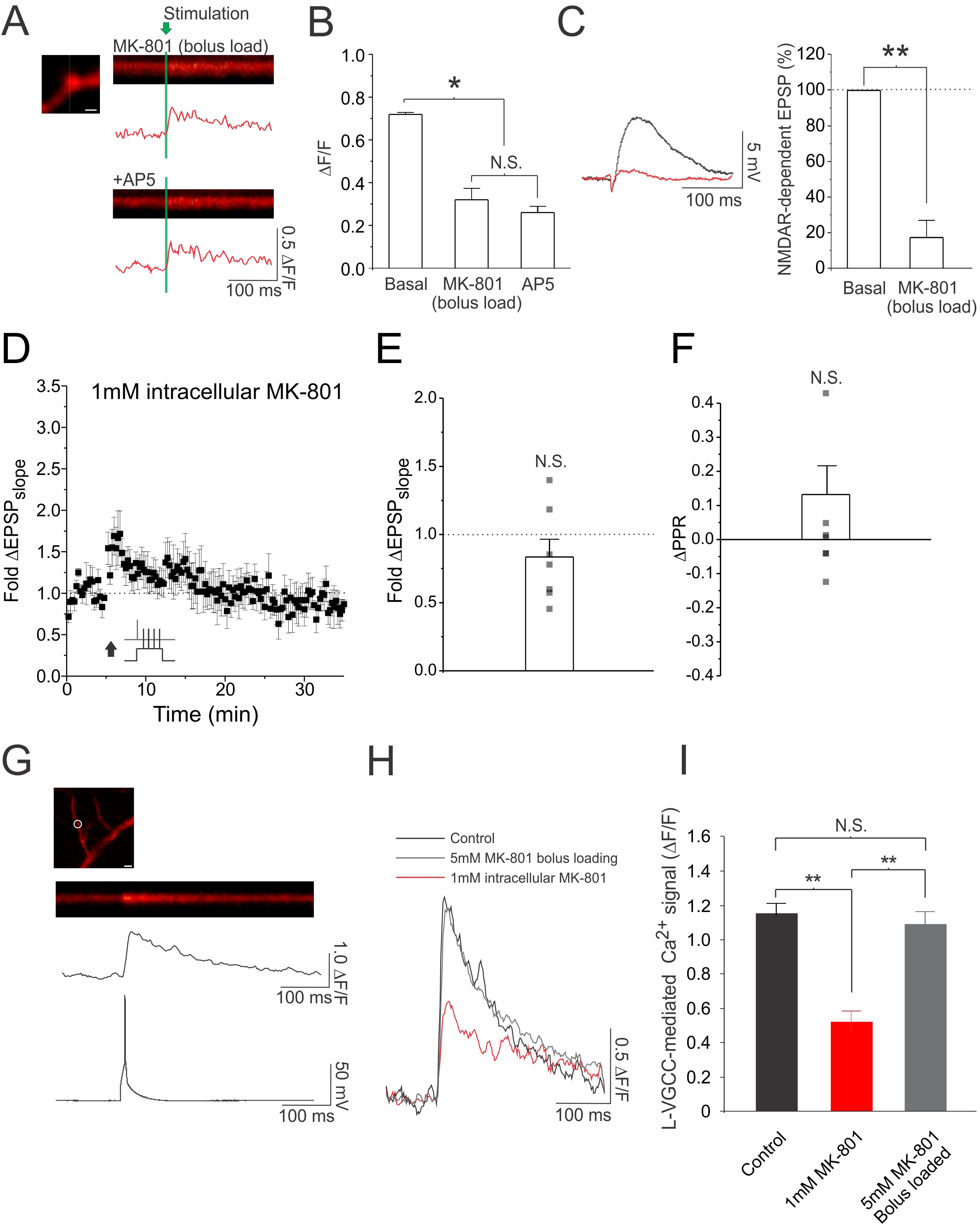
Bolus-loading of MK-801 intracellularly blocks NMDARs without effecting L-VGCCs. MK-801 bolus-loading consisted of transiently patching CA1 neurons (60s) with glass electrodes (4-8MΩ) containing 5mM MK-801. The drug was given approximately 20 minutes to diffuse and take effect before starting experiments. **(A)** Images of sample spine Ca^2+^ responses evoked by single presynaptic stimuli taken 20 minutes after bolus loading of MK-801, and following the bath application of AP5 (scale bar: 1μm). Below each image are traces quantifying the change in fluorescence intensity of the Ca^2+^ response (ΔF/F). Bolus-loading of MK-801 occluded the effects of AP5 on spine Ca^2+^ signals. **(B)** Average peak Ca^2+^ fluorescence amplitude (ΔF/F) evoked by electrical stimulation in control conditions, and following either bolus-loading of MK-801 or bath application of AP5 (50μM D-AP5); response amplitudes recorded in AP5 and MK-801 did not differ, and were significantly lower than those recorded in control conditions, in which NMDARs were not blocked. **(C)** Left. Sample trace of NMDAR-dependent EPSPs recorded in AMPAR blockade (10μm NBQX) prior to and following bolus loading of MK-801 in the recorded neuron. Bolus-loading of MK-801 abolished the NMDAR-dependent EPSP. Right. Average NMDAR-dependent EPSP recorded prior to and following bolus loading of MK-801. **(D)** Average fold change in EPSP slope plotted against time, before and after paired stimulation (black arrow). CA1 pyramidal neurons were recorded with patch-electrodes (4-8 MΩ) containing 1mM MK-801. **(E)** Average change in EPSP slope and **(F)** PPR. No LTP was obtained under these conditions. LTP, however, was obtained when MK-801 was bolus-loaded instead of being continuously present in the recording electrode (Main Figure 9A). **(G)** Sample image of a L-VGCC dependent dendritic Ca^2+^ signal evoked by an action potential (scale bar: 5μm). Ca^2^+ transients were recorded in a cocktail of VGCCs antagonists (10μM mibefradil, 0.3μM SNX-482, and 1μM ɯ-conotoxin-MVIIC) in order to isolate L-VGCC-mediated Ca^2+^ transients [cite]. Below the image is a trace quantifying the change in fluorescence intensity of the Ca^2+^ response (ΔF/F), and a trace showing the simultaneously recorded membrane potential. **(H)** Sample of L-VGCC mediated Ca^2+^ signals recorded from a patched CA1 neuron under control conditions, following bolus-loading of MK-801 (5mM for 60s), and under conditions in which the cell is patched with an electrode containing 1mM MK-801 for 5 minutes prior to imaging. To ensure dilution of postsynaptic factors did not confound experimental results, for the bolus-loading methods, neurons were re-patched for 5 minutes with pipettes containing standard internal solution prior to imaging. Bolus-loading of MK-801 did not effect L-VGCC Ca^2+^ transients triggered in response to single action potentials, whereas having 1mM MK-801 in the patch pipette did. **(I)** Average changes in the L-VGCC mediated Ca^2+^ signal. Error bars represent S.E.M. (n=5-12 spines/cells per condition). Asterisks denote significant differences (**p<0.01; *p<0.05; Wilcoxon signed rank test, Mann-Whitney test, or Kruskal-Wallis with post-hoc Dunn’s test). N.S. denotes no significant difference.

**Supplemental Figure 10.**
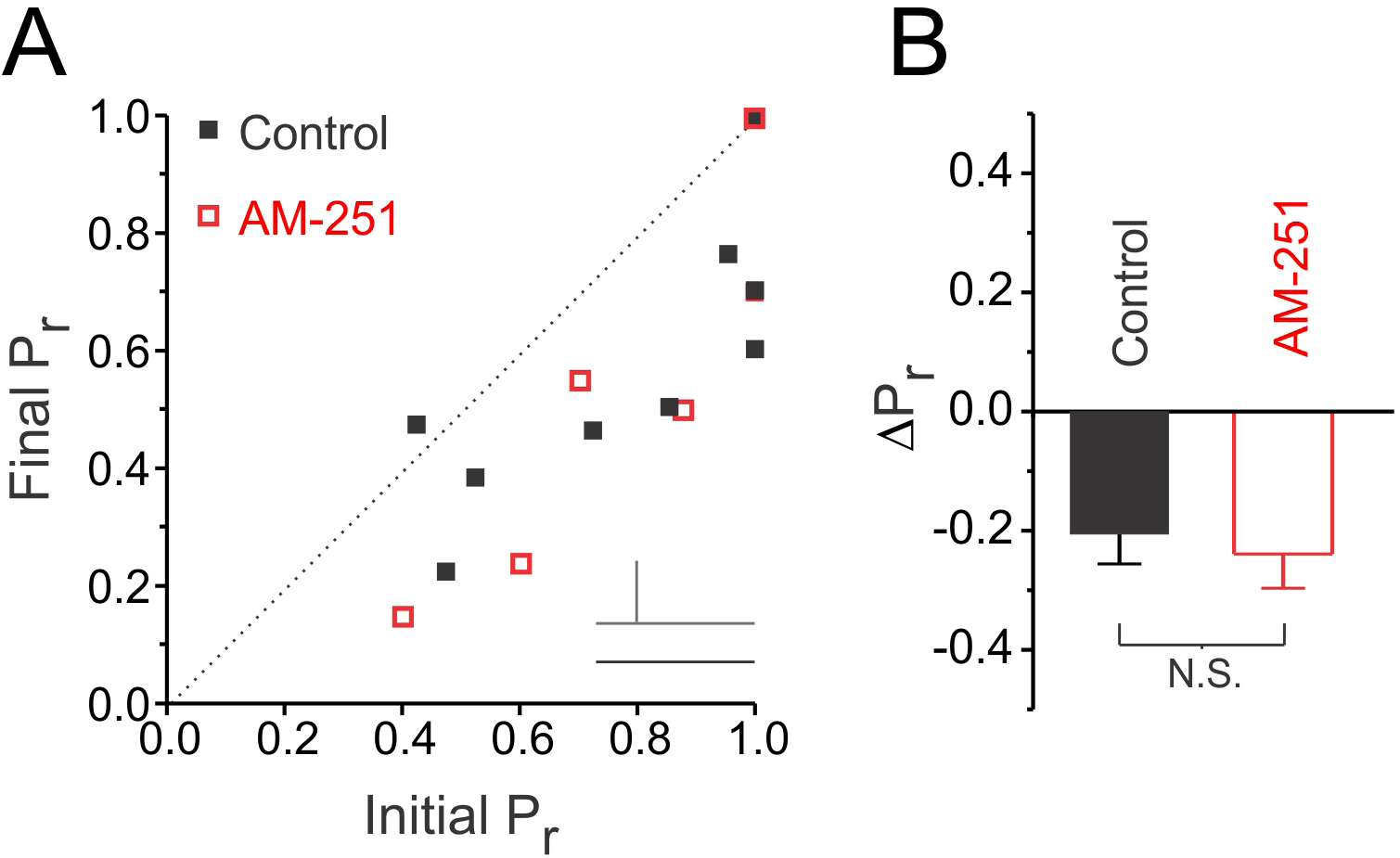
Presynaptic LTD does not require endocannabinoid signalling. **(A)** Ca^2+^ imaging was used to determine P_r_. LTD was induced by delivering 60 presynaptic stimuli at 5Hz at high P_r_ synapses either in the presence or absence of the endocannabinoid antagonist AM-251 (2μM). The final P_r_ measured 25-30 minutes following LTD induction was plotted against the initial P_r_ measured at baseline for each imaged synapse. The broken diagonal line represents the expected trend if P_r_ was unchanged. LTD was induced in both conditions. **(B)** Average changes in P_r_. Error bars represent S.E.M. (n=6-9 spines per condition). N.S. denotes no significant difference (Mann-Whitney test).

**Supplemental Figure 11.**
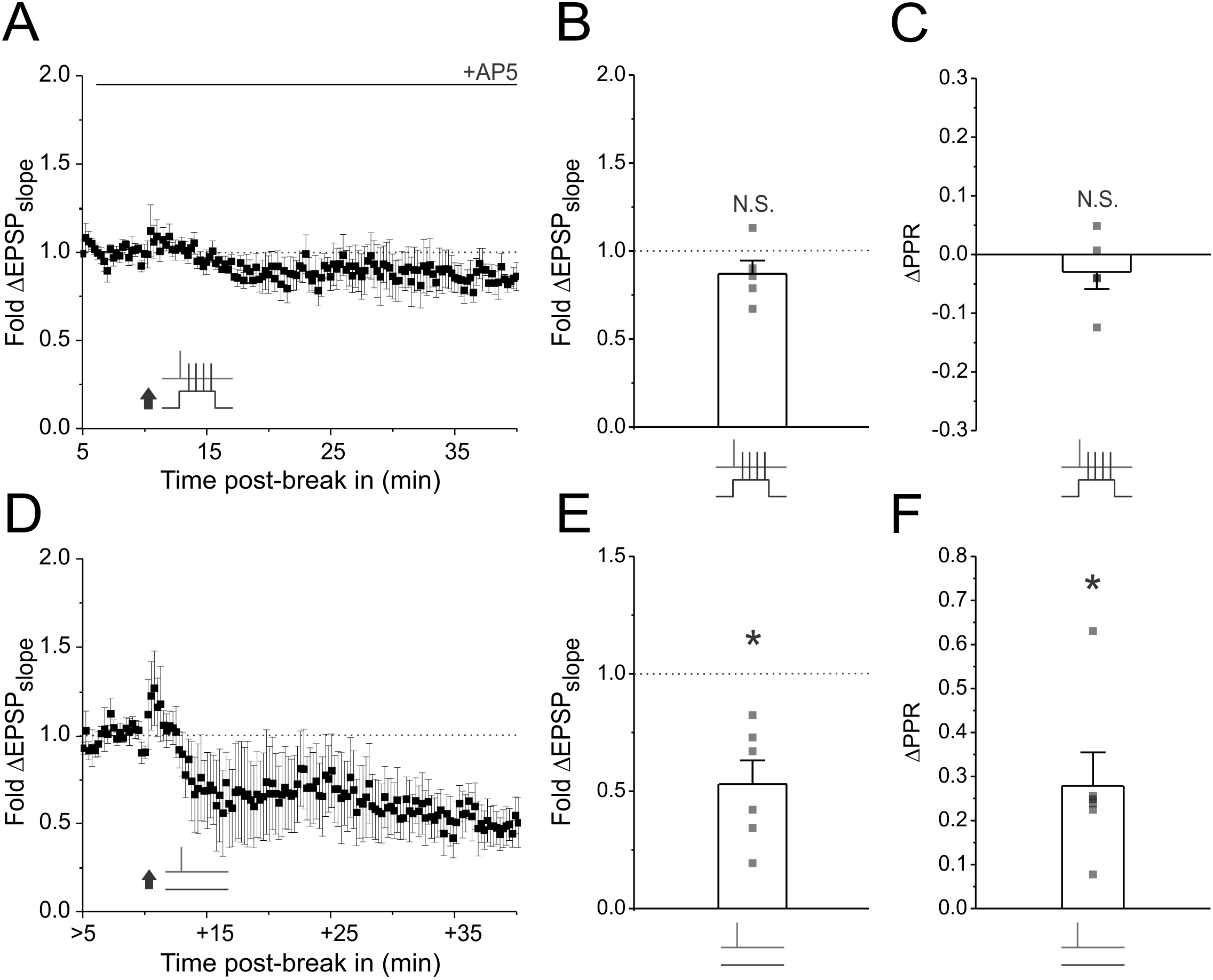
Presynaptic LTD, but not LTP, is resilient to dilution of postsynaptic factors during whole-cell patch recordings. **(A,D)** CA1 neurons were recorded with regular sized patch electrodes (4-8MΩ). Average fold change in EPSP slope is plotted against time at baseline and following plasticity induction under control conditions. Plasticity was induced either using paired stimulation (60 pairings at 5Hz) to induce LTP, or by delivering high frequency bursts of presynaptic stimuli (2 pulses at 200Hz repeated 60 times at 5Hz) in the absence of postsynaptic depolarization to induce LTD. In LTP experiments, plasticity was induced 10 minutes post-break in and in the presence of AP5 (50μM D-AP5) to isolate the presynaptic component of LTP expression. In LTD experiments, plasticity was induced between 10-60 minutes post-break in. **(B,E)** Average EPSP slope change across experiments. **(C,F)** Average PPR change across experiments. LTD but not LTP was resilient to dilution of postsynaptic factors. Error bars represent S.E.M. (n=5-6 cells per condition). Asterisks denote significant differences (*p<0.05;**p<0.01; Wilcoxon signed rank test). N.S. denotes no significant differences.

**Supplemental Figure 12.**
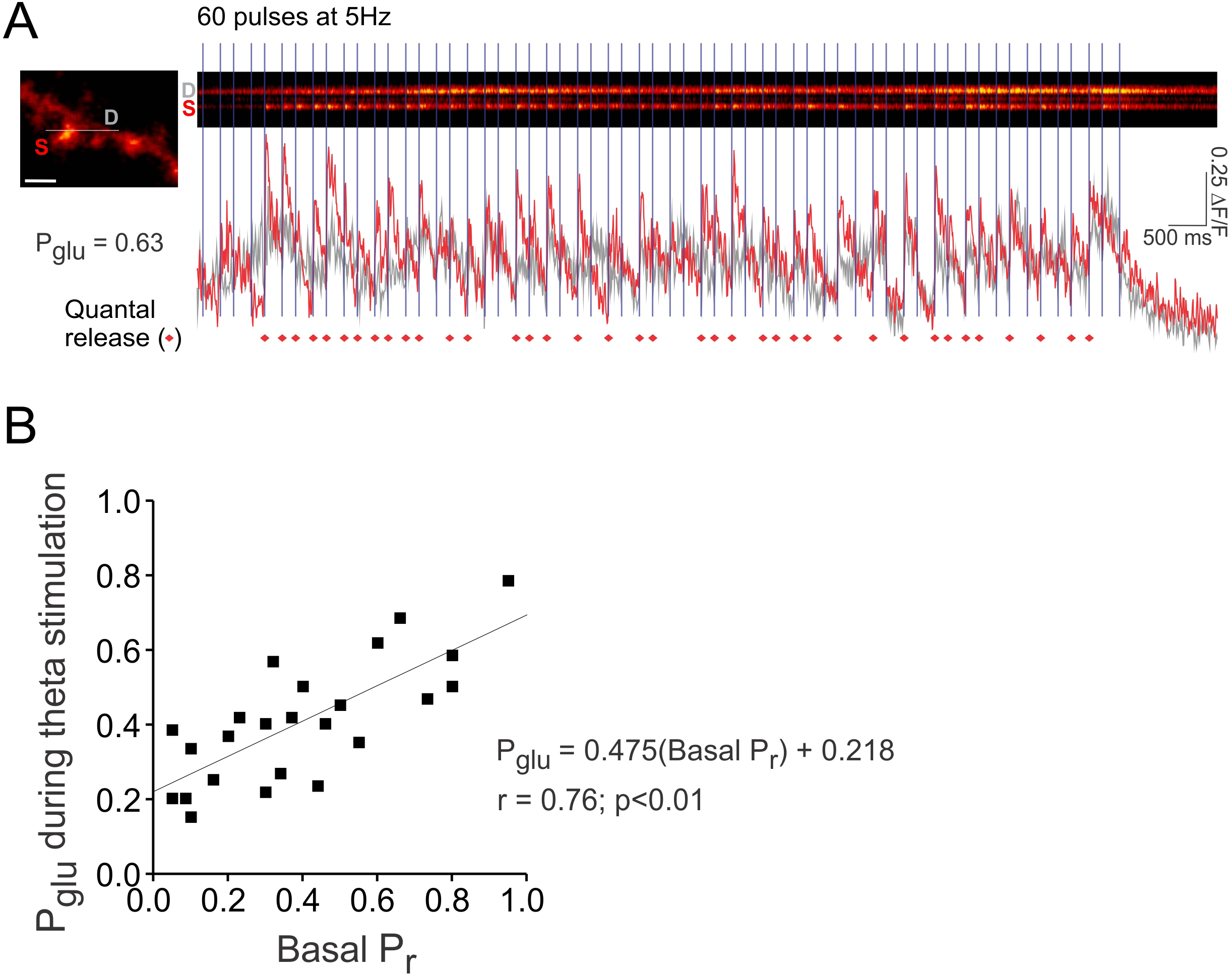
Estimating glutamate release probability during 5Hz theta stimulation. Ca^2+^ imaging was used to monitor glutamate release at single synapses during a presynaptic stimulus train consisting of 60 stimuli delivered at 5Hz. **(A)** Left. Sample image of a spine (S) responsive to stimulation (scale bar = 2μm). Laser scanning was restricted to a line (white line) across the spine and underlying dendrite (D). (Right) The resulting line scan is shown, along with the quantified fluorescence (ΔF/F) of the spine (red) and underlying dendrite (grey). During imaging, 60 presynaptic pulses were delivered at 5Hz (vertical bars). Quantal release of glutamate is marked (red diamonds) whenever the fluorescence of the spine, in response to stimulation, was greater than baseline noise and greater than the fluorescent signal in the dendrite. The comparison to dendritic fluorescence was necessary to rule out any fluorescence changes due to dendritic or somatic spikes evoked by the stimulation. The probability of glutamate release during the train (P_glu_) was calculated as the number of quantal release events divided by the number of presynaptic pulses (i.e. 60). **(B)** For each synapse imaged, P_glu_ was plotted against basal P_r_. The data was fit with a linear regression line to derive a basis by which to estimate P_glu_ given basal P_r_. The regression line was significant (n=24 spines; p<0.01).

## References

1 Padamsey, Z. & Emptage, N. J. Imaging synaptic plasticity. Mol Brain 4, 36, doi:10.1186/1756-6606-4-36 (2011).

2 Bliss, T. V. & Collingridge, G. L. Expression of NMDA receptor-dependent LTP in the hippocampus: bridging the divide. Mol Brain 6, 5, doi:1756-6606-6-5 [pii] 10.1186/1756-6606-6-5 (2013).

3 Larkman, A. U. & Jack, J. J. Synaptic plasticity: hippocampal LTP. Curr Opin Neurobiol 5, 324–334, doi:0959-4388(95)80045-X [pii] (1995).

4 Lisman, J. Long-term potentiation: outstanding questions and attempted synthesis. Philos Trans R Soc Lond B Biol Sci 358, 829–842, doi:10.1098/rstb.2002.1242 (2003).

5 Lisman, J. & Raghavachari, S. A unified model of the presynaptic and postsynaptic changes during LTP at CA1 synapses. Sci STKE 2006, re11, doi:stke.3562006re11 [pii] 10.1126/stke.3562006re11 (2006).

6 Padamsey, Z. & Emptage, N. Two sides to long-term potentiation: a view towards reconciliation. Philos Trans R Soc Lond B Biol Sci 369, 20130154, doi:10.1098/rstb.2013.0154 (2014).

7 Luscher, C. & Malenka, R. C. NMDA receptor-dependent long-term potentiation and longterm depression (LTP/LTD). Cold Spring Harb Perspect Biol 4, doi:cshperspect.a005710 [pii] 10.1101/cshperspect.a005710 (2012).

8 Garthwaite, J. & Boulton, C. L. Nitric oxide signaling in the central nervous system. Annu Rev Physiol 57, 683–706, doi:10.1146/annurev.ph.57.030195.003343 (1995).

9 Yang, Y. & Calakos, N. Presynaptic long-term plasticity. Frontiers in Synaptic Neuroscience 5, doi:10.3389/fnsyn.2013.00008 (2013).

10 Castillo, P. E. Presynaptic LTP and LTD of excitatory and inhibitory synapses. Cold Spring Harbor perspectives in biology 4, a005728 (2012).

11 Ryan, T. A., Ziv, N. E. & Smith, S. J. Potentiation of evoked vesicle turnover at individually resolved synaptic boutons. Neuron 17, 125–134, doi:S0896-6273(00)80286-X [pii] (1996).

12 Ratnayaka, A. et al. Recruitment of resting vesicles into recycling pools supports NMDA receptor-dependent synaptic potentiation in cultured hippocampal neurons. J Physiol 590, 1585–1597, doi:jphysiol.2011.226688 [pii] 10.1113/jphysiol.2011.226688 (2012).

13 Emptage, N. J., Reid, C. A., Fine, A. & Bliss, T. V. Optical quantal analysis reveals a presynaptic component of LTP at hippocampal Schaffer-associational synapses. Neuron 38, 797–804, doi:S0896627303003258 [pii] (2003).

14 Grover, L. M. & Teyler, T. J. N-methyl-D-aspartate receptor-independent long-term potentiation in area CA1 of rat hippocampus: input-specific induction and preclusion in a non-tetanized pathway. Neuroscience 49, 7–11, doi:0306-4522(92)90072-A [pii] (1992).

15 Grover, L. M. & Yan, C. Evidence for involvement of group II/III metabotropic glutamate receptors in NMDA receptor-independent long-term potentiation in area CA1 of rat hippocampus. J Neurophysiol 82, 2956–2969 (1999).

16 Blundon, J. A. & Zakharenko, S. S. Dissecting the components of long-term potentiation. Neuroscientist 14, 598–608, doi:1073858408320643 [pii] 10.1177/1073858408320643 (2008).

17 Zakharenko, S. S. et al. Presynaptic BDNF required for a presynaptic but not postsynaptic component of LTP at hippocampal CA1-CA3 synapses. Neuron 39, 975–990, doi:S0896627303005439 [pii] (2003).

18 Bayazitov, I. T., Richardson, R. J., Fricke, R. G. & Zakharenko, S. S. Slow presynaptic and fast postsynaptic components of compound long-term potentiation. J Neurosci 27, 11510–11521, doi:27/43/11510 [pii] 10.1523/JNEUROSCI.3077-07.2007 (2007).

19 Zakharenko, S. S., Zablow, L. & Siegelbaum, S. A. Visualization of changes in presynaptic function during long-term synaptic plasticity. Nat Neurosci 4, 711–717, doi:10.1038/89498 89498 [pii] (2001).

20 Stricker, C., Cowan, A. I., Field, A. C. & Redman, S. J. Analysis of NMDA-independent longterm potentiation induced at CA3-CA1 synapses in rat hippocampus in vitro. J Physiol 520 Pt 2, 513–525, doi:PHY_9173 [pii] (1999).

21 Grover, L. M. Evidence for postsynaptic induction and expression of NMDA receptor independent LTP. J Neurophysiol 79, 1167–1182 (1998).

22 Pigott, B. M. & Garthwaite, J. Nitric Oxide Is Required for L-Type Ca(2+) Channel-Dependent Long-Term Potentiation in the Hippocampus. Front Synaptic Neurosci 8, 17, doi:10.3389/fnsyn.2016.00017 (2016).

23 Harvey, C. D. & Svoboda, K. Locally dynamic synaptic learning rules in pyramidal neuron dendrites. Nature 450, 1195–1200, doi:nature06416 [pii] 10.1038/nature06416 (2007).

24 Matsuzaki, M., Honkura, N., Ellis-Davies, G. C. & Kasai, H. Structural basis of long-term potentiation in single dendritic spines. Nature 429, 761–766, doi:10.1038/nature02617 nature02617 [pii] (2004).

25 Makino, H. & Malinow, R. AMPA receptor incorporation into synapses during LTP: the role of lateral movement and exocytosis. Neuron 64, 381–390, doi:S0896-6273(09)00675-8 [pii] 10.1016/j.neuron.2009.08.035 (2009).

26 Slutsky, I., Sadeghpour, S., Li, B. & Liu, G. Enhancement of synaptic plasticity through chronically reduced Ca2+ flux during uncorrelated activity. Neuron 44, 835–849, doi:S0896627304007275 [pii] 10.1016/j.neuron.2004.11.013 (2004).

27 Larkman, A., Hannay, T., Stratford, K. & Jack, J. Presynaptic release probability influences the locus of long-term potentiation. Nature 360, 70–73, doi:10.1038/360070a0 (1992).

28 Hardingham, N. R., Hardingham, G. E., Fox, K. D. & Jack, J. J. Presynaptic efficacy directs normalization of synaptic strength in layer 2/3 rat neocortex after paired activity. J Neurophysiol 97, 2965–2975, doi:01352.2006 [pii] 10.1152/jn.01352.2006 (2007).

29 Saez, I. & Friedlander, M. J. Plasticity between neuronal pairs in layer 4 of visual cortex varies with synapse state. J Neurosci 29, 15286–15298, doi:29/48/15286 [pii] 10.1523/JNEUROSCI.2980-09.2009 (2009).

30 McGuinness, L. et al. Presynaptic NMDARs in the Hippocampus Facilitate Transmitter Release at Theta Frequency. Neuron 68, 1109–1127, doi:S0896-6273(10)00939-6 [pii] 10.1016/j.neuron.2010.11.023 (2010).

31 Andrade-Talavera, Y., Duque-Feria, P., Paulsen, O. & Rodriguez-Moreno, A. Presynaptic Spike Timing-Dependent Long-Term Depression in the Mouse Hippocampus. Cereb Cortex 26, 3637–3654, doi:10.1093/cercor/bhw172 (2016).

32 Sjostrom, P. J., Turrigiano, G. G. & Nelson, S. B. Neocortical LTD via coincident activation of presynaptic NMDA and cannabinoid receptors. Neuron 39, 641–654, doi:S0896627303004768 [pii] (2003).

33 Min, R. & Nevian, T. Astrocyte signaling controls spike timing-dependent depression at neocortical synapses. Nat Neurosci 15, 746–753, doi:nn.3075 [pii] 10.1038/nn.3075 (2012).

34 Enoki, R., Hu, Y. L., Hamilton, D. & Fine, A. Expression of long-term plasticity at individual synapses in hippocampus is graded, bidirectional, and mainly presynaptic: optical quantal analysis. Neuron 62, 242–253, doi:S0896-6273(09)00204-9 [pii] 10.1016/j.neuron.2009.02.026 (2009).

35 Thomas, M. J., Watabe, A. M., Moody, T. D., Makhinson, M. & O’Dell, T. J. Postsynaptic complex spike bursting enables the induction of LTP by theta frequency synaptic stimulation. J Neurosci 18, 7118–7126 (1998).

36 Remy, S. & Spruston, N. Dendritic spikes induce single-burst long-term potentiation. Proc Natl Acad Sci U S A 104, 17192–17197, doi:0707919104 [pii] 10.1073/pnas.0707919104 (2007).

37 Golding, N. L., Staff, N. P. & Spruston, N. Dendritic spikes as a mechanism for cooperative long-term potentiation. Nature 418, 326–331, doi:10.1038/nature00854 nature00854 [pii] (2002).

38 Hardie, J. & Spruston, N. Synaptic depolarization is more effective than back-propagating action potentials during induction of associative long-term potentiation in hippocampal pyramidal neurons. J Neurosci 29, 3233–3241, doi:29/10/3233 [pii] 10.1523/JNEUROSCI.6000-08.2009 (2009).

39 Ranck, J. B. Jr., Studies on single neurons in dorsal hippocampal formation and septum in unrestrained rats. I. Behavioral correlates and firing repertoires. Exp Neurol 41, 461–531 (1973).

40 Grienberger, C., Chen, X. & Konnerth, A. NMDA receptor-dependent multidendrite Ca(2+) spikes required for hippocampal burst firing in vivo. Neuron 81, 1274–1281, doi:S0896-6273(14)00019-1 [pii] 10.1016/j.neuron.2014.01.014 (2014).

41 Kowalski, J., Gan, J., Jonas, P. & Pernia-Andrade, A. J. Intrinsic membrane properties determine hippocampal differential firing pattern in vivo in anesthetized rats. Hippocampus 26, 668–682, doi:10.1002/hipo.22550 (2016).

42 Ward, B. et al. State-dependent mechanisms of LTP expression revealed by optical quantal analysis. Neuron 52, 649–661, doi:S0896-6273(06)00806-3 [pii] 10.1016/j.neuron.2006.10.007 (2006).

43 Grunditz, A., Holbro, N., Tian, L., Zuo, Y. & Oertner, T. G. Spine neck plasticity controls postsynaptic calcium signals through electrical compartmentalization. J Neurosci 28, 13457–13466, doi:28/50/13457 [pii] 10.1523/JNEUROSCI.2702-08.2008 (2008).

44 Emptage, N., Bliss, T. V. & Fine, A. Single synaptic events evoke NMDA receptor-mediated release of calcium from internal stores in hippocampal dendritic spines. Neuron 22, 115–124, doi:S0896-6273(00)80683-2 [pii] (1999).

45 Schiller, J. & Schiller, Y. NMDA receptor-mediated dendritic spikes and coincident signal amplification. Curr Opin Neurobiol 11, 343–348, doi:S0959-4388(00)00217-8 [pii] (2001).

46 Grover, L. M. & Yan, C. Blockade of GABAA receptors facilitates induction of NMDA receptor-independent long-term potentiation. J Neurophysiol 81, 2814–2822 (1999).

47 Chemin, J. et al. Mechanisms underlying excitatory effects of group I metabotropic glutamate receptors via inhibition of 2P domain K+ channels. EMBO J 22, 5403–5411, doi:10.1093/emboj/cdg528 (2003).

48 Losonczy, A. & Magee, J. C. Integrative properties of radial oblique dendrites in hippocampal CA1 pyramidal neurons. Neuron 50, 291–307, doi:S0896-6273(06)00213-3 [pii] 10.1016/j.neuron.2006.03.016 (2006).

49 Sattler, R. et al. Specific coupling of NMDA receptor activation to nitric oxide neurotoxicity by PSD-95 protein. Science 284, 1845–1848 (1999).

50 Stanika, R. I., Villanueva, I., Kazanina, G., Andrews, S. B. & Pivovarova, N. B. Comparative impact of voltage-gated calcium channels and NMDA receptors on mitochondria-mediated neuronal injury. J Neurosci 32, 6642–6650, doi:32/19/6642 [pii] 10.1523/JNEUROSCI.6008-11.2012 (2012).

51 Chen, X., Sheng, C. & Zheng, X. Direct nitric oxide imaging in cultured hippocampal neurons with diaminoanthraquinone and confocal microscopy. Cell biology international 25, 593–598, doi:10.1006/cbir.2001.0734 (2001).

52 von Bohlen und Halbach, O., Albrecht, D., Heinemann, U. & Schuchmann, S. Spatial nitric oxide imaging using 1,2-diaminoanthraquinone to investigate the involvement of nitric oxide in long-term potentiation in rat brain slices. Neuroimage 15, 633–639, doi:10.1006/nimg.2001.1045 S1053811901910456 [pii] (2002).

53 Rodriguez-Moreno, A. et al. Presynaptic self-depression at developing neocortical synapses. Neuron 77, 35–42, doi:S0896-6273(12)00956-7 [pii] 10.1016/j.neuron.2012.10.035 (2013).

54 Rodriguez-Moreno, A. & Paulsen, O. Spike timing-dependent long-term depression requires presynaptic NMDA receptors. Nat Neurosci 11, 744–745, doi:nn.2125 [pii] 10.1038/nn.2125 (2008).

55 Nevian, T. & Sakmann, B. Spine Ca2+ signaling in spike-timing-dependent plasticity. J Neurosci 26, 11001–11013, doi:26/43/11001 [pii] 10.1523/JNEUROSCI.1749-06.2006 (2006).

56 Corlew, R., Brasier, D. J., Feldman, D. E. & Philpot, B. D. Presynaptic NMDA receptors: newly appreciated roles in cortical synaptic function and plasticity. Neuroscientist 14, 609–625, doi:14/6/609 [pii] 10.1177/1073858408322675 (2008).

57 Corlew, R., Wang, Y., Ghermazien, H., Erisir, A. & Philpot, B. D. Developmental switch in the contribution of presynaptic and postsynaptic NMDA receptors to long-term depression. J Neurosci 27, 9835–9845, doi:27/37/9835 [pii] 10.1523/JNEUROSCI.5494-06.2007 (2007).

58 Cormier, R. J. & Kelly, P. T. Glutamate-induced long-term potentiation enhances spontaneous EPSC amplitude but not frequency. J Neurophysiol 75, 1909–1918 (1996).

59 Holbro, N., Grunditz, A., Wiegert, J. S. & Oertner, T. G. AMPA receptors gate spine Ca(2+) transients and spike-timing-dependent potentiation. Proc Natl Acad Sci U S A 107, 15975–15980, doi:1004562107 [pii] 10.1073/pnas.1004562107 (2010).

60 Wong, E. H. et al. The anticonvulsant MK-801 is a potent N-methyl-D-aspartate antagonist. Proc Natl Acad Sci U S A 83, 7104–7108 (1986).

61 Jaffe, D. B., Marks, S. S. & Greenberg, D. A. Antagonist drug selectivity for radioligand binding sites on voltage-gated and N-methyl-D-aspartate receptor-gated Ca2+ channels. Neurosci Lett 105, 227–232 (1989).

62 Kim, J. M. et al. Blockade of voltage-gated K(+) currents in rat mesenteric arterial smooth muscle cells by MK801. J Pharmacol Sci 127, 92–102, doi:10.1016/j.jphs.2014.11.005 (2015).

63 Kass, R. E. & Raftery, A. E. Bayes Factors. Journal of the American Statistical Association 90, 773–795, doi:10.1080/01621459.1995.10476572 (1995).

64 Voronin, L. L. & Cherubini, E. ‘Deaf, mute and whispering’ silent synapses: their role in synaptic plasticity. J Physiol 557, 3–12, doi:10.1113/jphysiol.2003.058966 jphysiol.2003.058966 [pii] (2004).

65 Stevens, C. F. Neurotransmitter release at central synapses. Neuron 40, 381–388, doi:S0896627303006433 [pii] (2003).

66 Bell, M. E. et al. Dynamics of Nascent and Active Zone Ultrastructure as Synapses Enlarge During Ltp in Mature Hippocampus. J Comp Neurol 522, 3861–3884, doi:10.1002/cne.23646 (2014).

67 Murthy, V. N., Sejnowski, T. J. & Stevens, C. F. Heterogeneous release properties of visualized individual hippocampal synapses. Neuron 18, 599–612, doi:S0896-6273(00)80301-3 [pii] (1997).

68 Collingridge, G. L., Kehl, S. J. & McLennan, H. Excitatory amino acids in synaptic transmission in the Schaffer collateral-commissural pathway of the rat hippocampus. J Physiol 334, 33–46 (1983).

69 Bashir, Z. I. et al. Induction of LTP in the hippocampus needs synaptic activation of glutamate metabotropic receptors. Nature 363, 347–350, doi:10.1038/363347a0 (1993).

70 Hofmann, F., Lacinova, L. & Klugbauer, N. Voltage-dependent calcium channels: from structure to function. Rev Physiol Biochem Pharmacol 139, 33–87 (1999).

71 Lacinova, L. & Hofmann, F. Ca2+- and voltage-dependent inactivation of the expressed L-type Ca(v)1.2 calcium channel. Arch Biochem Biophys 437, 42–50, doi:S0003-9861(05)00089-5 [pii] 10.1016/j.abb.2005.02.025 (2005).

72 Fuenzalida, M., Fernandez de Sevilla, D., Couve, A. & Buno, W. Role of AMPA and NMDA receptors and back-propagating action potentials in spike timing-dependent plasticity. J Neurophysiol 103, 47–54, doi:00416.2009 [pii] 10.1152/jn.00416.2009 (2010).

73 Kullmann, D. M., Perkel, D. J., Manabe, T. & Nicoll, R. A. Ca2+ entry via postsynaptic voltage-sensitive Ca2+ channels can transiently potentiate excitatory synaptic transmission in the hippocampus. Neuron 9, 1175–1183, doi:0896-6273(92)90075-O [pii] (1992).

74 Huber, K. M., Mauk, M. D. & Kelly, P. T. Distinct LTP induction mechanisms: contribution of NMDA receptors and voltage-dependent calcium channels. J Neurophysiol 73, 270–279 (1995).

75 Grover, L. M., Kim, E., Cooke, J. D. & Holmes, W. R. LTP in hippocampal area CA1 is induced by burst stimulation over a broad frequency range centered around delta. Learn Mem 16, 69–81, doi:16/1/69 [pii] 10.1101/lm.1179109 (2009).

76 Morgan, S. L. & Teyler, T. J. Electrical stimuli patterned after the theta-rhythm induce multiple forms of LTP. J Neurophysiol 86, 1289–1296 (2001).

77 Wilsch, V. W., Behnisch, T., Jager, T., Reymann, K. G. & Balschun, D. When are class I metabotropic glutamate receptors necessary for long-term potentiation? J Neurosci 18, 6071–6080 (1998).

78 Dan, Y. & Poo, M. M. Spike timing-dependent plasticity of neural circuits. Neuron 44, 23–30, doi:10.1016/j.neuron.2004.09.007 S0896627304005768 [pii] (2004).

79 Watanabe, S., Hoffman, D. A., Migliore, M. & Johnston, D. Dendritic K+ channels contribute to spike-timing dependent long-term potentiation in hippocampal pyramidal neurons. Proc Natl Acad Sci U S A 99, 8366–8371, doi:10.1073/pnas.122210599 122210599 [pii] (2002).

80 Gasparini, S., Losonczy, A., Chen, X., Johnston, D. & Magee, J. C. Associative pairing enhances action potential back-propagation in radial oblique branches of CA1 pyramidal neurons. J Physiol 580, 787–800, doi:jphysiol.2006.121343 [pii] 10.1113/jphysiol.2006.121343 (2007).

81 Hoffman, D. A., Magee, J. C., Colbert, C. M. & Johnston, D. K+ channel regulation of signal propagation in dendrites of hippocampal pyramidal neurons. Nature 387, 869–875, doi:10.1038/43119 (1997).

82 Johnston, D., Hoffman, D. A., Colbert, C. M. & Magee, J. C. Regulation of back-propagating action potentials in hippocampal neurons. Curr Opin Neurobiol 9, 288–292, doi:nb9302 [pii] (1999).

83 Migliore, M., Hoffman, D. A., Magee, J. C. & Johnston, D. Role of an A-type K+ conductance in the back-propagation of action potentials in the dendrites of hippocampal pyramidal neurons. J Comput Neurosci 7, 5–15 (1999).

84 Sjostrom, P. J. & Hausser, M. A cooperative switch determines the sign of synaptic plasticity in distal dendrites of neocortical pyramidal neurons. Neuron 51, 227–238, doi:S0896-6273(06)00471-5 [pii] 10.1016/j.neuron.2006.06.017 (2006).

85 Smith, S. L., Smith, I. T., Branco, T. & Hausser, M. Dendritic spikes enhance stimulus selectivity in cortical neurons in vivo. Nature 503, 115–120, doi:10.1038/nature12600 (2013).

86 Williams, J. H., Errington, M. L., Lynch, M. A. & Bliss, T. V. Arachidonic acid induces a longterm activity-dependent enhancement of synaptic transmission in the hippocampus. Nature 341, 739–742, doi:10.1038/341739a0 (1989).

87 Nikonenko, I., Jourdain, P. & Muller, D. Presynaptic remodeling contributes to activity-dependent synaptogenesis. J Neurosci 23, 8498–8505, doi:23/24/8498 [pii] (2003).

88 Stanton, P. K., Winterer, J., Zhang, X. L. & Muller, W. Imaging LTP of presynaptic release of FM1-43 from the rapidly recycling vesicle pool of Schaffer collateral-CA1 synapses in rat hippocampal slices. Eur J Neurosci 22, 2451–2461, doi:EJN4437 [pii] 10.1111/j.1460-9568.2005.04437.x (2005).

89 Johnstone, V. P. & Raymond, C. R. A protein synthesis and nitric oxide-dependent presynaptic enhancement in persistent forms of long-term potentiation. Learn Mem 18, 625–633, doi:18/10/625 [pii] 10.1101/lm.2245911 (2011).

90 Arancio, O. et al. Nitric oxide acts directly in the presynaptic neuron to produce long-term potentiation in cultured hippocampal neurons. Cell 87, 1025–1035, doi:S0092-8674(00)81797-3 [pii] (1996).

91 Zhuo, M., Small, S. A., Kandel, E. R. & Hawkins, R. D. Nitric oxide and carbon monoxide produce activity-dependent long-term synaptic enhancement in hippocampus. Science 260, 1946–1950 (1993).

92 Hopper, R. A. & Garthwaite, J. Tonic and phasic nitric oxide signals in hippocampal longterm potentiation. J Neurosci 26, 11513–11521, doi:10.1523/JNEUROSCI.2259-06.2006 (2006).

93 Akaike, T. et al. Antagonistic action of imidazolineoxyl N-oxides against endothelium-derived relaxing factor/.NO through a radical reaction. Biochemistry 32, 827–832 (1993).

94 Son, H. et al. The specific role of cGMP in hippocampal LTP. Learn Mem 5, 231–245 (1998).

95 Arancio, O. et al. Presynaptic role of cGMP-dependent protein kinase during long-lasting potentiation. J Neurosci 21, 143–149, doi:21/1/143 [pii] (2001).

96 Zabel, U. et al. Calcium-dependent membrane association sensitizes soluble guanylyl cyclase to nitric oxide. Nat Cell Biol 4, 307–311, doi:10.1038/ncb775 ncb775 [pii] (2002).

97 Nabavi, S. et al. Metabotropic NMDA receptor function is required for NMDA receptor-dependent long-term depression. Proc Natl Acad Sci U S A 110, 4027–4032, doi:1219454110 [pii] 10.1073/pnas.1219454110 (2013).

98 Sjostrom, P. J., Turrigiano, G. G. & Nelson, S. B. Multiple forms of long-term plasticity at unitary neocortical layer 5 synapses. Neuropharmacology 52, 176–184, doi:S0028-3908(06)00231-0 [pii] 10.1016/j.neuropharm.2006.07.021 (2007).

99 Dobrunz, L. E. & Stevens, C. F. Heterogeneity of release probability, facilitation, and depletion at central synapses. Neuron 18, 995–1008, doi:S0896-6273(00)80338-4 [pii] (1997).

100 Schultz, W. & Dickinson, A. Neuronal coding of prediction errors. Annu Rev Neurosci 23, 473–500, doi:10.1146/annurev.neuro.23.1.473 (2000).

101 Niv, Y. & Schoenbaum, G. Dialogues on prediction errors. Trends in cognitive sciences 12, 265–272, doi:10.1016/j.tics.2008.03.006 (2008).

102 Kim, J. & Alger, B. E. Random response fluctuations lead to spurious paired-pulse facilitation. J Neurosci 21, 9608–9618, doi:21/24/9608 [pii] (2001).

103 Schulz, P. E., Cook, E. P. & Johnston, D. Changes in paired-pulse facilitation suggest presynaptic involvement in long-term potentiation. J Neurosci 14, 5325–5337 (1994).

104 Faber, D. S. & Korn, H. Applicability of the coefficient of variation method for analyzing synaptic plasticity. Biophys J 60, 1288–1294, doi:S0006-3495(91)82162-2 [pii] 10.1016/S0006-3495(91)82162-2 (1991).

105 Ishikawa, D. et al. Fluorescent pipettes for optically targeted patch-clamp recordings. Neural Netw 23, 669–672, doi:S0893-6080(10)00046-8 [pii] 10.1016/j.neunet.2010.02.004 (2010).

106 Bettache, N., Carter, T., Corrie, J. E., Ogden, D. & Trentham, D. R. Photolabile donors of nitric oxide: ruthenium nitrosyl chlorides as caged nitric oxide. Methods Enzymol 268, 266–281 (1996).

107 Siegel, S. Nonparametric statistics for the behavioral sciences. (1956).

